# Bayesian bacterial GWAS model reveals threshold-dependent genetic changes in antimicrobial resistance phenotypes

**DOI:** 10.64898/2026.09.24.754083

**Authors:** Jackie Toussaint, Joel Hellewell, Tommi Mäklin, Samuel T. Horsfield, John A. Lees

**Affiliations:** European Bioinformatics Institute (EMBL-EBI), Hinxton, United Kingdom; University of Cambridge, Cambridge, United Kingdom; Jean Golding Institute, University of Bristol, Bristol, United Kingdom; Wellcome Sanger Institute, Hinxton, United Kingdom

**Keywords:** bayesian modeling, genome-wide association studies, fine mapping, microbial genomics, antimicrobial resistance

## Abstract

Genome-wide association studies (GWASs) utilize the association between genetic variants and phenotypic changes to identify causal variants underlying a phenotype of interest, and in some cases employ these correlations to perform phenotype prediction for novel genomes given their repertoire of genetic variants. Often utilizing a logistic or linear regression framework, bacterial GWAS methods have frequently been applied to infer variants associated with antimicrobial resistance (AMR) phenotypes measured using serial dilutions and recorded as minimum inhibitory concentrations (MICs). Beyond their increased granularity compared to resistant-sensitive classifications, MICs provide a more robust measure of AMR because they are not dependent on threshold values that can change over time and between data sources. One prominent challenge posed by both existing inference and prediction GWAS frameworks is the representation of MICs as censored ordered intervals rather than as binary resistant-sensitive or continuous approximations, a more suitable statistical model that has the potential to improve the discovery of lower-penetrance variants and MIC prediction. We show that Bayesian ordered logistic regression GWAS models can recover known antimicrobial resistance variants, identify variants with threshold-dependent effects without employing computationally-costly pairwise comparisons, and predict categorical MIC phenotypes. Specifically, the models recovered known resistance-conferring variants for *β*-lactams (in the penicillin-binding protein genes *pbp2x, pbp1a*, and *pbp2b*) and trimethoprim (in the dihydrofolate reductase gene *folA*) in *S. pneumoniae*, and for rifampicin (in the *rpoB* gene encoding the beta subunit of bacterial RNA polymerase) for *M. tuberculosis*, in addition to numerous lower-penetrance resistance variants. We found that ordered categorical models that allow for non-proportionality in the effect of each genetic variant across different MIC categories replicate the pattern of stepwise accumulation of resistance mutations in *pbp* genes in their relative effects at different MIC thresholds, and can improve balanced accuracy for MIC prediction compared to a proportional-effects model, suggesting that nonlinear effects may play a role in missing heritability. We also describe the use of lineage clustering as an alternative method of reducing false positives induced by genetic relatedness within a bacterial population. The methods described here incorporate a more accurate representation of minimum inhibitory concentration phenotypes into a highly flexible bacterial GWAS methodology capable of capturing nonlinear variant effects and suitable for both causal variant inference and prediction of MIC phenotypes.

## Introduction

Antimicrobial resistance poses amajor global public healththreat; in 2019, an estimated 4.95M AMR-associated deaths occurred, of which 1.27M were attributable to AMR (Antimicrobial Resistance Collaborators 2022). Genome-wide association studies (GWAS), which test for associations between genetic changes and phenotypic changes, have facilitated the identification of resistance-conferring genetic variants with great success (Lees, Mai, et al. 2020; Power et al. 2017; Kim et al. 2022). Interest in prediction of AMR from sequencing data has also grown in recent years, largely due to its potential utility in developing more rapid clinical diagnostics than are possible with laboratory-based phenotyping (Kim et al. 2022). Despite this interest, to our knowledge there have been no new bacterial GWAS association models developed since 2018 (Lees, Galardini, et al. 2018; Collins and Didelot 2018).

One GWAS methodology commonly applied to MIC data is a linear mixed model framework in which log_2_-transformed MICs are regressed on genetic variants with some measure of relatedness, such as a genetic-relatedness matrix (GRM) or principal components (PCs), included respectively as random or fixed effects, to reduce spurious correlations arising from population structure. A popular bacterial GWAS software implementing these methods is Pyseer (Lees, Galardini, et al. 2018). In addition to a multiple linear regression method intended for use with fixed-effect covariates for population structure control, such as PCs from pairwise distances, Pyseer offers a Fast-LMM option to quickly fit continuous traits using a GRM (calculated from a phylogeny or from genotype distances) as a random effect, with the option to add fixed-effect covariates such as known resistance variants to perform conditional analyses (Lees, Galardini, et al. 2018). The Fast-LMM method also implements an optional elastic-net mode to fit variants jointly with penalization, and produces an estimate of narrow-sense heritability (h^2^). Unitigs and k-mers, SNPs and indels, gene presence-absence, and burden-test genotypes are all supported (Lees, Galardini, et al. 2018).

Recently, Kulkarni et al. have described the use of multivariable logistic regression with L2 penalization as a promising catalogue-free alternative to the SOLO grading method based on the World Health Organization mutation catalogue for prediction of resistance in *M. tuberculosis*. The SOLO method identifies associations between each individual mutation and a binary resistance phenotype in a univariate logistic regression, then grades each mutation into five categories describing its likelihood to confer resistance based on confidence grading rules (which can include independent evidence from the literature); the resulting grades are used for phenotype prediction. The multivariate regression method was found to increase sensitivity and recover known resistance variants, including some compensatory mutations (Kulkarni et al. 2025). In addition to logistic regression, an L2-penalized linear regression was performed against log_2_-transformed MICs from largely non-overlapping data to validate the logistic model. 36% of variants associated with resistance in the logistic regression and found to be “Uncertain” using SOLO were found to be significant using linear regression with MICs, all of which reproduced effect direction; of those not found to be significant in the MIC model, 52% were found in fewer than 5 isolates in the MIC dataset. Importantly, analysis of hyper-susceptibility signals found not to be significant in the linear regression model indicated that linear regression displayed reduced susceptibility to sampling bias and variant multicollinearity compared to the logistic regression model. The authors note that logistic regression relies on the assumption that genetic effects are additive and well-modeled by a logistic function, which may not be true for all genetic relationships, and that effect sizes from linear regression models can also be interpreted as fold-changes on the MIC scale, which provides a more readily-interpretable metric of resistance than logistic regression odds ratios (Kulkarni et al. 2025).

Left- and right-censoring of MICs presents a potential challenge for linear regression models (Batisti Biffignandi et al. 2024); for example, right-censoring of the highest MIC breakpoint in the tested range can result in a sample’s true MIC value being several orders of magnitude greater than the break point itself, making fitting to breakpoints challenging. One proposed solution is to reframe the GWAS as a classification task with MIC categories. Batisti Biffignandi etal. recently evaluated the effect on model power of modeling MICs as a regression with log-scaled continuous data compared to a classification task with categorical data (expressing the phenotype as unordered MIC bins); pyseer’s FaST-LMM was benchmarked alongside elastic-net and random-forest models using phenotypes simulated with different underlying architecture as well as real MICs for four antibiotics in a large *K. pneumoniae* dataset. Batisti Biffignandi et al. found that model power is generally improved by encoding MICs as categorical when only four to seven dilution steps have been recorded and using log-scale linear regression for MIC datasets with greater resolution. However, while the change in power using unordered classification was explored, no ordered model was implemented, though this was mentioned as a possible future extension of their work (Batisti Biffignandi et al. 2024). Because ordered logistic models incorporate order as well as account for the ambiguity in breakpoints, we expected an ordered logistic method to increase model performance.

Conditioning on known resistance variants has also been found to improve identification of causal variants with smaller effects in linear regression against continuous MIC phenotypes. KC Ma et al. found that a unitig-based linear regression using log_2_-transformed azithromycin MICs in *N. gonorrhoeae* with known resistance mutations as fixed covariates identified an additional causal variant in *RplD* and displayed fewer false-positives (KC Ma et al. 2020). The linear regression model was also found to outperform a logistic regression with MICs binned into a binary resistant-sensitive phenotype (KC Ma et al. 2020). Multivariable regression has also previously been used in logistic models to control for the presence of known ARGs to better understand the effect of an ARG of interest (Lipworth et al. 2024). We aimed to develop a flexible model that could be easily expanded to use fixed covariates for this purpose.

Finally, even synonymous mutations within antibiotic-resistance genes (ARGs) have been found to result in decreased levels of resistance compared to the known reference gene, such as in the case of the *bla*_*TEM-1*_ gene conferring resistance to piperacillin–tazobactam and amoxicillin–clavulanic acid resistance in *E. coli* (Lipworth et al. 2024). We therefore sought to implement a model structure that, while allowing for the use of gene presence-absence, was also fast enough to be used with large numbers of individual variants, such as those arising from SNP- and unitig-based measures of variation.

In this paper, we introduce an ordered logistic Bayesian framework for bacterial GWAS that is uniquely suited to modeling phenotypes resulting from censored discrete observations of unobserved values on a continuous scale, such as MICs, with the aim of providing a fast, interpretable and flexible method to identify resistance-conferring variants, including those with threshold-dependent effects, and predict ordered categorical phenotypes.

## Methods

### Processing minimum inhibitory concentration phenotypes

Benzylpenicillin resistance and trimethoprim resistance were evaluated in *Streptococcus pneumoniae*, and rifampicin resistance was evaluated in *Mycobacterium tuberculosis*. All phenotypes used were originally reported as minimum inhibitory concentrations, which were used to generate binary, ordered categorical, and continuous phenotypes. For *S. pneumoniae* isolates, binary resistant-sensitive phenotypes were generated by applying the appropriate 2024 EUCAST breakpoint (0.06 µg mL^−1^ for penicillin, and S ≤ 1 and R ≥ 2 µg mL^−1^ for trimethoprim) to sample MICs (S3, S4); for *M. tuberculosis*, the commonly-used rifampicin breakpoint 0.125 µg mL^−1^ was applied since no EUCAST breakpoint has been made available for rifampicin resistance in *M. tuberculosis* (S5). Continuous phenotypes were defined as the log_2_-transformed MIC.

The precise breakpoints and thus number of ordinal categories *K* were determined by binning recorded MICs into larger intervals in accordance with the overall distribution of MICs in the dataset, such that extreme class imbalances and uninformative breaks (e.g. MIC ≤ 0.06 and MIC ≤ 0.065) were avoided. This was necessary as some MIC intervals contained very few samples. MICs were discretized using several automatic strategies: using all doubling dilutions containing ≥ 5% of samples or ≥ 10% of samples, or every other doubling dilution containing ≥ 5% of isolates (4-fold dilutions). For the *S. pneumoniae* penicillin resistance dataset, an additional dataset was created by placing breakpoints by hand at visual minima in the MIC distribution (the “dilution minima” method, see S3), analagous to an epidemiological cut-off (ECOFF). Once the breakpoints were determined, MICs were modeled as ordered categories expressed as integers from 1 to *K*.

Breakpoints for each of the methods and datasets are shown in S3, S4, S5.

### Train:test splits to evaluate model fit and predictive performance

To evaluate predictive performance across the different phenotypes, each prediction dataset was partitioned into training and test samples using two schemes (S6, S7). Firstly, the isolates were randomly assigned to an 80:20 train: test split. The same seed was used across datasets to allow for as similar a comparison of prediction metrics as possible. Secondly, a leave-one-subcluster-out (LOSO) split was performed, in which the largest lineage subcluster containing <20% of isolates was held out as the test set; this threshold was chosen to select a subcluster with a sufficient number of isolates to allow for calculation of predictive performance metrics while preventing the selection of a subcluster containing an extremely large proportion of the isolates. Because bacterial populations exhibit strong population structure, a random split in which close relatives of the test isolates are present in the training set can inflate predictive performance if non-causal lineage-associated variants are captured. LOSO addresses this by withholding all isolates from one clonal background, providing a more conservative estimate of predictive performance and general is ability to newly sequenced isolates.

To evaluate model fit on inference models, 20% of isolates (up to a maximum of 500) on which the model had been trained were randomly selected and their phenotypes predicted from the posterior likelihood. The same predictive performance metrics were applied. For ordered categorical models, an additional posterior predictive check evaluating the difference in the observed versus predicted category frequency was performed for the marginal MIC distribution (S29).

### Developing Bayesian models for different use cases

All models were written in Stan and are available for download from GitHub (see ‘Data and Code Access’). Models were fit using CmdStan (v2.37.0) with cmdstanr (v0.8.1) using the variational inference (VI) algorithm with seed 987654321. Four models were created, each with an inference and a prediction version: a logistic regression model for binary phenotypes, a proportional-odds ordered categorical model (POM) and a partial-proportional-odds ordered categorical model (PPOM) for binned MIC phenotypes, and a linear regression model for continuous (log_2_-transformed MIC) phenotypes ST2.

### Genotype standardization

Standardization of the binary genotype presence-absence matrix places all variants on a common scale so that the regularized horseshoe prior applies comparable shrinkage across variants of differing minor allele frequency (MAF), which has been previously recommended for multilevel logistic regression models (Gelman 2008). This parameterization ensures that a rare or recently-emerged variant is not penalized by MAF-dependent shrinkage, which is necessary when fitting jointly to all variants.

Prior to model fitting, each variant column *x*_*v*_ ∈ ℝ ^*N*^was centered and scaled to unit variance:

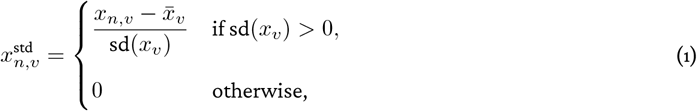

where 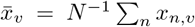 is the sample standard deviation. Though minor allele frequency filtering was applied in all datasets used, the models pin invariant columns (sd(*x*_*v*_) = 0) to zero to avoid division by zero. Standardization was performed with division by one standard deviation instead of two as all model predictors were binary (Gelman 2008).

Posterior effect sizes were backtransformed to unstandardized effects 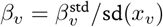 (for PPOM, unstandardized per-cutpoint effects 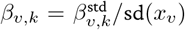 and per-allele cumulative odds ratios exp(*β*_*v*_) (for PPOM, exp(*β*_*v,k*_)), which were used for all results analyses.

### Regularized horseshoe prior for variant effect shrinkage

To perform variant selection across *V* highly-correlated candidate variants while allowing for some large effects, we placed a regularized horseshoe prior as described in Piironen and Vehtari (2017) on the standardized variant coefficients 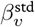. The horseshoe is a continuous analogue of the spike-and-slab prior, in which a component concentrated at zero (the “spike”) shrinks negligible coefficients toward zero while a diffuse component (the “slab”) allows large effects to escape shrinkage; the regularized form of Piironen and Vehtari (2017) replaces the horseshoe’s unbounded tail with a slab of finite variance 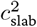, so that coefficients large enough to escape shrinkage have a prior of 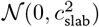 rather than an unbounded Cauchy tail. Writing 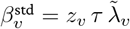, with

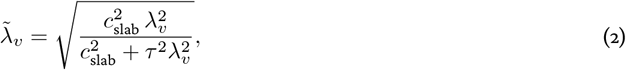

we assigned the priors

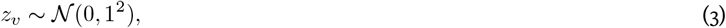

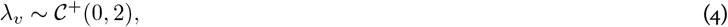

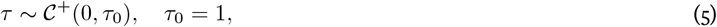

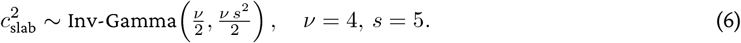

The local scales *λ*_*v*_ permit a small number of variants to escape shrinkage, while the global scale *τ* controls overall sparsity. This reflects our prior expectation that truly causal AMR variants are sparse, with a few target-site substitutions that shift MIC by many dilutions while most genome-wide variation has negligible effect. The slab variance 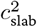 bounds the effective prior variance of non-zero coefficients to approximately 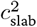, preventing the heavy tails of the standard horseshoe from inducing implausibly large effects and improving posterior geometry. The slab cap is deliberately generous because we expect that a single resistance substitution can raise MIC by orders of magnitude, unlike the small per-variant effects more typical of human complex traits.

Pooling a single set of shrinkage parameters per variant across all *K* − 1 cutpoints, as in the POM, logistic, and linear models, introduced a shrinkage penalty (due to the proportionality constraint) on the non-proportional variant effects that the PPOM is designed to recover. We therefore developed an independent regularized horseshoe prior on each (variant, cutpoint) coefficient 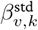, for *v* = 1,…, *V* and *k* = 1,…, *K* − 1. Writing 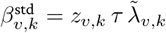, with

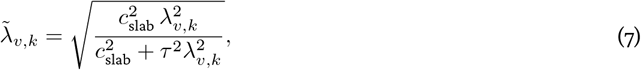

and with the new priors

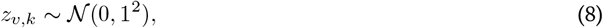

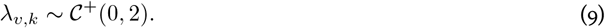

This parameterization estimates local scales *λ*_*v,k*_ independently per (variant, cutpoint), allowing a variant’s effect to be shrunk toward zero at some MIC thresholds while remaining non-zero at others. This ensures that the capture of, for example, a compensatory or second-step mutation with no measurable effect on a susceptible background that contributes only to high-level resistance will not be inhibited. The global scale *τ* and slab variance 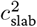 are shared across all *V* (*K* −1) coefficients, providing joint control of sparsity and tail behavior across the full effect matrix.

For all cases, *τ* _0_ was fixed rather than calculated using a variant of Equation 3.12 from Piironen and Vehtari (2017) for two reasons: firstly, because it is difficult to determine a plausible number of causal variants *a priori* given variation in linkage disequilibrium structure across species and in the underlying genetic architecture for different AMR phenotypes, and secondly, because a solution has not been made readily available for the ordered logistic case.

### Fast and effective population structure correction using lineage clusters and subclusters

Sub-populations within an overall population often have different baseline phenotype values, which can lead to variants being incorrectly identified as causal if only the relationship between individual sample genotypes and phenotypes is considered (Earle et al. 2016). For example, any variants commonly observed in a lineage exhibiting high AMR will display a stronger level of association with resistance, even if they are not responsible for the resistance phenotype (Earle et al. 2016; Hoffman 2013).

These lineage effects are most commonly addressed in human and bacterial GWAS though the inclusion of a genetic relatedness or “kinship” matrix (GRM) introduced as a random effect within a linear mixed model framework (Hoffman 2013). GRMs denote the proportion of shared variants between each pair of samples and thus scale quadratically with sample size. In contrast, assigning each sample to a lineage cluster and subcluster based on genetic distances scales linearly while replicating the population structure captured by GRMs (Lees, Harris, et al. 2019). The ability to input lineage clusters from any software also allows users to tailor the model’s population structure correction to the needs of their dataset, offering flexibility and possible computational savings.

To reduce spurious correlations resulting from population structure, we implemented a hierarchical lineage subclustering approach that fits each lineage cluster effect under a weakly informative Gaussian prior and partially pools each sub-cluster effect toward its “parent” lineage cluster via a non-centered parameterization with a shared scale; the resulting sub-cluster effects are then re-centered to sum to zero within each parent lineage.

In this implementation, *S* lineage subclusters are nested within *L* lineage clusters *π*(*s*) ≤ {1,…, *L*]. Lineage effects 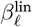 are drawn from a weakly informative prior,

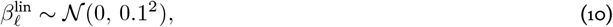

and lineage subcluster effects are partially pooled toward their parent lineage via a non-centered parameterization,

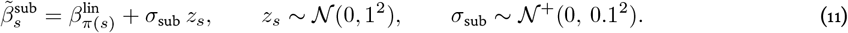

To make these effects identifiable alongside the model’s location anchor (intercept *α* for the continuous and logistic models, cutpoint *c*_1_ for the PO and PPO models), a treatment-contrastencoding was applied. The lineage subcluster with the minimum mean phenotype value (most susceptible) is dropped from the design matrix **X**_sub_ ≤ R^*N* ×(*S*−1)^, so that all remaining lineage subcluster effects are interpreted as deviations from that susceptible reference.

We further impose a within-lineage sum-to-zero constraint by re-centering

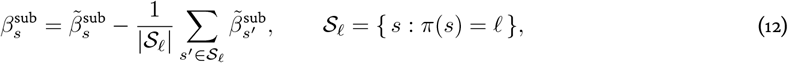

so that lineage subcluster effects represent within-lineage deviations while lineage effects absorb the lineage-level mean. The lineage contribution to the linear predictor for sample *n* is then 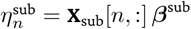, and is constrained to be proportional across cutpoints in both the POM and PPOM. For simplicity, we write **s**_*n*_ = **X**_sub_[*n*, :] for the *n*-th row of the subcluster design matrix and ***ω*** = ***β***^sub^ for the corresponding (re-centered) subcluster coefficient vector in the likelihoods Equation 22 and Equation 23.

In prediction models, test samples assigned to lineage subclusters not present in the training dataset inherit a subcluster effect sampled only from its prior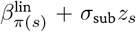, which shrinks toward the parent lineage effect. If the parent lineage is also unobserved during training, the parent lineage effect is similarly defined as its *N* (0, 0.1^2^) prior.

To examine the effectiveness of the lineage subcluster-based population structure correction, we also implemented a GRM-based version of the logistic model, in which the subcluster term **s**_*n*_***ω*** of Equation 25 is replaced by a per-isolate random effect *u*_*n*_. The haploid VanRaden GRM **K** (VanRaden 2008) computed from the allele-frequency-standardized genotype matrix (see “Datasets”) was incorporated in a non-centered parameterization,

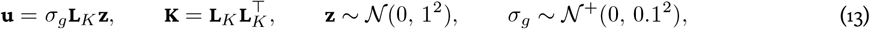

where **L**_*K*_ is the lower Cholesky factor of **K**, and the prior on *σ*_*g*_ was chosen to match the scale of *σ*_sub_ (Equation 11) that it replaces. **K** was scaled to unit mean diagonal so that *σ*_*g*_ is on the same scale as *σ*_sub_, and givena 10^*-*6^ diagonal jitter since centering the genotype columns renders **K** singular.

### Lineage subcluster-based population structure correction performs comparably to GRM-based method

Linear mixed models incorporating a genetic relatedness matrix as a random effect are the standard approach to population structure control in human and bacterial GWAS (Hoffman 2013; Earle etal. 2016). A recent benchmark of bacterial association methods found that an LMM with a phylogeny-derived relatedness matrix as a random effect provided the best control of false positives compared to the use of multidimensional scaling (MDS) components as fixed covariates, an elastic net, and a LASSO regression. However, power was lowest with this method, and in highly clonal populations, high false-positive rates have been reported in highly clonal populations where the kinship matrix is not carefully chosen (Lees, Mai, et al. 2020).

Using *S. pneumoniae* with the binary penicillin resistance phenotype, we compared the logistic model that uses lineage subcluster-based population structure correction to the same model using a GRM-based method, as described in Methods. Despite being faster to fit than a GRM (0.0138s per gradient evaluation versus 0.0151s per gradient evaluation, a 9% performance improvement), the model with subclusters identified the same loci with similar null distributions of effect sizes and RATEs (S15). The GRM scales quadratically with N, so the performance improvement of the linearly-scaling subclusters method is expected to grow as N increases. When we attempted to perform this analysis using the 11,622-isolate *M. tuberculosis* dataset, the GRM created was 2.7 GB, nearly twice the 1.43 GB required by the entire LD-pruned genotype matrix, which rendered streaming into Stan models using `cmdstanr::write_stan_json`() impossible due to its 2^31^ 1 byte limit. (Incontrast, the lineage subcluster method for this dataset added only *L* − 1= 7 lineage cluster columns and *S* 1 = 78 lineage subcluster columns to the design matrix.)

Both models identified variants in *pbp2x* and *pbp2b* as the only loci in which there was a variant with an 89% posterior credible interval excluding zero, and both assigned their highest RATE value to a *pbp2b* variant. The same missense variant in *pbp2x* was called by both, but the models identified a different SNP within *pbp2b* (at position 1,613,379 under the hierarchical model versus 1,613,302 using the relatedness matrix, 77 bp apart). The relatedness model additionally called a missense variant (Val27Ile) in *SPN23F03030*, an uncharacterised membrane protein whose coding sequence lies 1.9 kb upstream of *pbp2x*; this variant is 2.45 kb from the shared *pbp2x* variant, and possibly results from linkage disequilibrium with this locus rather than representing an independent signal, since the hierarchical model assigned it a negligible effect 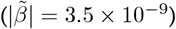.

### Intercept priors for the logistic and continuous models

The continuous and logistic model intercept *α* is anchored on a data-informed empirical baseline computed from the reference lineage subcluster:

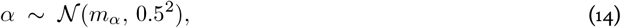

where the prior standard deviation of 0.5 corresponds to approximately one log_2_-dilution. For the continuous model, the baseline is the empirical mean log_2_-MIC of the reference subcluster:

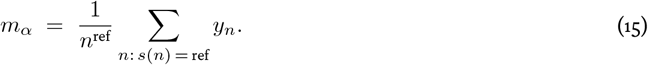

For the logistic model, the baseline is computed from the susceptible fraction of the reference subcluster:

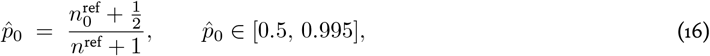

where 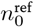 is the number of susceptible isolates in the reference subcluster. The reference subcluster being the lowest-mean-phenotype subcluster guarantees 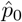 is high. The intercept anchor is then

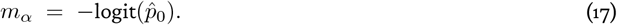

### Cutpoint priors for ordered categorical models

Rather thanfitting anintercept withcutpointsfixedatthe known MICbreakpoints, *K* −1 cutpoints **c** = (*c*_1_,…, *c*_*K*−1_) were estimated under a strict ordering constraint *c*_1_ < *c*_2_ < … < *c*_*K*−1_, with each cutpoint given a wide normal prior centered on a latent-scale anchor *m*_*c,k*_ derived from the corresponding MIC breakpoint:

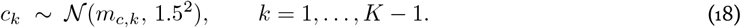

The prior standard deviation of 1.5 is intentionally wide on the logistic-latent scale to prevent misspecification: because the latent variable is the underlying continuous concentration at which growth is inhibited, of which a recorded MIC is a censored and discretised observation, the cutpoints represent the dilution steps at which the assay partitions that continuum, which have been previously found to be reproducible only to within roughly one doubling dilution.

The anchors *m*_*c,k*_ are constructed from two components: the log_2_-MIC breakpoint grid ***K*** = log_2_(mic_breakpoints), which provides the spacing between cutpoints under the assumption that one log_2_-dilution step corresponds to one unit on the logistic latent scale, and a level shift derived from the reference subcluster that aligns the first cutpoint *c*_1_ with the logit of its empirical category-1 frequency, so that the implied *P* (*Y* = 1) matches that frequency. The Laplace-smoothed fraction of reference-subcluster samples in MIC category 1 is

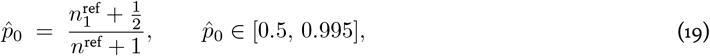

with the same clamp justification as in the logistic model (Anscombe 1956; Gart and Zweifel 1967). Note that 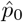 in the logistic model (Equation 16) is built from the count of phenotype category 0 (susceptible isolates,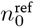), but in the ordinal models is built from the count in MIC category 1 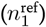. The level shift is

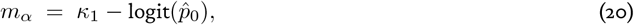

and each cutpoint anchor is the corresponding log_2_-MIC breakpoint shifted by this constant,

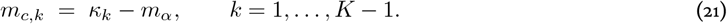

This shifts the MIC grid so that *c*_1_ sits near logit 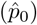 while preserving the relative spacing of the MIC breakpoints. The per-cutpoint drift *c*_*k*_ − *m*_*c,k*_ can then be used to identify categories where the MIC anchor may be misspecified or uninformative (S1, S28).

### Summary with likelihood equations

The proportional-odds ordered categorical model (POM) pools the estimate of each variant’s coefficient ***β*** across all cutpoints:

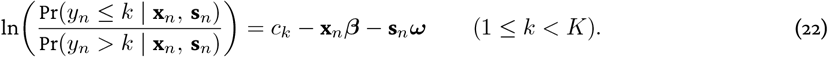

For the partial-proportional-odds model (PPOM), the proportional-odds assumption is relaxed for variant effects only by indexing ***β*** by cutpoint, while the lineage subcluster effects ***ω*** remain fixed across cutpoints (S2). The PPOM likelihood is implemented as *K* −1 Bernoulli terms per sample:

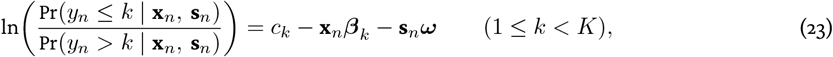

where *k* is an ordered response category, **x**_*n*_ is a vector of variants for sample *n*, ***β***_k_ is a vector of logit coefficients that are unconstrained between cutpoints, **s**_*n*_ is a vector of lineage subclusters, ***ω*** is a vector of subcluster logit coefficients that are fixed across cutpoints, and *c*_*k*_ is a cutpoint (Fullerton and J Xu 2012).

The continuous model log_2_-MIC outcome was modeled as:

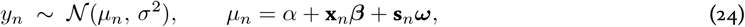

with a half-normal prior ~ *N* ^+^(0, 1^2^) centered on one log_2_-dilution.

For the logistic model, the same linear predictor was used with the logit link function:

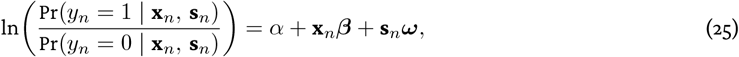

with *y*_*n*_ ≤ {0, 1} being the sensitive-resistant phenotype.

### Estimating heritability

After model fitting, narrow- and broad-sense heritability were estimated from the additive genetic (variant effects) component, the population-structure (lineage cluster and subcluster effects) component, and the residuals. Here **x**_*n*_ = **X**^std^[*n*, :] is the *n*-th row of the standardized genotype matrix and **s**_*n*_ = **X**_sub_[*n*, :] the *n*-th row of the subcluster design matrix, consistent with the likelihoods above:

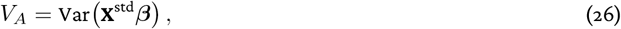

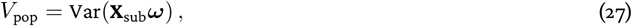

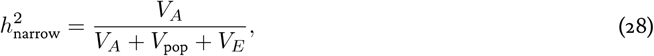

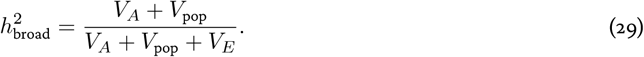

The residual variance was defined as *V*_*E*_ = *σ*^2^ for the continuous model, and *V*_*E*_ = *π*^2^/3 on the logistic-latent (liability) scale for the logistic, POM, and PPOM models. Because the PPOM generates per-variant effects, *V*_*A*_ and *h*^2^ were computed for each cutpoint *k* using ***β***_k_. Because the regularized horseshoe shrinks ***β*** toward zero, 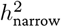 should be interpreted as a lower bound. For brevity, we write 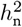 and 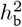 for narrow- and broad-sense heritability in the remainder of the text.

### Phenotype prediction

Phenotype prediction was performed by computing the linear predictor from posterior draws of the variant and subcluster coefficients, then sampling from the corresponding outcome distribution. For the continuous model, *ŷ*_*n*_ was drawnfrom *N* (*µ*_*n*_, *π*^2^) using `normal_rng`; for the logisticmodel, *ŷ*_*n*_ ≤ {0, 1} was drawn using `bernoulli_logit_rng`; and for the POM, *ŷ*_*n*_ ≤ {1,…, *K*} was drawn using `ordered_logistic_rng`. As Stan doesn’t directly support partial-proportional-odds in ordered categorical models, per-sample category probabilities were instead computed analytically from the cumulative cutpoint logits and returned as an *N*_test_ × *K* probability matrix.

All predictive performance metrics were computed in R (v4.4.3). Classification metrics for the binary and ordinal models (sensitivity, specificity, balanced accuracy (bACC), support-weighted positive predictive value (PPV), and F_1_) were calculated using yardstick (v1.4.0), the area under the ROC curve (AUROC) using pROC (v1.19.0.1), and confusion matrices with caret(v7.0.1). Ranked probability scores (RPS), ranked probability skill scores (RPSS), and the continuous ranked probability score (CRPS) were computed using scoringRules (v1.1.3). Brier score, RMSE, MAE, coefficient of determination (*R*^2^), and essential agreement were calculated in base R.

### Interpreting ordinal prediction metrics

To evaluate prediction accuracy for ordered categorical models, we report the categorical classification accuracy metrics balanced accuracy (bACC) and positive predictive value (PPV), and the ordinal categorical scoring rules ranked probability score (RPS) and ranked probability skill score (RPSS).

Because AMR datasets are often dominated by susceptible isolates, we first calculated bACC and PPV, which are computed over the MIC categories observed in the test set and are bounded on [0, 1]. bACC averages per-class sensitivity and specificity so that performance is not inflated by high-frequency MIC categories, while PPV is the support-weighted mean of per-class precision.

Since MIC calls can vary by roughly one dilution step between replicates, Batisti Biffignandi et al. and Eyre et al. recommend treating “off-by-one” classifications as correct during model evaluation with large numbers of classes. However, categorical measures treat every misclassification as equally incorrect. In contrast, RPS captures the degree of disagreement by penalising a misclassification in proportion to its distance from the true category, downweighting off-by-one errors without discarding them entirely (Batisti Biffignandi et al. 2024). We compute RPS per isolate from its predicted category probabilities, scaling it by 1/(*K* −1) so that it lies on [0, 1], where 0 denotes no error and 1 denotes the worst possible misclassification (the category farthest from ground truth), and report both its mean and its median across the test set (S12, S13, S14).

RPSS expresses the mean RPS as skill relative to a reference model, in which 1 indicates perfect prediction, 0 indicates performance at parity with the reference model, and negative values indicate a performance worse than the reference model. We calculated RPSS against two baselines: a model that assigns equal probability to each MIC category (RPSS_unif_), and a model that assigns probability to each class in alignment with their true frequencies in the dataset (RPSS_freq_) (ST8).

To compare how well the ordered logistic POM and PPOM were able to capture variant effects relative to a binary or continuous representation of the phenotype, phenotype prediction was performed on the same datasets using logistic and continuous prediction models (ST6, ST7). Because the difficulty of prediction increases as the probability of a correct random guess decreases, the resulting prediction metrics should be interpreted with respect to the granularity of the phenotype (binary, multi-category, or continuous).

The logistic model was scored using balanced accuracy, sensitivity and specificity (recall of resistantand of susceptible isolates), the area under the ROC curve (AUC; the probability that a randomly chosen resistant isolate is assigned a higher predicted resistance probability than a susceptible one, displaying how well the model separates resistant from susceptible isolates independently of the breakpoint threshold), the Brier score (the mean squared error of the predicted resistance probability for measurement of how well the probability is calibrated), the F_1_ score (the harmonic mean of precision and recall for the resistant class, summarising detection of resistance when resistant isolates are the minority), and very major and major error rates (VME, a resistant isolate predicted to be susceptible, and ME, the inverse).

The continuous model predicts log_2_ MIC directly, so its errors are reported on the doubling-dilution scale: the root-mean-squared error (RMSE) and mean absolute error (MAE) measure the deviation between predicted and observed log_2_ MIC in units of two-fold dilutions (such that lower is better), coefficient of determination *R*^2^ is the fraction of MIC variance explained (at most 1, and negative when the model predicts worse than the mean), and the continuous ranked probability score (CRPS) is the distributional analogue of MAE that scores the full posterior predictive rather than a point estimate. We also include a more interpretable metric for AMR GWAS with MIC phenotypes, essential agreement (EA), which is the fraction of isolates predicted within one doubling dilution of the true MIC.

### Evaluation of variant significance using relative centrality and cppRATE

As *p*-values in the classical frequentist meaning are not applicable to a Bayesian model, a different approach is required to measure variant significance in a Bayesian GWAS. RelATive cEntrality (RATE) values introduced in (Crawford et al. 2019) offer an alternative framework of Bayesian hypothesis testing that is based on comparing the relative importance of each obtained regression coefficient *β*_*v*_ against a “null hypothesis” where the effect of each *β*_*v*_ is assumed to be 0.

Briefly, (Crawford et al. 2019) base the RATE of a regression coefficient *β*_*v*_ on comparing the posterior distribution of the vector ***β***_*-*v_ containing all regression coefficients except the variant of interest *β*_*v*_, denoted as *p (β*_*-*v_), against the conditional posterior distribution of ***β***_*-*v_ on *β*_*v*_, denoted as *p* (***β***_*-*v_ |*β*_*v*_). This comparison is performed by computing the Kullback-Leibler divergence (KLD) between these two distributions, which we denote by KLD [*p (****β***_*-*v_) ∥ *p*(***β***_*-*v_ | *β*_*v*_)]. Importantly, the KLD is non-negative and takes the value of 0 if and only if the posterior distribution of ***β***_*-*v_ is independent of *β*_*v*_, implying support for the “null hypothesis” that *β*_*v*_ is not important relative to the other coefficients. Larger values in contrast indicate more important variables *β*_*v*_ and provide evidence against the “null hypothesis”. A closed form for computing KLD [*p*(***β***_*-*v_ ∥*p (****β****-*v |*β*_*v*_)} for each *v* ≤ {1,…, *V*} is given in (Crawford et al. 2019) assuming approximately multivariate normal distribution for the full vector ***β***.

Since the magnitude of the KLD can vary highly depending on the model under evaluation, determining a thresh-old for what constitutes a significant variant can be challenging. To get around this, (Crawford et al. 2019) define the

RATE of eachvariant*v* ≤ {1,…, *V*} by computing the individual values foreach *β*_*v*_, KLD [*β*_*v*_]= KLD [*p (****β***_*-*v_) ∥*p(* ***β***_*-*v_ | *β*_*v*_)], and using a transformation that scales the KLD of each variant *v* by the sum of the KLDs of all variants. This results in values of the form RATE 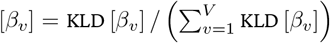, where RATE [*β*_*v*_] ≤ [0, 1] and 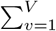 [*β*_*v*_]= 1. In this relativized form, a value of 1 implies a variant that fully determines the posterior distribution, while values close to 0 imply unimportant variants.

In practice, a convenient method of obtaining RATE values is through our C++ implementation of the RATE algorithm, called cppRATE (https://github.com/tmaklin/cpprate), which provides a more computationally efficient and parallelisable version of the original R code provided in (Crawford et al. 2019). Regardless of the implementation, the downside of using RATE values is that the memory consumption scales with the number of variants tested. We address this by using linkage disequilibrium pruning to reduce the number of variants that need to be tested (see ‘Linkage disequilibrium pruning with BacPrune-Rust’), making RATE computation feasible on the pruned set of variants.

For the PPOM, RATE values were computed separately at each cutpoint*k* using the corresponding coefficient vector ***β***_k_, so that 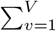 RATE [*β*_*v,k*_]= 1 within each cutpoint.

### Linkage disequilibrium pruning with BacPrune-Rust

Linkage disequilibrium (LD) refers to the non-random association of alleles at different loci and is often observed in regions of the genome with low recombination rates (Prunier etal. 2019). Perfect correlation between predictor variables, such as variants in perfect linkage, can cause a range of model fitting problems including global non identifiability. Even imperfectly correlated predictors induce a “grouping” or sharing effectanalogous to thatof the elastic-net, in which the signal at a locus is divided among its correlated variants such that no individual coefficient reaches the magnitude of the true effect (Lees, Mai, et al. 2020). For this reason, variants in strong linkage are often removed (“pruned”) prior to model fitting, leaving one representative, then subsequently “de-pruned” prior to downstream analysis such that each pruned variant is assigned the coefficient of its representative. The aim of LD pruning in GWAS is to improve model performance and minimize redundant information while retaining the relevant genetic variation across the genome (Prunier et al. 2019).

I developed and used the linkage disequilibrium (LD) pruning software BacPrune-Rust (v0.9.2), which allows for pruning of haploid genotype matrices using D’ or r scores with optional additional filtering by minor allele frequency (MAF). BacPrune-Rust is written in Rust and designed to scale to matrices with millions of variants and thousands of isolates, such as the complete CRyPTIC Consortium *M. tuberculosis* dataset used in this paper, which contained 1.7M variants and 11,622 isolates. It is available on bioconda and GitHub (https://github.com/bacpop/BacPrune-Rust/). Because two variants with identical presence and absence will inherently have the same effect size, we only performed pruning on variants in perfect correlation as measured by Pearson *r*^2^; this reduces the pathological geometry induced by strong correlations while obtaining accurate effect sizes for all variants.

### GHOST: Creating a workflow in R for ease of use

Models were fitted using GHOST(“Gwas with a Hierarchical& Ordered Stan Toolkit”, availableat https://github.com/bacpop/GHOST). GHOST runs different models according to user specification, generating all files required after each step and outputting the results into a standardized folder structure (Figure 2). In addition to the existing models, GHOST can be used with custom Stan models that take the same inputs using the `–model` flag. The pipeline can be run on an HPC cluster and includes an example Slurm submission script. GHOST facilitates easier reproduction of analyses and submission of datasets with multiple sets of pipeline options, as well as significantly reduces the amount of time the user must spend modifying inputs for the various programs.

Runtimes and memory consumption for LD pruning with BacPrune-Rust, Stan model fitting, and cppRATE using the GHOST workflow are shown in S8. For the 616-isolate *S. pneumoniae* penicillin dataset (32,406 variants, 32 CPUs), LD pruning completed in under 2 s, model fitting took 8–9 min for the logistic, POM and continuous models and 41 min for the PPOM, and cppRATE took 35–55 min. For the 11,622-isolate *M. tuberculosis* rifampicin dataset (75,272 variants, 48 CPUs), the same three stages took 3–6 min, 73 min–42.9 h and 3.5–5.5 h respectively; the PPOM is consistently the slowest to fit, as expected given that it estimates *V* (*K* −1) rather than *V* coefficients. Maximum memory was 63 GB for LD pruning, 55 GB for model fitting and 30 GB for cppRATE. LD pruning was therefore the most memory-intensive stage of the workflow despite being the fastest. BacPrune-Rust is currently single-threaded, so parallelising this computation could reduce the amount of memory per thread.

### Datasets

Each GWAS dataset we created comprises three major components: sample phenotypes, sample genotypes, and a representation of the population structure to control for spurious associations. We first processed the MIC phenotypes for each isolate to obtain a binary, ordered categorical, or continuous phenotype as described in “Processing minimum inhibitory concentration phenotypes”. We then produced a genotype matrix recording the presence or absence of each genetic variant in each isolate by calling SNPs and short indels against a reference genome, which was also used to annotate the variants. (Reference-free representations such as unitigs, *k*-mers, and gene presence–absence matrices could also be used as input.) Variants observed in too few isolates to support an effect estimate (those with very small minor allele frequency) were filtered out. Variants in perfect linkage disequilibrium are statistically indistinguishable from one another, so all but one representative were pruned before fitting and their coefficients assigned afterwards (see “Linkage disequilibrium pruning with BacPrune-Rust”). Finally, we used lineage clusters and subclusters to describe population structure. Because bacterial populations are clonal, any variant that happens to mark a lineage with elevated resistance will correlate with the phenotype whether or not it causes it. The lineage clusters and subclusters (and, for comparison, a genetic relatedness matrix) enter the model as covariates so that this shared ancestry is absorbed rather than attributed to individual variants (see “Fast and effective population structure correction using lineage clusters and subclusters”).

#### *Streptococcus pneumoniae* (benzylpenicillin, trimethoprim)

616 samples, 32,406 variants (fitted on 26,512 representative variants after pruning with BacPrune-Rust (v0.9.2) at *r*^2^ = 1). These data were obtained from the Massachusetts asymptomatic carriage collection sampled from children attending routine primary-care visits in Massachusetts, USA, across three cross-sectional nasopharyngeal colonisation surveys conducted in the spring of 2001 (shortly after the introduction of the seven-valent pneumococcal conjugate vaccine), the winter and spring of 2004, and the winter and spring of 2006–2007 (Croucher, Finkelstein, Pelton, Mitchell, et al. 2013; Croucher, Finkelstein, Pelton, Parkhill, et al. 2015). We obtained lineage clusters with PopPUNK (v2.7.2) by sketching each genome at *k* = 13, 17, 21, 25, 29 with a sketch size of 10^4^ and calling clusters from a refined network boundary (`refine` model) initialised from a two-componentmixture fit to the core/accessory distance distribution. Lineage subclusters were obtained with PopPIPE (v1.2.0) run on the PopPUNK database with `min_cluster_size = 3`, using SKA2 (v0.3.0) split *k*-mers (*k* = 31, frequency filter 0.9), within-strain trees from IQ-TREE (v2.3.5) under the 012310+G+ASC model, and two hierarchical levels of fastbaps (v1.0.8) clustering; the first fastbaps level was used as the subcluster label (Lees, Harris, et al. 2019). PopPUNK and PopPIPE identified 62 lineage clusters and 137 subclusters with a median of 2 isolates per subcluster. Variant calling from the draft assemblies was performed with Snippy (v4.6.0) in contig mode (`–ctgs`) against the ATCC 700669 reference genome (Spain^23F^ ST81; NCBI Reference Sequence: NC_011900.1) (Danecek et al. 2021). Per-sample VCFs were merged with bcftools (v1.21, `merge –missing-to-ref`), converted to haploid genotype calls (`+fixploidy - f 1`) and filtered to minor allele frequency (MAF) 0.05 (`filter -e ‘INFO/MAF<0.05’`). Where multiple minor alleles were present at a given chromosomal position, variants at that site were independently filtered by MAF before multiallelics were merged. Variants were annotated using SnpEff (v4.3.1) and SnpSift (v5.2) with the Streptococcus_pneumoniae_atcc_700669 SnpEff database (based on NCBI Reference Sequence: NC_011900.1) (Cingolani et al. 2012; Ruden et al. 2012). Loci containing only “modifier” alleles as defined by SnpEff (putative upstream and downstream effectors) were excluded. A haploid VanRaden genetic relatedness matrix was calculated from the allele-frequency–standardized VCF using R (v4.2.2). Where gene names were not available in the reference, eggNOG-mapper (v2.1.13) was used to annotate the reference proteins using DIAMOND (v2.0.15) in sensitive mode with iterated search (`–sensmode sensitive –dmnd_iterate yes`) to search the bacterial database bacteria.dmnd and the eggNOG orthology database (v5.0.2) for functional annotation (Buchfink et al. 2021; Huerta-Cepas et al. 2019; Cantalapiedra et al. 2021).

#### *Mycobacterium tuberculosis* (rifampicin)

11,622 samples, 75,272 variants (fitted on 40,941 representative variants after pruning with BacPrune-Rust (v0.9.2) at *r*^2^ = 1). These data were obtained from the Comprehensive Resistance Prediction for Tuberculosis: an International Consortium (CRyPTIC) compendium’s global collection of clinical isolates, which were sequenced and phenotyped as part of a large-scale international drug-resistance surveillance effort that spanned 27 countries and six continents and was deliberately enriched for drug-resistant isolates (Consortium 2022). “Re-genotyped” VCFs from individual isolates were obtained from the CRyPTIC Consortium (dataset version v2.1.2) (Consortium 2022), merged with bcftools (v1.21; `merge -m none`) in 20 equally sized intervals spanning NC_000962.3 (4,411,532 bp), converted to haploid calls (`+fixploidy -f 1`), tagged with allele frequencies (`+fill-tags -t MAF`) and filtered to MAF 0.0004 (which corresponds to a minimum presence in five samples) before the chunks were concatenated (Danecek et al. 2021). This was necessary due to the size of the data. Genotypes were binarised with `bcftools +setGT` (missing calls set to 0, any non-reference call set to 1) and the variant × sample matrix was transposed with GNU datamash (v1.9) due to its size. Variants were annotated using SnpEff (v4.3.1) and SnpSift (v5.2) using the Mycobacterium_tuberculosis_h37rv SnpEff database (based on the H37Rv reference genome; NCBI Reference Sequence: NC_000962.3) (Cingolani et al. 2012; Ruden et al. 2012). Where gene names were not available in the reference, eggNOG-mapper (v2.1.13) was used to annotate the reference proteins using DIAMOND (v2.0.15) in sensitive mode to search the bacterial database bacteria.dmnd and the eggNOG orthology database(v5.0.2) forfunctional annotation(Buchfink et al. 2021; Huerta-Cepas et al. 2019; Cantalapiedra et al. 2021). Lineage clustering and sub-clustering was performed with Fastlin (0.2.2) on the sequencing reads using the MTBC barcode set, with *k* = 25,a minimum of 4 occurrences per *k*-mer, a minimum of 3 barcodes per lineage call anda maximum *k*-mercoverage of 80× (Derelle et al. 2023). Samples receiving more than one lineage designation (indicative of mixed or contaminated samples) were excluded. Fastlin identified 8 lineages and 79 subclusters with a median of 23 isolates per subcluster, when using the first level of the barcode as the lineage cluster and the complete barcode as the subcluster.

## Results

### Ordinal logistic regression with partial proportional odds identifies known resistance loci with threshold-dependent effects

An ordered categorical model with proportional odds (POM) attempts to fit one coefficient per variant that describes the change in log odds of being in a category equal to or less than a given cutpoint; this coefficient is estimated jointly across cutpoints. Non-proportional (threshold-specific) variant effects can be accounted for by estimating their coefficients marginally at each cutpoint, such that each coefficient describes the change in log odds at that cutpoint (Figure 1). As the measure of population structure, lineage subclusters, remain constrained to have proportional effects across cutpoints, this implementation is categorized as a “partial proportional odds model” (PPOM).

**Figure 1.**
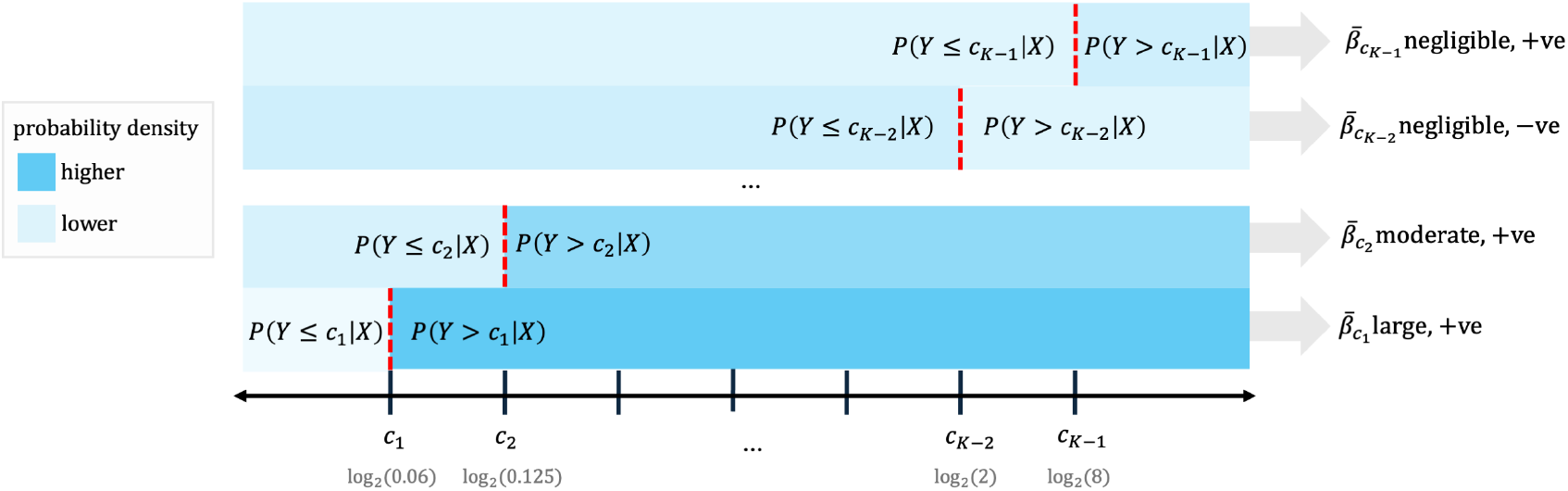
Example estimation of variant coefficients 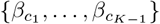 in a partial proportional odds (PPO) GWAS model with *K* ordered categories and one variant, which displays non-proportional effects. Priors for cutpoints {*c*_1_,…, *c*_*K*−1_] are centered around the categories’ log_2_-transformed minimum inhibitory concentration breakpoints, expressed here in µg mL^−1^. In a proportional odds (PO) model, the estimation of the coefficients are constrained to remain proportional across cutpoints, resulting in a single coefficient *β*_*c*_; to fit this coefficient, the large effects displayed by this variant at some cutpoints must necessarily be diluted. As the PPO model retains the effects observed at lower cutpoints and can optimize their fit, the signal for this example variant is likely to be markedly decreased in the PO model compared to the PPO model.

**Figure 2.**
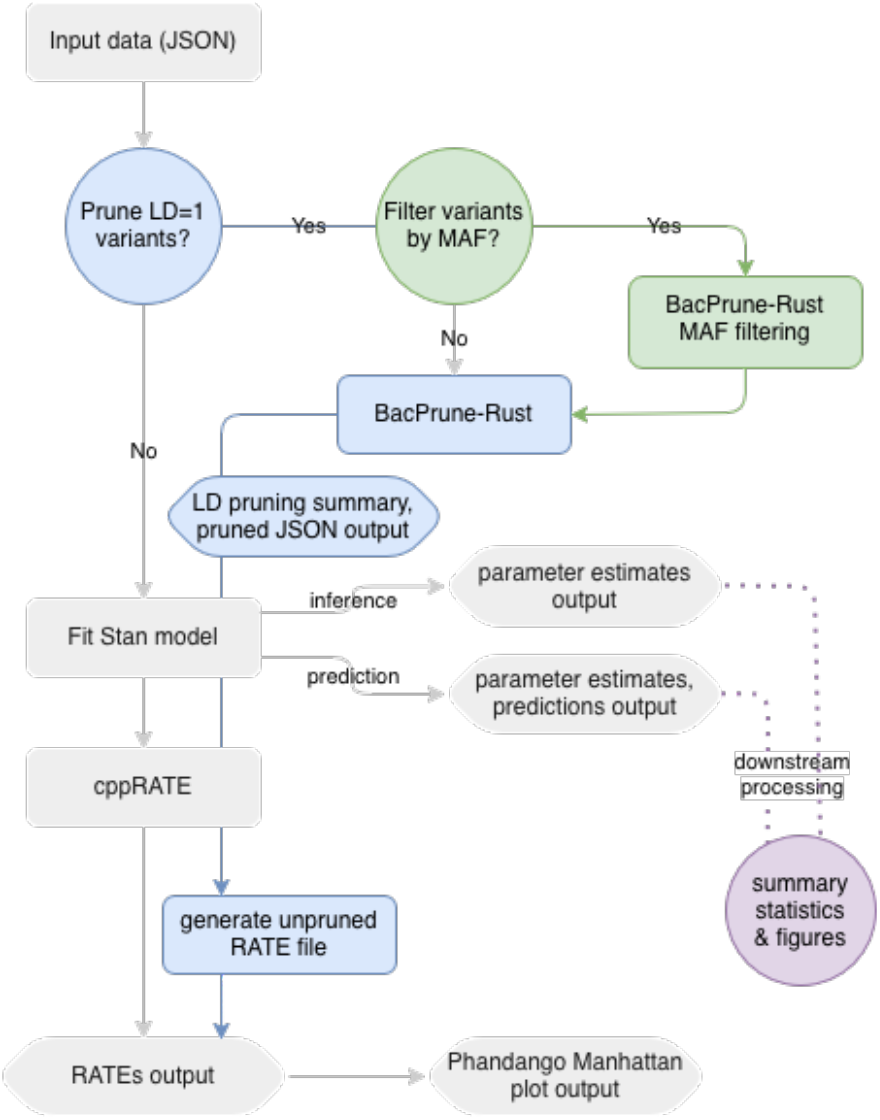
Overview of the GWAS workflow used. An option to fit models prior to linkage disequilibrium (LD) pruning was retained for testing purposes, but should not be used in practice as it may introduce parameter nonidentifiability.

We first applied the POM and PPOM to penicillin resistance in *S. pneumoniae* using the Massachusetts carriage collection with four MIC discretization strategies (see “Datasets”) to determine whether the ordered models recover knownresistance determinants andwhetherrelaxing the proportional-odds constraintcanidentify varianteffects that differ between MIC thresholds. Inapreviousanalysisof penicillin resistance in this *S. pneumoniae* dataset, Chewapreecha et al. found significant resistance-associated loci in the penicillin-binding protein (PBP) genes *pbp2x* (*pbpX*), *pbp2b* (*penA*), and *pbp1a*. Of the more than 300 SNPs reported, many nonsynonymous substitutions were previously implicated in reduced *β*-lactam binding (Chewapreecha et al. 2014). Heritability analyses confirmed that the core PBP resistance regionsaccount for approximately 50% of thevariationinpenicillin MICs across clinical isolates (Mallawaarachchi et al. 2022).

The PPOM recovered these loci with high effect sizes for all discretization strategies, as did the logistic model (Figure 3, S17, S18, S21, S22, S16). Small effectsizes of variants in *pbp1b* and *pbp2a*, whichare notimplicated in *β*-lactam resistance, are shown as negative controls ensuring that the model does not assign large effects to PBP loci indiscriminately (Figure 4, S9, S10). However, the POM with the most granular discretization strategy did not discover *pbp1a* prominently (S25). This aligns with findings from Batisti Biffignandi et al. that categorical models (tested as non-ordinal invariant expression models) most significantly improve performance over linear regression models when MICs are discretized into three to five bins as opposed to *>* 5.

**Figure 3.**
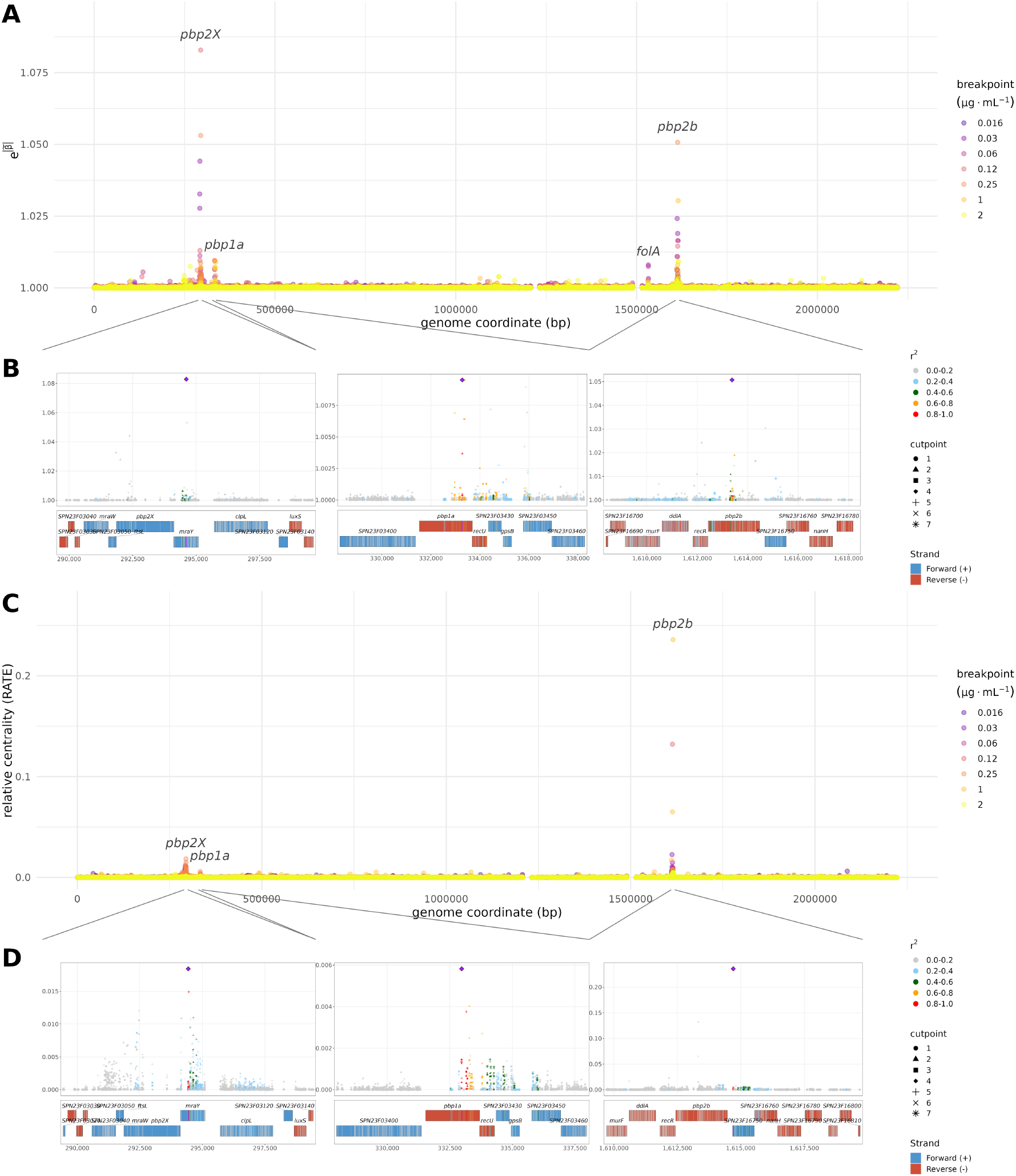
(A) Odds ratios calculated from the magnitude of variant coefficients’ medians, 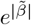, and (C) their relative centrality (RATE) values, for 32,406 variants associated with benzylpenicillin resistance in *S. pneumoniae* isolates using a partial proportional odds model (PPOM). (B,D) Locus zooms at the *pbp2x, pbp1a*, and *pbp2b* loci, for odds ratios and RATE values respectively. MIC breakpoints were calculated using the doubling dilutions (minimum 5% frequency) phenotype discretization strategy, which uses every doubling dilution containing a minimum of 5% of isolate phenotypes (S3). Coefficients from different breakpoints are overlayed.

In addition to the expected PBPs, we also found variants with high RATE values in the dihydrofolate reductase gene *folA* (*dyr*), a locus that was previously identified by Chewapreecha et al. In their study of epistatic interactions, Skwark etal. found strong long-range couplings of variants in dihydrofolate reductase and the PBPs, despite no obvious mechanism that would suggest *folA* is capable of increasing penicillin resistance, and despite their locations being far apart on the genome. Due to the strong link, Skwark et al. hypothesized that these two loci are coselected, presumably due to penicillin-trimethoprim co-threatment, which would explain why we find this signal even though it is almost certainly spurious.

When examining the *pbp2x* locus more closely, we see that the allele with the highest effect size is found in *mraY* instead of *pbp2x* in the most granular discretization (S9, Figure 3). *mraY* encodes a phospho-N-acetylmuramoyl-pentapeptide-transferase and is also involved in cell wall biogenesis (Skwark et al. 2017). This *mraY* variant does not appear to be in strong linkage with many *pbp2x* variants despite *pbp2x* being directly upstream; this is in contrast to the top variant at the *pbp2b* locus, which is located in the adjacent gene *SPN23F16750* but exhibits strong linkage with variants in *pbp2b*. Skwark et al. found *mraY* to exhibit strong connections to the PBPs, and noted that it was previously predicted that variants in this transferase that are associated with *β*-lactam resistance could be compensatory mutations that ameliorate the costs of evolving *β*-lactam resistance. *mraY* was also found to be significant in both this dataset and a larger 3,085-isolate dataset (Chewapreecha et al. 2014). However, laboratory validation is required to determine whether variants within *mraY* are likely to be truly causal or to result from linkage.

Low-level pneumococcal *β*-lactam resistance is known to involve loci beyond the PBP transpeptidases themselves, including cell-wall biosynthesis and integrity factors functionally coupled to those targets (for example, the CiaRH regulatory system) (Mascher et al. 2006). Because pneumococcal *β*-lactam resistance evolves largely through homologous recombination that generates mosaic PBP alleles, mutations in recombination machinery may also affect *β*-lactam resistance (Nishimoto et al. 2022). While the three canonical PBP loci and *folA* dominated the GWAS, several other genes appeared in the top 30 most significant genes (as measured by the highest-magnitude effect size of all variants within the gene across all cutpoints) in at least two discretization strategies using the PPOM (S17, S18, S21, S22). These included *ddl* (d-Ala–d-Ala ligase), which ligates two d-Ala residues into the d-Ala–d-Ala dipeptide, and *murF*, the cytoplasmic Mur ligase that adds that dipeptide onto UDP-MurNAc-tripeptide to complete the UDP-MurNAc-pentapeptide (the precursor of peptidoglycan), both of which are adjacent to *pbp2b* in the *dcw* (division and cell-wall) cluster (Garde et al. 2026; *Mechanism of action of penicillins* 2026); two further *dcw* genes, the aforementioned *mraY* and *mraW*, which are adjacent to *pbp2x* (Massidda etal. 1998); *clpL* (Clp ATPase/chaperone), a heat-shock protein reported to modulate PBP2x, cell-wall thickness, and penicillin tolerance in *S. pneumoniae* (Tran et al. 2011); division and cell-shape machinery that acts with the PBPs at the septum, namely *gpsB* and *ftsL*, which help coordinate septal PG synthesis with the class-B PBPs (Land et al. 2013; Rued et al. 2017), and *mapZ*, which positions the FtsZ ring and marks the division site (Fleurie et al. 2014); *recR*,a RecFOR recombination mediator (Nirwal et al. 2023); and *recU*,a Holliday-junction resolvase involved in homologous recombination, DNA repair, and chromosome segregation (Pereira et al. 2013). With the exception of *ddl, murF*, and *mapZ*, mutations in these genes were previously reported as associated with penicillin resistance in Chewapreecha et al.

### PPOM recovers order of stepwise acquisition of resistance variants

Resistance to the *β*-lactam amoxicillin has been found to emerge through a stepwise accumulation of mosaic alleles in the three canonical PBP genes; Gibson et al. demonstrated that initial mutations in *pbp2x* reduce amoxicillin susceptibility, followed by alterations in *pbp2b*, and culminate in high-level resistance only when *pbp1a* is also replaced with resistant alleles (Gibson et al. 2022). Interestingly, the median effect size coefficients recovered across cutpoints for variants in these genes display this pattern, including for the most granular discretization (Figure 4, S9, S10). This indicates that epistatic interactions can potentially be detected by identifying increases in the importance of certain variants at different phenotypic breakpoints.

**Figure 4.**
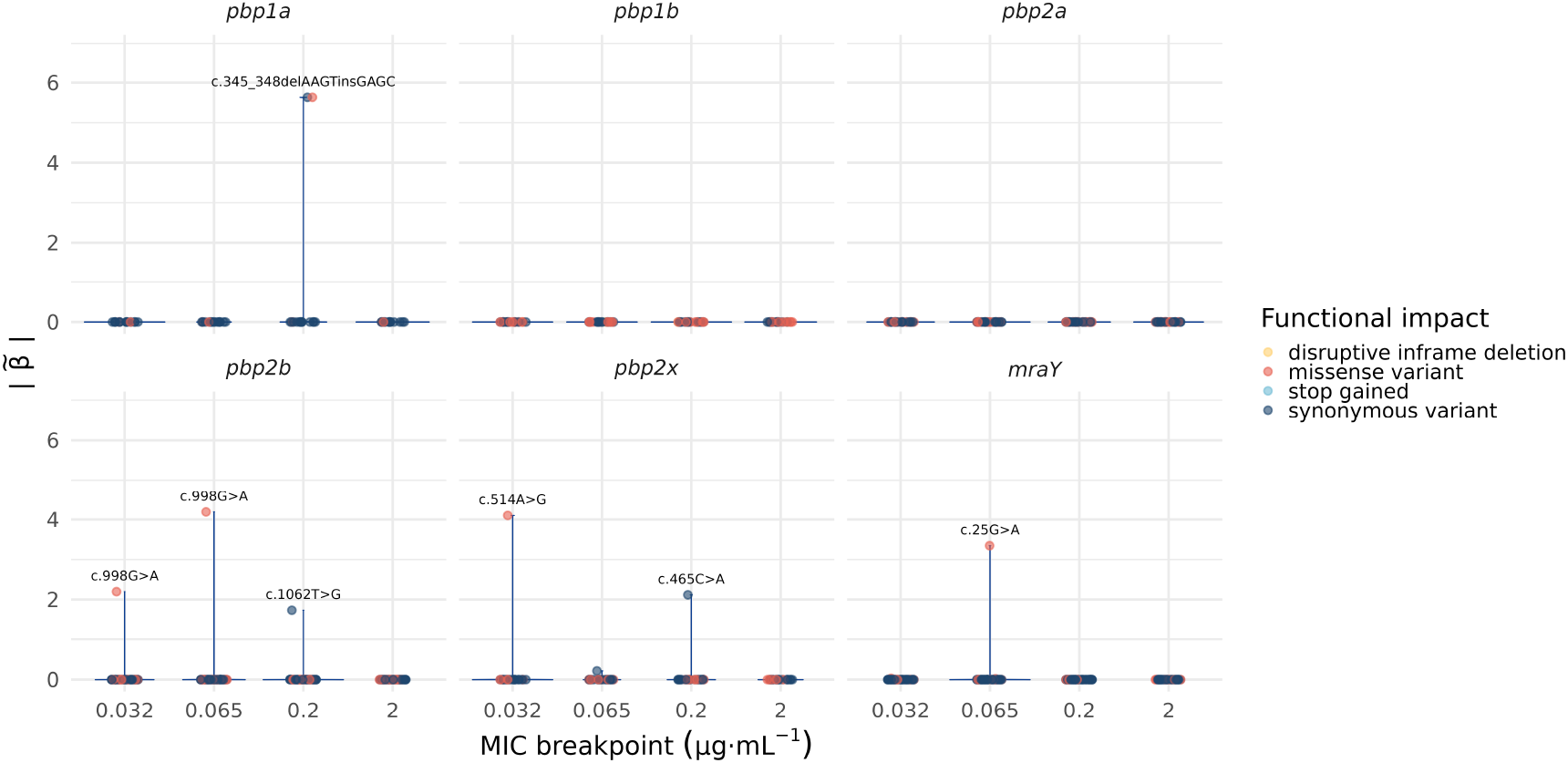
Median coefficient magnitude, 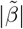, of all variants located within several key genes known to be associated with benzylpenicillin resistance in *S. pneumoniae* (*pbp2x, pbp2b, pbp1a*, and *mraY*) as well as those believed not to be involved in resistance (*pbp1b, pbp2a*). A violin plot of the coefficient distribution for each gene is shown in blue. Effects were fitted in a partial proportional odds model (PPOM) using MIC breakpoints calculated using the dilution minima phenotype discretization strategy, which places breakpoints at local minima in the phenotype MIC distribution (S3).

Interestingly, we also find that variants in *pbp1a* have the highest effect sizes in the MIC interval immediately preceding 1µg mL^−1^ in both the least and most granular datasets (0.20µg mL^−1^ and 0.25µg mL^−1^, respectively); this aligns with previous findings by Zhou et al., who reported that all tested strains that displayed penicillin MICs 1µg mL^−1^ had substitutions in *pbp1a* active sites. In other words, beyond replicating the order of stepwise accumulation of mutations, these findings indicate that the particular MIC breakpoint at which effect sizes have the greatest magnitude may be informative of the real MIC they confer. However, more validation would be needed to confirm this.

While the *pbp2x* and *pbp2b* loci dominate the effects in the logistic model, an additional RATE peak was observed in the region of the serine–threonine protein kinase *stkP* (*stk1*) (S16). Excluding variants in the *pbp2x* and *pbp2b* regions, the nine next most significant genes by RATE were located in the 28 kbp genomic window surrounding *stkP* (*fmt, sstT, stkP, mvaA, mvaS, sun, yqeG, ykuT*), with the exception of *ribC*. The top variants were missense variants Glu75Asp in the formyl-methionyl transferase *fmt*, Ile281Val in the single eukaryotic-type serine–threonine protein kinase (*stkP*), and Leu481Met in the serine–threonine transporter (*sstT*/*SPN23F17610*).

*stkP* regulates peptidoglycan synthesis and cell division and is responsible for localizing *pbp2x* to the division septum through interaction with its extracellular PASTA domain (Beilharz et al. 2012; Morlot et al. 2013). Loss of function in *stkP* results in hypersensitivity to penicillin, indicating that *stkP* acts upstream of the PBPs in the *β*-lactam response. However, no resistance-associated *stkP* alleles were detected among clinical isolates, whose MICs were instead determined by their PBP alleles (Dias et al. 2009). Whether this association is truly causal of penicillin resistance remains uncertain, as the effects of missense mutations have not been characterized. Similarly, Min et al. found that missense mutations in *fmt* induced resistance to the antibacterial peptide deformylase (PDF) inhibitor GSK1322322 at high frequency in 6 of 21 *S. pneumoniae* strains tested, but noted that these mutations were associated with severe *in vitro* and/or *in vivo* fitness costs (Min et al. 2015).

### Trimethoprim resistance in *S. pneumoniae*

In *S. pneumoniae*, trimethoprim resistance is conferred by mutations in the dihydrofolate reductase gene *folA* that reduce the binding affinity of trimethoprim without affecting dihydrofolate binding, the most prevalent of which is Ile100Leu. In clinical settings, trimethoprim is almost exclusively administered in conjunction with sulfamethoxazole (a combination known as co-trimoxazole). Therefore, variants in the *folP* gene encoding dihydropteroate synthase (DHPS) implicated in sulfamethoxazole resistance are expected to appear in trimethoprim resistance GWASs despite not being causal of trimethoprim resistance (Adrian and Klugman 1997; Safari etal. 2021). In contrastto other pneumococcal antimicrobial resistance mechanisms in *S. pneumoniae*, trimethoprim resistance appears to exclusively involve target-gene mutations; to our knowledge, no compensatory mutations in other genes have been documented, indicating trimethoprim resistance imposes minimal fitness cost.

Variants in *folA* and *folP* were recovered at very high effect sizes and RATEs in all models and discretization strategies (Figure 5, Figure, 6, S19S23, S26). Within *folP*, both POM and PPOM models identified Ser4Ile as the most significant variant (PPOM: RATE 0.0992). Sulfamethoxazole resistance in pneumococcal *folP* is conferred by short in-frame insertions and duplications in the sulfonamide-binding region spanning residues 58–67 (Maskell, Sefton, and LM Hall 1997; Haasum et al. 2001; Cornick et al. 2014); however, no in-frame insertions were called in residues 58–67 (or elsewhere in *folP*), so Ser4Ile’s significance may reflect linkage with uncaptured variation. In *folA*, all models identified either His81Gln (PPOM: RATE 0.0224–0.0337) or Ser70Pro (POM: RATE 0.0177–0.0206) as top variants depending on binning strategy. Missense mutations at these positions directly alter the trimethoprim-binding pocket of DHFR; His81Gln lies within the trimethoprim resistance-determining region of *folA* found approximately between codons 75–115. Several further N-terminal substitutions co-occur in resistant clinical isolates but are dispensable for the resistant phenotype, as the canonical Ile100Leu substitution within this region is on its own sufficient to confer resistance (Adrian and Klugman 1997; Pikis et al. 1998; Maskell, Sefton, and LMC Hall 2001). In this dataset, Leu100Ile exists in strong LD with a conserved set of co-occurring coding changes (*r*^2^ with His81Gln = 0.94, Pro111Leu = 0.89, Phe135Leu = 0.88, Leu31Trp = 0.83, Ser70Pro = 0.69), reflecting the haplotypic structure described in resistant strains rather than independently segregating sites (Maskell, Sefton, and LMC Hall 2001; Cornick et al. 2014). Although His81Gln is in strong linkage disequilibrium with Leu100Ile (*r*^2^ = 0.94), the presence of two additional synonymous alleles at the Leu100Ile site reduced this LD to *r*^2^ = 0.84, attenuating its phenotypic association. This highlights the importance of splitting multi-allelic sites when performing fine-mapping. In addition, beyond obfuscating true causal variants, combining all non-reference alleles can re-introduce population structure confounding as the multi-allelic locus correlates with the reference genome used.

**Figure 5.**
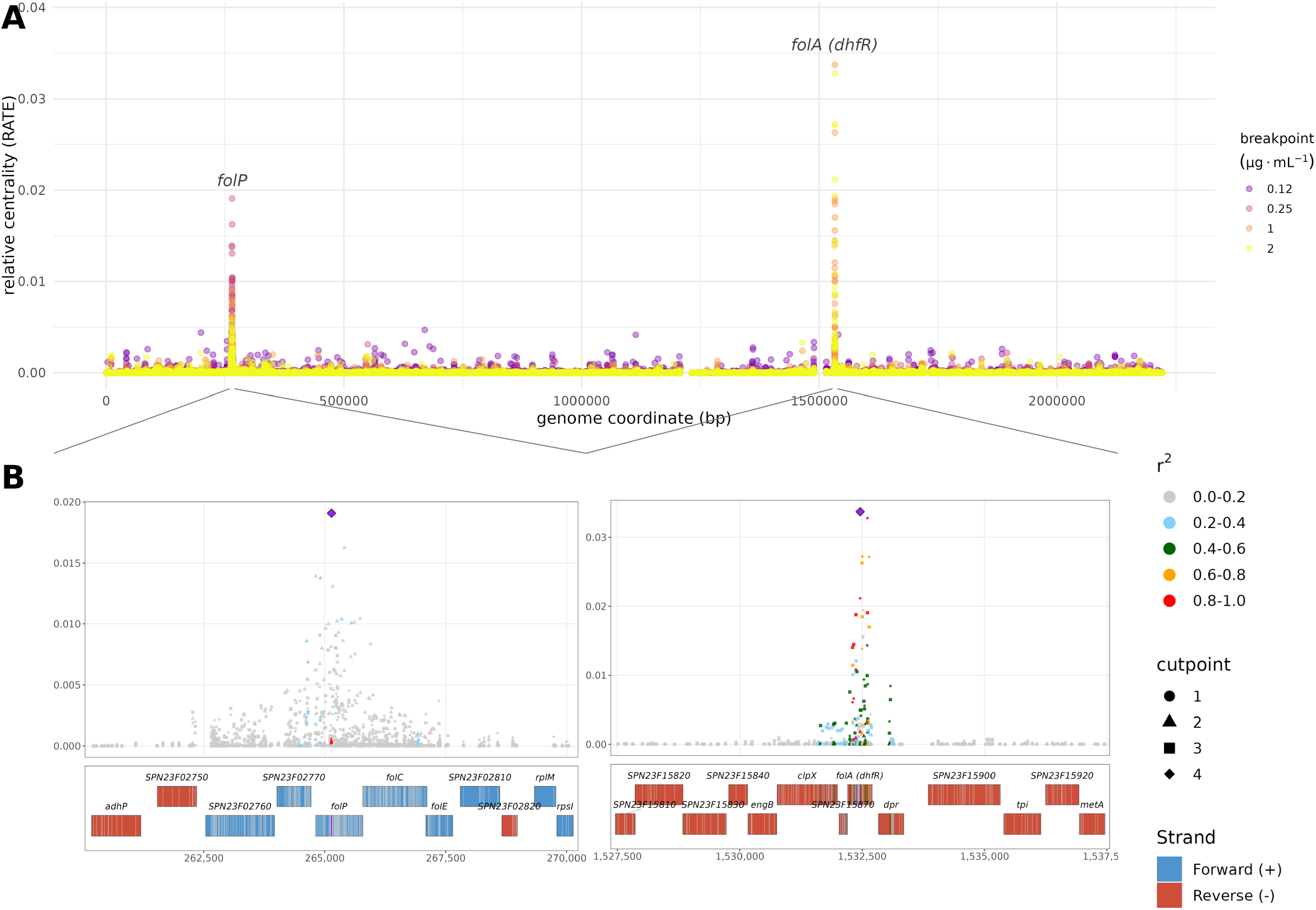
(A) Relative centrality (RATE) values for 32,406 variants associated with trimethoprim resistance in *S. pneumoniae* isolates using a partial proportional odds model (PPOM). (B) Locus zoom of RATE values at the *folP* and *folA* loci. Trimethoprim MIC breakpoints were calculated using the doubling dilutions (minimum 5% frequency) phenotype discretization strategy, which uses every doubling dilution containing a minimum of 5% of isolate phenotypes (S4).

**Figure 6.**
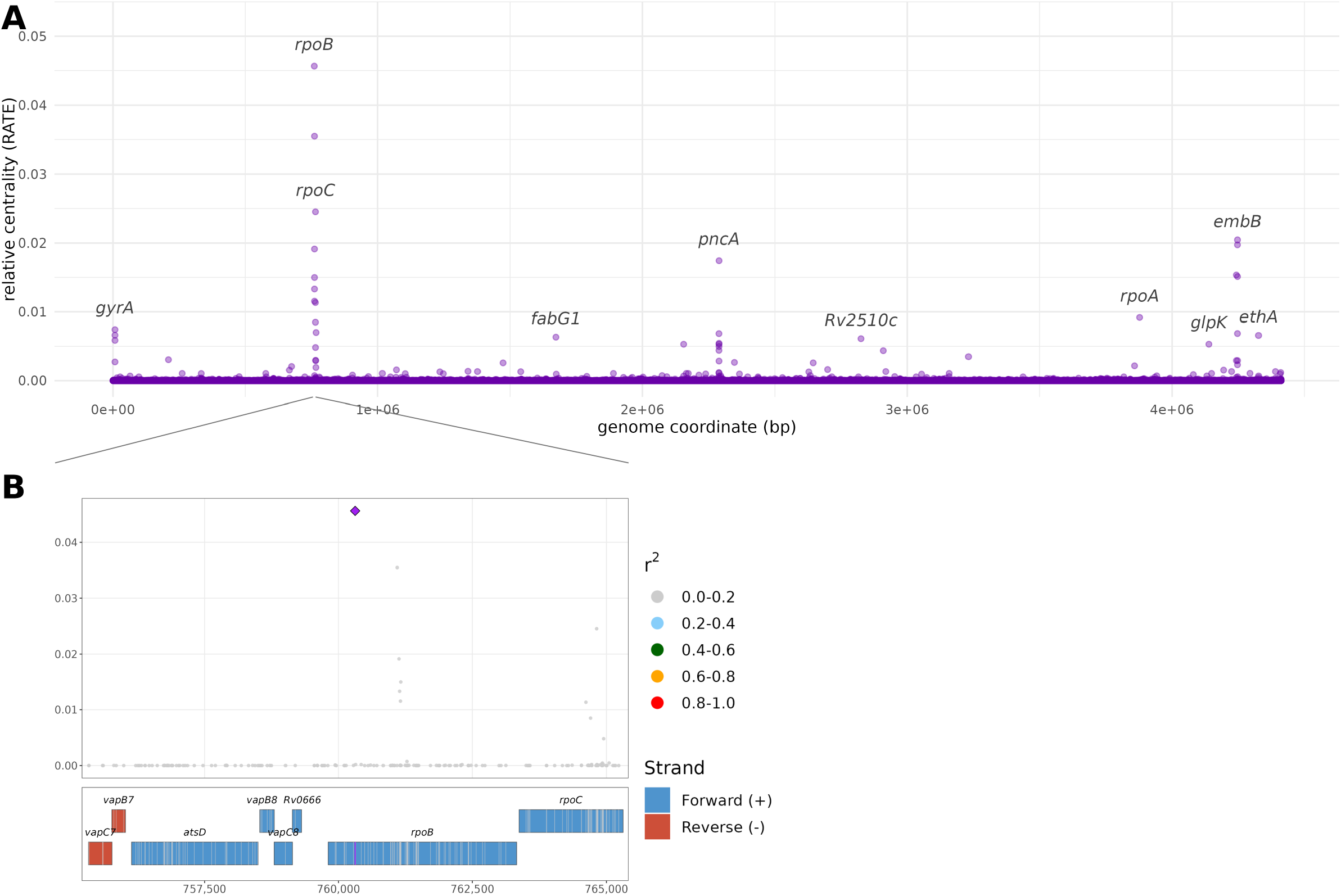
(A) Relative centrality (RATE) values for 75,272 variants associated with rifampicin resistance in *M. tuberculosis* isolates using a logistic regression model. (B) Locus zoom of RATE values at the *rpoB* locus. S5).

### Rifampicin resistance in *M. tuberculosis*

While *rpoB* mutations are the main driver of rifampicin resistance in *M. tuberculosis*, they can incur a significant fitness cost, which is often ameliorated by compensatory mutations in *rpoA* and *rpoC*, which have been found to offset the fitness cost of *rpoB* RRDR mutations by restoring transcription efficiency (P Ma et al. 2021; Zuo et al. 2018; Z Xu et al. 2018); there is also some evidence suggesting epistatic interactions with *gyrA* can confer high-level resistance (Trauner et al. 2021; Brandis and Hughes 2018).

For all binning strategies, the POM and PPOM models assigned variants in *rpoB* the highest effect size and RATE values. While the particular top variants differed by binning strategy and POM/PPOM, all PPOM models identified Ser450 in *rpoB* as the most significant variant (ST5, S20, S24). Missense mutations at position 450, particularly Ser450Leu and Ser450Trp, directly alter the rifampicin-binding pocket of RNA polymerase beta subunit, reducing rifampicin affinity and conferring high-level resistance; in some populations, these mutations account for *>* 95% of rifampicin resistance mutations present within the 81-bp RRDR (P Ma et al. 2021).

The POM models did notdiscover *rpoA* and *rpoC* with high effect sizes, with the exception of POM doubling 10% which discovered *rpoC* Pro1040 (RATE 0.0101), but all PPOM models found both genes with high RATE values in at least one cutpoint (ST5, S27, S20). The POM and PPOM models identified Thr187Ala and Val183 in *rpoA* as the top variants depending on the binning strategy (RATE 9.28 ×10^*-*5^–0.00546); notably, while Thr187Ala is an established compensatory variant (Napier et al. 2023), Val183 has not been previously reported in the literature and may represent a novel compensatory mutation. In *rpoC*, the PPOM models identified Asn698Ser across doubling ≥5% and ≥10% models (RATE 0.0253–0.0349), which is a canonical compensatory variant explicitly documented among established compensatory mutations (Napier et al. 2023).

Beyond these major known genes, additional genes that appeared with high effect sizes and RATE values in multiple models can be broadly sorted into either 1) mutations in genes known to confer resistance to other antimicrobials or 2) putative compensatory mutations. The variants found with high significance most frequently across models and discretizations included Ser315 in *katG*, which confers isoniazid resistance in approximately 64% of INH-resistant isolates worldwide (Lempens et al. 2018); Ser94Ala in *inhA*, an alternative pathway to isoniazid resistance that reduces isoniazid–NAD adduct binding to the enoyl-ACP reductase target (Eyles 2004); Met306Leu/Val and Asp354Ala/Gly in *embB*, conferring ethambutol resistance in 77.4% of EMB-resistant isolates (Moure et al. 2014); Met1 frameshift and stop-gained variants in *pncA*, where 88% of pyrazinamide-resistant MDR strains harbored *pncA* mutations (Pang et al. 2017); and Asp94Ala/Gly/Val in *gyrA*, found in 83–87% of phenotypically fluoroquinolone-resistant isolates ina systematic review of 3,846 isolates across 18 countries (Avalos et al. 2015). Putative compensatory mutations beyond *rpoA* and *rpoC* included Val192 variants in *glpK* (a glycerol kinase for nutrient adaptation), Ser403 frameshift and Leu451 polymorphisms in *amt* (an ammonium transporter for nitrogen uptake under starvation), and Val183 in *rpoA*, alongside the major *rpoC* variants, which collectively restore growth rates in RIF-resistant strains; these compensatory mutations enable survival during chronic infection and are preferentially selected in RIF-resistant isolates, with strains carrying S450L showing a 41.7% likelihood of accumulating compensatory mutations (Zuo et al. 2018).

Rifampicin resistance in *M. tuberculosis* illustrates clearly how differentmodels uncover differentstructures of varianteffects. Using a logistic regression framework with this dataset, many variants appear involved in increasing the log odds of resistance above the EUCAST breakpoint, including those with very large effects that immediately confer highlevel resistance (such as in *rpoB*), those that confer resistance from below the breakpoint to above the breakpoint but may have no effect on resistance given a background resistance level above the breakpoint (such as in *rpoA* and *rpoC*), and those with small, consistently-positive additive effects (such as non-causal variation in *embB* and *katG*) (Figure 6, S16). In contrast, the proportional odds model identifies variants that confer consistent increases in log-odds across multiple breakpoints, which may or may not represent meaningful differentiations in resistance. Using the POM, we see a reduced number of loci identified compared to the logistic model, but an increase in the relative importance of the loci; *rpoB* is by far the most prominent locus using these models (S27).

### Heritability estimates differ across MIC breakpoints in PPO models

Because variant effects are estimated at each MIC breakpoint in the PPOM, narrow- and broad-sense heritability, 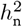 and 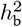, are calculated at each cutpoint. Comparing these per-breakpoint heritability estimates to those estimated jointly across breakpoints using the POM, we see that some breakpoints display considerably higher or lower heritability than the joint estimate. For example, the narrow-sense heritability estimate for penicillin resistance in *S. pneumoniae* using the POM with doubling dilutions (*≥*5% min. frequency) was 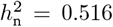 (95% credible interval, *CI*, 0.351–0.675); using the PPOM, this dataset displays the lowest heritability at MIC *≤* 0.016µg mL^−1^ 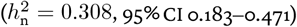 and the highest heritability at MIC *≤* 0.25µg mL^−1^ 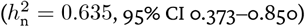.

The interpretationof the per-cutpointheritability is analogoustothe heritability estimatedfromthe logistic model; it tells us how informative the variant predictors are, given our model and its population structure correction, of the log-odds of resistance falling above or below that MIC breakpoint. In this sense, we can interpret differing heritability estimates as a quantitative measure of the genetic variants’ ability, given an MIC breakpoint, to meaningfully distinguish resistance phenotypes. As expected under this framework, we see that breakpoints centered at EUCAST break-points tend to display very high heritabilities in line with known estimates: in penicillin resistance in *S. pneumoniae*, the highest narrow- and broad-sense heritability fall at MIC *≤* 0.06µg mL^−1^ in the “minima” method (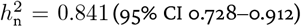, 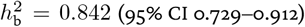), and in trimethoprim resistance in *S. pneumoniae* the highest heritability estimates fall at MIC *≤* 0.25µg mL^−1^ in the doubling (*≥*10% min. frequency) 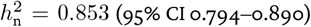, 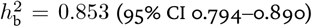. When examining the cutpoint distributions of those breakpoints with low heritabilities, we see that the cutpoints were not well-distinguished despite the strong priors placed on them, leading to mild cutpoint collapse (S28). “Collapsed” cutpoints, those estimated to have nearly identical latent-scale positions, produce negligible predicted probability of the interval they delimit regardless of the data. Low per-cutpoint heritability and cutpoint collapse thus both imply that a breakpoint has failed to partition the latent resistance continuum into distinguishable phenotypes. The same breakpoints showing both collapsed cutpoints and low heritability, while those near established clinical breakpoints show neither, supports the use of per-cutpoint *h*^2^ as a diagnostic of whether a proposed dilution boundary separates meaningful phenotypes.

We also find that phenotype discretization matters significantly in determining the heritability estimates of individual cutpoints. For example, when discretizing the penicillin resistance dataset in *S. pneumoniae* using the doubling (5% min. frequency) method, 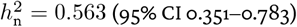 at MIC *≤* 0.06µg mL^−1^, while using the “minima” method causes this cutpoint to be estimated as having a heritability of 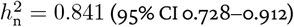. This suggests that the inclusion of non-informative breakpoints reduces the ability of the model to capture signal from true causal variants, despite the cutpoints being estimated separately, potentially due to the joint estimation of lineage cluster and subcluster-based population structure. This is supported by the POM heritability estimates, the credible intervals of which appear to span the interval between the highest and lowest cutpoint heritabilities in the corresponding PPOM. Penicillin resistance in *S. pneumoniae* has previously been reported to have a narrow-sense heritability in the range 0.67–0.83, with Mallawaarachchi et al. reporting an LD-score regression (LDSC) estimate of 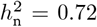 for penicillin MIC phenotypes, and Mai et al. reporting a heritability of ≥80% using a continuous (inhibition-zone) phenotype (Mallawaarachchi et al. 2022; Mai et al. 2021). This matches our logistic model’s 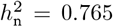 (95% CI 0.683–0.899) and the distribution of PPOM scores (S11). Similarly, co-trimoxazole resistance in *S. pneumoniae* was reported with a heritability of ≥71% using a continuous phenotype by Mai et al., matching our logistic model’s 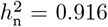 (95% CI 0.801–0.984) and PPOM scores (S11).

Rifampicin MIC heritability in *M. tuberculosis* has previously been estimated at 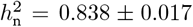 using a linear mixed model with a genetic relatedness matrix (GEMMA) (Farhat et al. 2019). More recently, the larger CRyP-TIC dataset used in our analysis was estimated to have MIC heritabilities of 94.6–94.9% for isoniazid and rifampicin (Consortium 2022). In our models, the highest heritability was not observed in the logistic 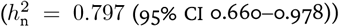 or POM (4-fold ≥5% min. frequency, 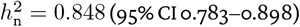 butinthe MIC *≤* 0.06µg mL^−1^ cutpoint of the doubling 10% min. frequency dataset 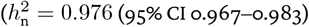. Given the considerations outlined above, this could be symptomatic of better breakpoint selection in this discretization, which had consistently high heritabilities across all phenotype categories (min. 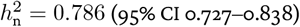, see S11).

Lastly, we find that broad-sense heritability doesn’t appear to be significantly higher than narrow-sense heritability in any model, which may be due to the relatively small contribution of the population-structure variance *V*_pop_ = Var(**X**_sub_***ω***) to the total genetic variance.

### Predicting MICs using the partial proportional odds model

Another advantage of Bayesian methodology is the ease with which an inference model can be adapted to include prediction from the fitted posterior. Here, we display the results of minimum inhibitory concentration interval prediction in *S. pneumoniae* and *M. tuberculosis* using the PO and PPO models (Figure 7). Two train:test splits were performed. In the first, a random subset of samples was withheld using an 80:20 train:test split; in the second, all samples from the largest lineage subcluster comprising < 20% of the total number of isolates were used as the test set (see “Methods”).

**Figure 7.**
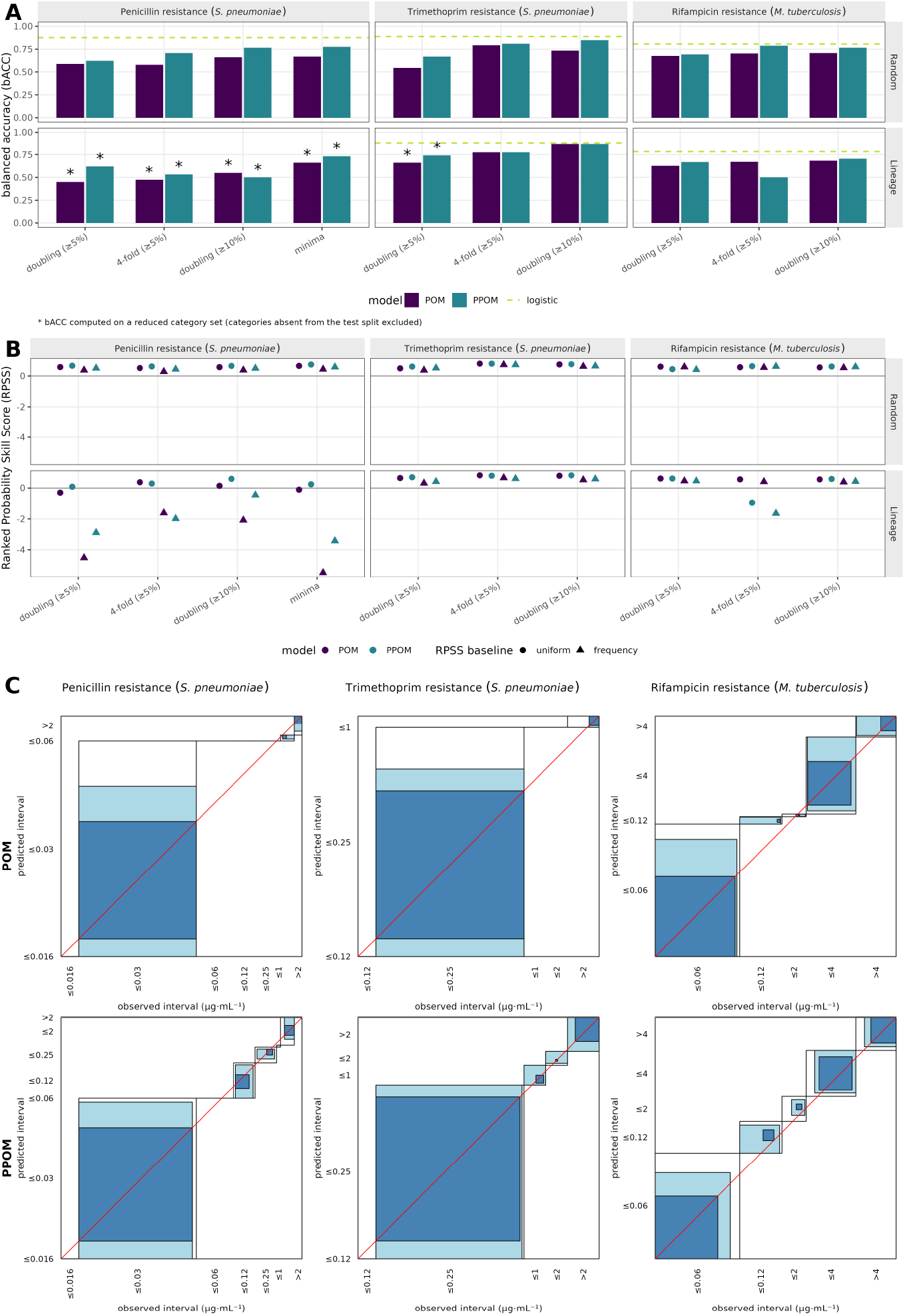
Prediction of ordered categorical phenotypes in *S. pneumoniae* and *M. tuberculosis* with varying MIC binning strategies. (A) Balanced accuracy (bACC) for a random 80:20 train: test split (top) and a withheld lineage subcluster test dataset (bottom) using a proportional-odds model (POM), partial-proportional-odds model (PPOM), and a logistic model in which the MICs were binned into only two categories using the EUCAST or clinical breakpoint (see Methods and S3, S4, S5). As some lineage subclusters did not contain all possible MIC categories, categories absent from the test split were excluded from bACC estimation in some splits, which are marked with an asterisk. (B) Ranked Probability Skill Score (RPSS) for the POM and PPOM, calculated against both a MIC category probability baseline drawn from uniform category frequencies and from the true category frequencies within the dataset. (C) Agreement plots displaying the observed versus predicted MIC intervals from the random 80:20 train:test splits using the most granular discretization method (doubling dilutions with ≥5% frequency). Dark blue indicates a correct classification, light blue an adjacent category, and white two or more categor3ie5s away. The alignment of corners on the red diagonal line shows over- or underestimation of each category frequency.

### Comparison to logistic and continuous models

Reducing MIC phenotypes to a binary resistant-sensitive call at the clinical breakpoint gave the highest predictive accuracy of the three encodings, as expected given the reduced difficulty of classification. Balanced accuracy on the held-out random split was 0.88 for penicillin, 0.89 for trimethoprim and 0.81 for rifampicin, with AUC values of 0.88–0.99 (ST6). The resistance probabilities were also reasonably well calibrated, with Brier scores on the random split of 0.11 for penicillin, 0.04 for trimethoprim and 0.14 for rifampicin (lower is better), indicating that the predicted probabilities would be meaningful enough to flag borderline isolates for confirmatory phenotyping. We found that accuracy was highly asymmetric between specificity (recall of susceptible isolates), which was consistently high at 0.95–0.96 on the random split, and sensitivity (recall of resistant isolates), which was lower at 0.67–0.82 on the random split. For this reason, the very major error rate (in which a resistant isolate is predicted as susceptible) was often substantially higher than the major error rate. For example, rifampicin resistance in *M. tuberculosis* had VME = 0.33 and ME = 0.05. This has significant implications for clinical practice, in which VME has a critical impact on patient welfare and ME on antibiotic stewardship (Bartoletti et al. 2022).

At the opposite extreme, we represented MIC phenotypes on the linear scale using the continuous model. Root-mean-squared error was 1.8–2.8 doubling dilutions and the coefficient of determination was close to zero on the held-out random splits (*R*^2^ = 0.04, 0.04 and 0.13 for penicillin, trimethoprim and rifampicin), suggesting that the regression explained little of the sample-to-sample variance in exact MIC even where it classified resistance well. Mean absolute error was lower than RMSE for every drug (1.69 vs 2.03 for penicillin, 1.38 vs 1.76 for trimethoprim and 2.52 vs 2.81 for rifampicin), indicating that the RMSE was primarily inflated by a minority of isolates with large prediction errors; this could result from the strong variable selection shrinking some true causal variants to 0, especially in regions with multiple highly correlated resistance variants. Essential agreement ranged from 0.14 to 0.52 on the random split (ST7), mirroring the agreement between adjacent categories seen in the ordinal models. We also note that narrow-sense heritability estimated from the continuous model was consistently lower than that estimated from the logistic and ordinal models on the same data (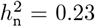, 0.001 and 0.06 for penicillin, trimethoprim and rifampicin respectively; S11), indicating that the model was less able to capture the genetic signal underlying the phenotype and may benefit from a different parameterization.

In sum, predictive performance scaled inversely with the resolution of these alternative representations of the same underlying MIC. The two-class breakpoint model displayed the highest predictive performance, the ordinal dilution-category models intermediate, and the continuous MIC regression the worst, with balanced accuracy falling as the number of categories increased (for example, trimethoprim resistance with the random train: test split fell from 0.89 with two classes to 0.81 with three categories and 0.67 with five categories).

As in the ordinal models, prediction on an unobserved lineage subcluster was consistently poorer in the logistic model and frequently poorer in the continuous model, indicating that population structure correction with lineage subclusters rather than phenotype discretization is responsible for the decrease in predictive accuracy. For example, balanced accuracy fell from 0.81 to 0.78 and F_1_ from 0.77 to 0.68 for rifampicin resistance prediction in *M. tuberculosis* and AUC from 0.99 to 0.92 for trimethoprim resistance in *S. pneumoniae* in the logistic model. This trend was less consistent in the continuous model, in which *R*^2^ fell from 0.04 to 18.6 and essential agreement from 0.24 to 0.04 for penicillin resistance in *S. pneumoniae*, but rifampicin resistance in *M. tuberculosis* improved on RMSE (2.81 to 2.55) and MAE (2.52 to 2.36) while worsening on *R*^2^ and essential agreement, and trimethoprim resistance in *S. pneumoniae* improved on every metric reported, with RMSE falling from 1.76 to 1.42 and essential agreement rising from 0.52 to 0.72 (ST7). Additionally, calibration improved on the lineage split for all three logistic datasets (Brier 0.104 versus 0.110, 0.023 versus 0.037 and 0.117 versus 0.135; ST6), consistentwith the withheld subclusters being phenotypically more homogeneous than a random sample.

### PPOM exhibits improved prediction compared to POM by capturing intermediate-category isolates

Across all three drugs and every MIC discretisation, the partial proportional odds model (PPOM) predicted MIC category more accurately than the fully constrained proportional odds model (POM). Balanced accuracy (bACC), which weights every observed category equally and so is not dominated by the abundant extreme categories, was higher for PPOM than POM in all ten within-training (PPC) comparisons and in all ten held-out random-split comparisons (ST8, Figure 7). When examining performance on a random subsample, improvement in bACC ranged from +0.017 (*M. tuberculosis* with a rifampicin resistance phenotype, doubling dilutions with ≥5% min. frequency) to +0.129 (*S. pneumoniae* with a penicillin resistance phenotype, coarse binning), with comparable gains elsewhere(e.g. 0.788 to 0.703 for rifampicin resistance in *M. tuberculosis*, coarse binning). Ranked probability skill score RPSS followed a similar pattern in most comparisons, with the notable exception of rifampicin resistance in *M. tuberculosis* using doubling dilutions with ≥5% frequency, where the POM displayed higher RPSS (0.604 vs 0.441 against the uniform baseline) and PPV (0.586 vs 0.568) despite having a lower bACC (0.678 vs 0.695) (Figure 7). The only cases in which POM matched or exceeded PPOM were the leave-one-lineage-subcluster-out splits; for instance, prediction in rifampicin resistance in *M. tuberculosis* using the coarse binning strategy decreased from 0.670 using the POM to 0.501 with the PPOM. In this case, because the subcluster has not previously been observed, its effect is drawn from the prior 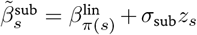 which shrinks toward the parent lineage effect 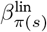 if that lineage was present in the training data, or toward zero 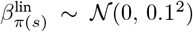 if the parent lineage was also unobserved (Equation 10, Equation 11); this leads to poorer predictive performance across all models, which may explain this discrepancy.

The underlying cause of the predictive performance improvement may reflect which types of variant effects are better captured by the PPOM. Because POM assumes a single shared odds ratio across all category cutpoints, it tends to place most probability mass on the highly resistant and highly susceptible categories and to collapse the sparser intermediate MIC categories onto these extremes. Relaxing the proportional-odds constraint for individual predictors as in the PPOM allows those intermediate categories to be predicted with appreciably higher frequency. This is visible in the agreement plots (Figure 7), in which POM concentrates agreement at the corners of the scale while PPOM fills the diagonal at intermediate doubling dilutions. This is also reflected in how PPOM is most clearly superior when scored with bACC, which rewards correct prediction of the under-populated middle categories rather than in metrics dominated by the majority classes, while this relationship was less apparent or even inverted (in the case of rifampicin resistance in *M. tuberculosis*) when scored with RPSS.

Finally, this relationship appeared both when predicting category probabilities for data the model was trained on (the posterior predictive check, or PPC) and for held-out subsets (the random and lineage splits), indicating that the additional intermediate-category structure captured by PPOM is not due to overfitting. Taken together, these results demonstrate thatrelaxing the proportional-odds assumption allows the model to resolve variation that moves isolates into the intermediate MIC categories that POM is structurally less able to represent.

## Discussion

Existing bacterial GWAS frameworks typically represent minimum inhibitory concentrations, which are ordered censored intervals, either as binary resistant-sensitive phenotypes that discard the resolution of the dilution series and are sensitive to the clinical breakpoint applied, or as a continuous log_2_ value that treats a censored interval as a point on an unbounded scale (Batisti Biffignandi et al. 2024). Both of these methods model a variant’s effect as constant regardless of genetic background, attenuating low-penetrance and resistance-dependent variants, such as compensatory mutations. Machine-learning approaches such as elastic net and random forest instead treat MICs as a classification task with unordered categories such that an isolate two dilutions from the truth is penalised identically to one adjacent to it, and the fitted model cannot take into account the latent concentration underlying the bins. Random forests also return variable importances rather than effect sizes with uncertainty, and the L1 component of an elastic net divides signal among correlated variants. In these implementations, population structure is also handled by sequence reweighting rather than by explicit lineage terms, so lineage effects cannot be separated from variant effects or used to estimate heritability. Curation-based approaches such as the WHO SOLO method involve building a catalogue of graded variants by testing each mutation univariately against a binary phenotype using only isolates in which it is the sole candidate variant, then assigning confidence grades using rules external to the data, including literature evidence (Kulkarni et al. 2025). Kulkarni et al. showed that multivariable L2-penalised logistic regression on 52,567 *M. tuberculosis* genomes recovers 450/457 (98.5%) of SOLO’s resistance-associated variants, grades 221 more, and removes the need to pre-specify neutral variants, because co-occurring mutations are fitted jointly. However, it restricts testing to curated Tier 1 candidate genes, drops isolates with intermediate or missing calls, and assumes additive effects on a logistic scale (using binary phenotypes). These methodological limitations are critical to resolve given the drastic increase in the availability of large MIC datasets, with the release of notable datasets such as the CRyPTIC consortium’s 11,622 MIC-phenotyped isolates used in this analysis (Consortium 2022) and the new CABBAGE database with approximately 1.7M AMR-phenotyped genomes across WHO priority pathogens (Dickens et al. 2026). Despite the growing need for scalable GWAS methods tailored for MIC phenotypes, no MIC-based bacterial GWAS method has yet been developed.

Bayesian ordered logistic regression with partial proportional odds provides a compelling alternative GWAS framework to existing methods for minimum inhibitory concentration phenotypes. The models presented here allow for fitting ordered censored intervals explicitly and jointly across variants with recovery of per-threshold effects and prediction from the same model without reliance on external grading rules. We found that ordered logistic PPO models recovered known resistance variants in *S. pneumoniae* and *M. tuberculosis* for benzylpenicillin, trimethoprim, and rifampicin across every phenotype discretization tested. Beyond the canonical high-penetrance variants, PPOM captured variants with effects conditional on a resistant background genotype that the POM was unable to identify. For instance, every PPOM fit identified both compensatory mutation loci *rpoA* and *rpoC* at high RATE in at least one cut-point in the rifampicin resistance GWAS in *M. tuberculosis*, including the documented compensatory substitution *rpoC* Asn698Ser (Napier et al. 2023), whereas the POM recovered neither with substantial effect except *rpoC* Pro1040 in a single discretization (ST5). Consistent with this, the PPOM exceeded the POM on balanced accuracy in all ten within-training and all ten held-out random-split comparisons (+0.017 to +0.129), with the gain concentrated in the sparse intermediate MIC categories that a proportional model collapses onto the extremes of the scale (ST8).

Relaxing the proportional-oddsconstraintadditionallyyieldedper-cutpointcoefficientsthatreproducethe known order of accumulation of stepwise penicillin resistance mutations. Using these threshold-specific effect sizes, we observe that the effect sizes of variants in *pbp2x* are highest at low MIC breakpoints, variants in *pbp2b* at intermediate breakpoints, and variants in *pbp1a* only at high breakpoints (Figure 4), recovering the experimentally determined sequence of Gibson et al. from carriage data alone. The breakpoint at which a gene’s effect is maximal also appears informative of the resistance level it confers, with *pbp1a* effects peaking immediately below 1 µg mL^−1^ in both the least and most granular discretizations, consistent with the finding that almost all strains at or above that MIC carry *pbp1a* active-site substitutions (Zhou et al. 2022). We therefore conclude that epistatic structure is detectable as a change in a variant’s importance across breakpoints without explicit pairwise interaction testing.

Estimating variant effects per breakpoint allows for the calculation of heritability at each breakpoint. We found that narrow-sense heritability varied substantially across cutpoints within a single fit, and the highest values fell consistently at or near established clinical breakpoints. Since a breakpoint with low heritability is one that no variants in the dataset can distinguish well, this provides a quantitative metric for whether a proposed dilution boundary separates meaningful phenotypes (S28). In the future, this could be used to reduce the number of serial dilutions required to capture genetic variants associated with resistance without reducing power.

We also introduce a hierarchical lineage subclustering approach to population structure correction. Genetic relatedness matrices are the standard method in both human and bacterial GWAS (Hoffman 2013), but scale quadratically with increasing sample size. Because the number of lineages in a species is bounded by its population structure rather than by sample size, the cost of the lineage-based method is close to constant as *N* grows, and scoring a new isolate requires only a lineage assignment rather than recomputing its relatedness to every training sample. When fitted to the same *S. pneumoniae* penicillin resistance dataset, the lineage cluster method recovered the same loci as the GRM-based method with comparable null distributions of effect sizes and RATE values (S15). When applied to the larger 11,622-isolate *M. tuberculosis* dataset, the resulting GRM was 2.7 GB (nearly twice the 1.43 GB required by the entire LD-pruned genotype matrix), and could not be serialized into Stan given the 2^31^ 1 byte limit, whereas the equivalent lineage encoding added only 7 lineage cluster and 78 subcluster columns, showcasing the utility of this method.

Finally, the model allows for prediction of MIC phenotypes with probabilities of each MIC interval. Direct comparison of predictive performance against these methods is limited by differences in species, drug, dataset and method of evaluation. On *K. pneumoniae* MICs binned into 4–10 unordered categories, the elastic net and random forest implemented by Batisti Biffignandi etal. (2024) achievedbalancedaccuracies of 0.58–0.93 and test-set *R*^2^ of 0.19–0.72, with training *R*^2^ consistently higher, indicating overfitting. Our PPO models reached bACC 0.62–0.85 on held-out random splits across 3–8 categories (ST8), with posterior predictive and held-out values close to one another, and additionally return a calibrated probability for every MIC interval (RPSS_unif_ 0.441–0.798 on the random splits). Both analyses show accuracy declining as categories increase, as their lowest balanced accuracies were for meropenem with ten concentration intervals (0.58–0.70), matching the reduction we observe for trimethoprim from 0.89 with two classes to 0.67 with five. Because RPS penalises a misclassification in proportion to its distance from the truth, it also supersedes the off-by-one correction (off-by-one accuracy 0.64–1 in Batisti Biffignandi et al.) that is otherwise needed to keep models with many categories from being unfairly penalised. Against catalogue-based methods, the multivariable regression of Kulkarni et al. attained a mean F_1_ of 70.2% across 15 drugs versus 69.2% for SOLO, with average gains of +3.2% in sensitivity against 1.0% in specificity and 1.6% in PPV. These estimates were computed on the data from which the gradings were derived, whereas our logistic F_1_ of 0.772 for rifampicin (0.677 on the withheld lineage; ST6) was computed from a held-out test dataset and a hypothesis-free genome-wide fit rather than curated candidate genes.

Our very major error rate for rifampicin (VME = 0.33) nonetheless remains far from the WHO target product profile of *>* 95% sensitivity, indicating some resistance variants were not detected, perhaps due to strong variable shrinkage. Finally, our continuous model displayed an *R*^2^ of 0.04, 0.04 and 0.13 against the 0.48–0.72 reported for random forests on real MICs (Batisti Biffignandi et al. 2024).

The strong regularized horseshoe prior used for locus selection was applied in the prediction models, but these may instead benefit from a less aggressive penalty such as a LASSO or ridge prior to improve predictive performance, as implemented in the elastic-netmode of Pyseer (Lees, Galardini, etal. 2018). Reparameterizing the model after fitting also presents an opportunity to improve prediction for novel lineage clusters and subclusters while removing the need for lineage identification inquery samples altogether; currently, atestisolate assignedto a lineage subcluster *s* that was unobserved during training inherits an effect drawn from its prior alone, 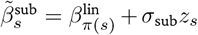, which shrinks toward the parent lineage effect 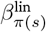, or toward zero 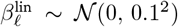 where the parent lineage is also unobserved (Equation 10, Equation 11).

The framework therefore addresses four limitations of current practice: it represents MICs as ordered censored intervals rather than binary or continuous values; it removes dependence on a fixed external breakpoint by estimating cutpoints under priors anchored to the dilution grid; it recovers threshold-dependent and conditional effects in one joint fit rather than through pairwise comparison of variants; andit provides both inference and prediction capabilities, including within the same model. All models are implemented in Stan and distributed with GHOST, an R workflow that runs the full analysis from linkage disequilibrium pruning through model fitting to downstream summaries (Figure 2), so that the methods described here can be applied to new datasets without reimplementation.

In the future, the models presented here could be expanded to incorporate additional parameters to improve discovery of novel variants and fine-mapping of hit loci. Examples of information that could be exploited include predicted functional impact of variants (for example, implementing harsher priors on synonymous mutations compared to loss-of-function mutations, or for mutations in noncoding regions), Tajima’s *D*, functional enrichment, and gene-based priors that incorporate copy number, functional category, *d*_N_/*d*_S_, and loss-of-function burden testing.

While we used small-variant calling to call SNPs and short indels against a reference genome, the models accept any binary genotype matrix and thus are immediately usable with unitigs, *k*-mers, and gene presence–absence matrices. Reference-free representations with unitigs would be particularly valuable for accessory-genome and mobile-element determinants, which a reference-based SNP analysis has poor ability to detect, and for species with very open pangenomes. Similarly, because the ordered models are agnostic to what the ordered categories represent, these models could be applied to other bacterial phenotypes recorded as censored intervals such as growth or virulence assays. Joint modeling of several antimicrobial resistance phenotypes could also be implemented in this framework, which could be valuable for the reduction of spurious signal due to co-selection.

Further work could also involve applying these methods to the substantially larger datasets made available through resources such as CABBAGE (Dickens et al. 2026). Datasets of that size are where the linear scaling of the lineage sub-cluster correction becomes decisive, and where the conditional and low-penetrance variants the PPOM is designed to detect should become statistically accessible.

Finally, as previously mentioned, multivariable logistic regression has recently been used to identify resistance mutations in *M. tuberculosis* with success compared to the WHO mutation catalogue (Kulkarni et al. 2025). Kulkarni et al. discuss MIC- and lineage-based methods as potential progressions to reduce reliance on grading rules and prior knowledge of resistance mutations. The ordered logistic PPO methodology with lineage subcluster covariates takes a substantial step towards bridging ordered categorical (MIC) phenotypes and lineage designations with a flexible, hypothesis-free model.

## Data and Code Access

All Bayesian models, the complete GWAS pipeline, and the Rust code used for LD pruning are available on GitHub at qtoussaint/bayesian-gwas, qtoussaint/gwas_workflow and https://github.com/bacpop/BacPrune-Rust. All additional code used to run analyses and generate figures is available on GitHub at qtoussaint/thesis_code. Data are publicly available and construction of each JSON used in the workflow is displayed in the relevant code, as data JSONs and Stan fits produced from the GWAS analyses exceed 500GB. However, these are available upon request and can be passed to the GWAS pipeline, or directly to the Stan model if no pre-pruning is desired.

## Acknowledgments

Thanks to Zamin Iqbal and Matthew Russell for their suggestions and advice.

## References

Adrian PV and Klugman KP. 1997. Mutations in the dihydrofolate reductase gene of trimethoprim-resistant isolates of Streptococcus pneumoniae. Antimicrobial Agents and Chemotherapy. 41: 2406–2413.

Anscombe FJ. 1956. On Estimating Binomial Response Relations.

Antimicrobial Resistance Collaborators. 2022. Global burden of bacterial antimicrobial resistance in 2019: a systematic analysis. Lancet (London, England). 399: 629–655.

Avalos E, Catanzaro D, Catanzaro A, Ganiats T, Brodine S, Alcaraz J, and Rodwell T. 2015. Frequency and Geographic Distribution of gyrA and gyrB Mutations Associated with Fluoroquinolone Resistance in Clinical Mycobacterium Tuberculosis Isolates: A Systematic Review. PLOS ONE. 10: e0120470.

Bartoletti M et al. 2022. Clinical consequences of very major errors with semi-automated testing systems for antimicrobial susceptibility of carbapenem-resistant Enterobacterales. Clinical Microbiology and Infection. 28: 1290.e1–1290.e4.

Batisti Biffignandi G, Chindelevitch L, Corbella M, Feil EJ, Sassera D, and Lees JA. 2024. Optimising machine learning prediction of minimum inhibitory concentrations in Klebsiella pneumoniae. Microbial Genomics. 10:

Beilharz K, Nováková L, Fadda D, Branny P, Massidda O, and Veening JW. 2012. Control of cell division in Streptococcuspneumoniaebytheconserved Ser/Thr proteinkinase StkP. Proceedings of the National Academy of Sciences. 109: E905–E913.

Brandis G and Hughes D. 2018. Mechanisms of fitness cost reduction for rifampicin-resistant strains with deletion or duplication mutations in rpoB. Scientific Reports. 8: 17488.

Bruchmann S, Dötsch A, Nouri B, Chaberny IF, and Häussler S. 2013. Quantitative Contributions of Target Alteration and Decreased Drug Accumulation to Pseudomonas aeruginosa Fluoroquinolone Resistance. Antimicrobial Agents and Chemotherapy. 57: 1361–1368.

Buchfink B, Reuter K, and Drost HG. 2021. Sensitive protein alignments at tree-of-life scale using DIAMOND. Nature Methods. 18: 366–368.

Cantalapiedra CP, Hernández-Plaza A, Letunic I, Bork P, and Huerta-Cepas J. 2021. eggNOG-mapper v2: Functional Annotation, Orthology Assignments, and Domain Prediction at the Metagenomic Scale. Molecular Biology and Evolution. 38: 5825–5829.

Cetinkaya Y, Falk P, and Mayhall CG. 2000. Vancomycin-Resistant Enterococci. Clinical Microbiology Reviews. 13: 686–707.

Chewapreecha C et al. 2014. Comprehensive Identification of Single Nucleotide Polymorphisms Associated with Beta-lactam Resistance within Pneumococcal Mosaic Genes. PLOS Genetics. 10: e1004547.

Cingolani P, Platts A, Wang LL, Coon M, Nguyen T, Wang L, Land SJ, Lu X, and Ruden DM. 2012. A program for annotating and predicting the effects of single nucleotide polymorphisms, SnpEff: SNPs in the genome of Drosophila melanogaster strain w1118; iso-2; iso-3. Fly. 6: 80–92.

Collins C and Didelot X. 2018. A phylogenetic method to perform genome-wide association studies in microbes that accounts for population structure andrecombination. PLOS Computational Biology. 14: e1005958.

Consortium TC. 2022. Genome-wide association studies of global Mycobacterium tuberculosis resistance to 13 antimicrobials in 10,228 genomes identify new resistance mechanisms. PLOS Biology. 20: e3001755.

Cornick JE, Harris SR, Parry CM, Moore MJ, Jassi C, Kamng’ona A, Kulohoma B, Heyderman RS, Bentley SD, and Everett DB. 2014. Genomic identification of a novel co-trimoxazole resistance genotype and its prevalence amongst Streptococcus pneumoniae in Malawi. Journal of Antimicrobial Chemotherapy. 69: 368–374.

Crawford L, Flaxman SR, Runcie DE, and West M. 2019. VARIABLE PRIORITIZATION IN NONLINEAR BLACK BOX METHODS: A GENETIC ASSOCIATION CASE STUDY. The annals of applied statistics. 13: 958–989.

Croucher NJ, Finkelstein JA, Pelton SI, Mitchell PK, Lee GM, Parkhill J, Bentley SD, Hanage WP, and Lipsitch M. 2013. Population genomics of post-vaccine changes in pneumococcal epidemiology. Nature Genetics. 45: 656–663.

Croucher NJ, Finkelstein JA, Pelton SI, Parkhill J, Bentley SD, Lipsitch M, and Hanage WP. 2015. Population genomic datasets describing the post-vaccine evolutionary epidemiology of Streptococcus pneumoniae. Scientific Data. 2: 150058.

Danecek P et al. 2021. Twelve years of SAMtools and BCFtools. GigaScience. 10: giab008.

Derelle R, Lees J, Phelan J, Lalvani A, Arinaminpathy N, and Chindelevitch L. 2023. fastlin: an ultra-fast program for Mycobacterium tuberculosis complex lineage typing. Bioinformatics. 39: btad648.

Dias R, Félix D, Caniça M, and Trombe MC. 2009. The highly conserved serine threonine kinase StkP of Streptococcus pneumoniae contributes to penicillin susceptibility independently from genes encoding penicillin-binding proteins. BMC Microbiology. 9: 121.

Dickens E et al. 2026. A comprehensive AMR genotype-phenotype database (CABBAGE). en. ISSN: 2692-8205 Pages: 2025.11.12.688105 Section: New Results.

Earle SG et al. 2016. Identifying lineage effects when controlling for population structure improves power in bacterial association studies. Nature Microbiology. 1: 1–8.

Eyles S. 2004. Mycobacterium tuberculosis pks12 Produces a Novel Polyketide Presented by CD1c to T Cells. Journal of Experimental Medicine.

Eyre DW et al. 2017. WGStopredict antibiotic MICsfor Neisseriagonorrhoeae. Journal of Antimicrobial Chemotherapy. 72: 1937–1947.

Farhat MR et al. 2019. GWAS for quantitative resistance phenotypes in Mycobacterium tuberculosis reveals resistance genes and regulatory regions. Nature Communications. 10: 2128.

Fleurie A et al. 2014. MapZ marks the division sites and positions FtsZ rings in Streptococcus pneumoniae. Nature. 516: 259–262.

Fullerton AS and Xu J. 2012. The proportional odds with partial proportionality constraints model for ordinal response variables. Social Science Research. 41: 182–198.

Garde S, Chodisetti PK, and Reddy M. 2026. Peptidoglycan: Structure, Synthesis, and Regulation. EcoSal Plus. 9: eESP–0010–2020.

Gart JJ and Zweifel JR. 1967. On the Bias of Various Estimators of the Logit and Its Variance with Application to Quantal Bioassay.

Gelman A. 2008. Scaling regression inputs by dividing by two standard deviations. Statistics in Medicine. 27: 2865–2873.

Gibson PS, Bexkens E, Zuber S, Cowley LA, and Veening JW. 2022. The acquisition of clinically relevantamoxicillin resistance in Streptococcus pneumoniae requires ordered horizontal gene transfer of four loci. PLOS Pathogens. 18: e1010727.

Haasum Y, Ström K, Wehelie R, Luna V, Roberts MC, Maskell JP, Hall LMC, and Swedberg G. 2001. Amino Acid Repetitions in the Dihydropteroate Synthase of Streptococcus pneumoniae Lead to Sulfonamide Resistance with Limited Effects on SubstrateKm. Antimicrobial Agents and Chemotherapy. 45: 805–809.

Hoffman GE. 2013. Correcting for population structure and kinship using the linear mixed model: theory and extensions. PloS One. 8: e75707.

Huerta-Cepas J et al. 2019. eggNOG 5.0: a hierarchical, functionally and phylogenetically annotated orthology resource based on 5090 organisms and 2502 viruses. Nucleic Acids Research. 47: D309–D314.

Kim JI, Maguire F, Tsang KK, Gouliouris T, Peacock SJ, McAllister TA, McArthur AG, and Beiko RG. 2022. Machine Learning for Antimicrobial Resistance Prediction: Current Practice, Limitations, and Clinical Perspective. Clinical Microbiology Reviews. 35: e00179–21.

Kulkarni SG, Laurent S, Miotto P, Walker TM, Chindelevitch L, Nathanson CM, Ismail N, Rodwell TC, and Farhat MR. 2025. Multivariable regression models improve accuracy and sensitive grading of antibiotic resistance mutations in Mycobacterium tuberculosis. Nature Communications. 16: 2149.

Land AD, Tsui HCT, Kocaoglu O, Vella SA, Shaw SL, Keen SK, Sham LT, Carlson EE, and Winkler ME. 2013. Requirement of essential Pbp2x and GpsB for septal ring closure in Streptococcus pneumoniae D39. Molecular Microbiology. 90: 939–955.

Lees JA, Galardini M, Bentley SD, Weiser JN, and Corander J. 2018. pyseer: a comprehensive tool for microbial pangenome-wide association studies. Bioinformatics. 34: 4310–4312.

Lees JA, Harris SR, Tonkin-Hill G, Gladstone RA, Lo SW, Weiser JN, Corander J, Bentley SD, and Croucher NJ. 2019. Fast and flexible bacterial genomic epidemiology with PopPUNK. Genome Research. 29: 304–316.

Lees JA, Mai TT, Galardini M, Wheeler NE, Horsfield ST, Parkhill J, and Corander J. 2020. Improved Prediction of Bacterial Genotype-Phenotype Associations Using Interpretable Pangenome-Spanning Regressions. mBio. 11: e01344–20.

Lempens P, Meehan CJ, Vandelannoote K, Fissette K, Rijk P de, Van Deun A, Rigouts L, and Jong BC de. 2018. Isoniazid resistance levels of Mycobacterium tuberculosis can largely be predicted by high-confidence resistance-conferring mutations. Scientific Reports. 8: 3246.

Lipworth S, Crook D, Walker AS, Peto T, and Stoesser N. 2024. Exploring uncatalogued genetic variation in antimicrobial resistance gene families in Escherichia coli: an observational analysis. The Lancet Microbe. 5:

Ma KC, Mortimer TD, Duckett MA, Hicks AL, Wheeler NE, Sánchez-Busó L, and Grad YH. 2020. Increased power from conditional bacterial genome-wide association identifies macrolide resistance mutations in Neisseria gonorrhoeae. Nature Communications. 11: 5374.

Ma P, Luo T, Ge L, Chen Z, Wang X, Zhao R, Liao W, and Bao L. 2021. Compensatory effects of M. tuberculosis rpoB mutations outside the rifampicin resistance-determining region. Emerging Microbes & Infections. 10: 743–752.

Mai TT, Turner P, and Corander J. 2021. Boosting heritability: estimating the genetic component of phenotypic variation with multiple sample splitting. BMC Bioinformatics. 22: 164.

Mallawaarachchi S, Tonkin-Hill G, Croucher NJ, Turner P, Speed D, Corander J, and Balding D. 2022. Genomewide association, prediction and heritability in bacteria with application to Streptococcus pneumoniae. NAR Genomics and Bioinformatics. 4: lqac011.

Mascher T, Heintz M, Zähner D, Merai M, and Hakenbeck R. 2006. The CiaRH System of Streptococcus pneumoniae Prevents Lysis during Stress Induced by Treatment with Cell Wall Inhibitors and by Mutations in pbp2x Involved in *β*-lactam Resistance. Journal of Bacteriology. 188: 1959–1968.

Maskell JP, Sefton AM, and Hall LM. 1997. Mechanism of sulfonamide resistance in clinical isolates of Streptococcus pneumoniae. Antimicrobial Agents and Chemotherapy. 41: 2121–2126.

Maskell JP, Sefton AM, and Hall LMC. 2001. Multiple Mutations Modulate the Function of Dihydrofolate Reductase in Trimethoprim-ResistantStreptococcus pneumoniae. Antimicrobial Agents and Chemotherapy. 45: 1104–1108.

Massidda O, Anderluzzi D, Friedli L, and Feger G. 1998. Unconventional organization of the division and cell wall gene cluster of Streptococcus pneumoniae. Microbiology. 144 (Pt 11): 3069–3078.

Mechanism of action of penicillins 2026. Mechanism of action of penicillins: a proposal based on their structural similarity to acyl-D-alanyl-D-alanine. en.

Min S et al. 2015. Frequencyof Spontaneous Resistanceto Peptide Deformylase Inhibitor GSK1322322 in Haemophilus influenzae, Staphylococcus aureus, Streptococcus pyogenes, and Streptococcus pneumoniae. Antimicrobial Agents and Chemotherapy. 59: 4644–4652.

Morlot C, Bayle L, Jacq M, Fleurie A, Tourcier G, Galisson F, Vernet T, Grangeasse C, and Di Guilmi AM. 2013. Interaction of Penicillin-Binding Protein 2x and Ser/Thr protein kinase StkP, two key players in Streptococcus pneumoniae R6 morphogenesis. Molecular Microbiology. 90: 88–102.

Moure R, Español M, Tudó G, Vicente E, Coll P, Gonzalez-Martin J, Mick V, Salvadó M, and Alcaide F. 2014. Characterization of the embB gene in Mycobacterium tuberculosis isolates from Barcelona and rapid detection of main mutations related to ethambutol resistance using a low-density DNA array. Journal of Antimicrobial Chemotherapy. 69: 947–954.

Napier G, Campino S, Phelan JE, and Clark TG. 2023. Large-scale genomic analysis of Mycobacterium tuberculosis reveals extent of target and compensatory mutations linked to multi-drug resistant tuberculosis. Scientific Reports. 13: 623.

Nirwal S, Czarnocki-Cieciura M, Chaudhary A, Zajko W, Skowronek K, Chamera S, Figiel M, and Nowotny M. 2023. Mechanism of RecF–RecO–RecR cooperation in bacterial homologous recombination. Nature Structural & Molecular Biology. 30: 650–660.

Nishimoto AT, Dao TH, Jia Q, Ortiz-Marquez JC, Echlin H, Vogel P, Opijnen Tv, and Rosch JW. 2022. Interspecies recombination, not de novo mutation, maintains virulence after-lactam resistance acquisition in Streptococcus pneumoniae. Cell Reports. 41:

Pang Y, Zhu D, Zheng H, Shen J, Hu Y, Liu J, and Zhao Y. 2017. Prevalence and molecular characterization of pyrazinamide resistance among multidrug-resistant Mycobacterium tuberculosis isolates from Southern China. BMC Infectious Diseases. 17: 711.

Pereira AR, Reed P, Veiga H, and Pinho MG. 2013. The Holliday junction resolvase RecU is required for chromosome segregation and DNA damage repair in Staphylococcus aureus. BMC Microbiology. 13: 18.

Périchon B and Courvalin P. 2009. VanA-Type Vancomycin-Resistant Staphylococcus aureus. Antimicrobial Agents and Chemotherapy. 53: 4580–4587.

Piironen J and Vehtari A. 2017. Sparsity information and regularization in the horseshoe and other shrinkage priors. Electronic Journal of Statistics. 11:

Pikis A, Donkersloot JA, Rodriguez WJ, and Keith JM. 1998. AConservative Amino Acid Mutationinthe Chromosome-Encoded Dihydrofolate Reductase Confers Trimethoprim Resistance in Streptococcus pneumoniae. The Journal of Infectious Diseases. 178: 700–706.

Power RA, Parkhill J, and Oliveira T de. 2017. Microbial genome-wide associationstudies: lessons fromhuman GWAS. Nature Reviews Genetics. 18: 41–50.

Praski Alzrigat L, Huseby DL, Brandis G, and Hughes D. 2017. Fitness cost constrains the spectrum of marR mutations in ciprofloxacin-resistant Escherichia coli. Journal of Antimicrobial Chemotherapy. 72: 3016–3024.

Prunier J, Lemaçon A, Bastien A, Jafarikia M, Porth I, Robert C, and Droit A. 2019. LD-annot: A Bioinformatics Tool to Automatically Provide Candidate SNPs With Annotations for Genetically Linked Genes. Frontiers in Genetics. 10:

Robicsek A, Jacoby GA, and Hooper DC. 2006. The worldwide emergence of plasmid-mediated quinolone resistance. The Lancet Infectious Diseases. 6: 629–640.

Ruden DM, Cingolani P, Patel VM, Coon M, Nguyen T, Land SJ, and Lu X. 2012. Using Drosophila melanogaster as a Model for Genotoxic Chemical Mutational Studies with a New Program, SnpSift. Frontiers in Genetics. 3:

Rued BE et al. 2017. Suppression and Synthetic-Lethal Genetic Relationships of ΔgpsB Mutations Indicate That GpsB Mediates Protein Phosphorylation and Penicillin-Binding Protein Interactions in Streptococcus pneumoniae D39. Molecular microbiology. 103: 931–957.

Safari D, Putri HFM, Bimantari A, Paramaiswari WT, Tafroji W, Khoeri MM, and Salsabila K. 2021. Genetic characterization of co-trimoxazole non-susceptible Streptococcus pneumoniae isolates from Indonesia. Access Microbiology. 3:

Skwark MJ et al. 2017. Interacting networks of resistance, virulence and core machinery genes identified by genome-wide epistasis analysis. PLOS Genetics. 13: e1006508.

Tran TDH, Kwon HY, Kim EH, Kim KW, Briles DE, Pyo S, and Rhee DK. 2011. Decrease in Penicillin Susceptibility Due to Heat Shock Protein ClpL in Streptococcus pneumoniae. Antimicrobial Agents and Chemotherapy. 55: 2714–2728.

Trauner A et al. 2021. Expression Dysregulation as a Mediator of Fitness Costs in Antibiotic Resistance. Antimicrobial Agents and Chemotherapy. 65: 10.1128/aac.00504-21.

VanRaden PM. 2008. Efficient Methods to Compute Genomic Predictions. Journal of Dairy Science. 91: 4414–4423.

Xu Z, Zhou A, Wu J, Zhou A, Li J, Zhang S, Wu W, Karakousis PC, and Yao YF. 2018. Transcriptional Approach for Decodingthe MechanismofrpoCCompensatory Mutationsforthe Fitness Costin Rifampicin-Resistant Mycobacterium tuberculosis. Frontiers in Microbiology. 9: 2895.

Zhou M, Wang L, Wang Z, Kudinha T, Wang Y, Xu Y, and Liu Z. 2022. Molecular Characterization of Penicillin-Binding Protein2x, 2b and 1a of Streptococcus pneumoniae Causing Invasive Pneumococcal Diseases in China: A Multicenter Study. Frontiers in Microbiology. 13:

Zuo TY, Liu QY, Gan MY, and Gao Q. 2018. [Comprehensive identification of compensatory mutations in rifampicin-resistant Mycobacterium tuberculosis strains]. Zhonghua jie he he hu xi za zhi = Zhonghua jiehe he huxi zazhi = Chinese journal of tuberculosis and respiratory diseases. 41: 207–212.

