## Supplementary Figures and Tables for "Bayesian bacterial GWAS model reveals threshold-dependent genetic changes in antimicrobial resistance phenotypes"

**Supplementary Table ST1. Notation used in the four Bayesian GWAS models.** Bold lower-case symbols are vectors and bold upper-case symbols matrices; $\mathcal{N}, \mathcal{N}^+$ and $\mathcal{C}^+$ denote normal, half-normal and half-Cauchy distributions. A tilde denotes a posterior median (e.g. $\tilde{\beta}$).

| Symbol | Definition | Eq. |
|---|---|---|
| **Indices and dimensions** | | |
| $N, n$ | Number of isolates in the dataset, and the index of an isolate | – |
| $V, v$ | Number of variants fitted (after MAF filtering and LD pruning), and the index of a variant | – |
| $K, k$ | Number of ordered MIC categories after discretization, and the index of a cutpoint, $k \leq K-1$ | – |
| $L, \ell; S, s$ | Lineage clusters and lineage subclusters, and their indices | – |
| $\pi(s), \mathcal{S}_\ell$ | Parent lineage of subcluster $s$; set of subclusters nested within lineage $\ell$ | 11, 12 |
| $n^{\text{ref}}, n_0^{\text{ref}}, n_1^{\text{ref}}$ | Isolates in the reference (lowest-mean-phenotype) subcluster, and those susceptible (logistic) or in MIC category 1 (ordinal) | 16, 19 |
| $N_{\text{test}}$ | Held-out isolates on which predictive metrics are computed | – |
| **Observed data** | | |
| $y_n, \hat{y}_n$ | Observed and posterior-predictive phenotype of isolate $n$: binary, ordered category, or $\log_2$ MIC | – |
| $x_{n,v}; \bar{x}_v, \text{sd}(x_v)$ | Unstandardized binary genotype of isolate $n$ at variant $v$; column mean and standard deviation | 1 |
| $x_{n,v}^{\text{std}}, \mathbf{X}^{\text{std}}, \mathbf{x}_n$ | Standardized genotype, the $N \times V$ standardized genotype matrix, and its $n$-th row (invariant columns pinned to 0) | 1 |
| $\mathbf{X}_{\text{sub}}, \mathbf{s}_n$ | $N \times (S-1)$ subcluster design matrix under treatment contrasts, and its $n$-th row | 12 |
| $\boldsymbol{\kappa}, \kappa_k$ | $\log_2$ MIC breakpoint grid and its $k$-th breakpoint, setting the spacing of the cutpoint anchors | 20 |
| $\mathbf{K}, \mathbf{L}_K$ | Haploid VanRaden GRM (unit mean diagonal, $10^{-6}$ jitter) and its lower Cholesky factor | 13 |
| **Location parameters** | | |
| $\alpha, m_\alpha$ | Intercept of the logistic and continuous models and its data-informed anchor; in ordinal models, $m_\alpha$ is the level shift applied to the breakpoint grid | 17, 15, 20 |
| $\hat{p}_0$ | Laplace-smoothed fraction of reference-subcluster isolates that are susceptible (logistic) or in MIC category 1 (ordinal), clamped to $[0.5, 0.995]$ | 16, 19 |
| $c_k, m_{c,k}$ | $k$-th ordered cutpoint on the logistic latent scale and its prior anchor; the drift $c_k - m_{c,k}$ identifies uninformative breakpoints | 18, 21 |
| $\mu_n, \sigma$ | Conditional mean $\log_2$ MIC of isolate $n$ and the residual standard deviation (continuous model) | 24 |
| **Variant effects and the regularized horseshoe prior** | | |
| $\beta_v^{\text{std}}, \beta_v$ | Effect of variant $v$ on the standardized scale, and its backtransform $\beta_v = \beta_v^{\text{std}}/\text{sd}(x_v)$ | 1 |
| $\beta_{v,k}, \boldsymbol{\beta}_k$ | PPOM effect of variant $v$ at cutpoint $k$, unconstrained across cutpoints, and the coefficient vector at cutpoint $k$ | 23 |
| $\tilde{\beta}, \lvert\tilde{\beta}\rvert$ | Posterior median of a coefficient and its magnitude | – |
| $\exp(\beta_v), \exp(\beta_{v,k})$ | Per-allele (cumulative) odds ratio: constant across MIC thresholds in the POM, threshold-specific in the PPOM | – |
| $z_v, z_{v,k}$ | Standard normal auxiliaries in the non-centred horseshoe parameterization | 3, 8 |

*continued on next page*

*Table ST1 continued*

| Symbol | Definition | Eq. |
|---|---|---|
| $\lambda_v, \tilde{\lambda}_v$ | Local shrinkage scales, $\mathcal{C}^+(0, 2)$, and their regularized counterparts (per variant, or per variant–cutpoint in the PPOM) | 2, 7 |
| $\tau, \tau_0$ | Global shrinkage scale, $\mathcal{C}^+(0, \tau_0)$ with $\tau_0 = 1$ fixed; shared across all $V(K-1)$ PPOM coefficients | 5 |
| $c^2_{\text{slab}}; \nu, s$ | Slab variance capping the effective prior variance of non-zero coefficients, with degrees of freedom $\nu = 4$ and scale $s = 5$ ($s$ here is the slab scale, not the subcluster index) | 6 |
| **Population structure** | | |
| $\beta^{\text{lin}}_\ell$ | Lineage cluster effect, $\mathcal{N}(0, 0.1^2)$; absorbs the lineage-level mean | 10 |
| $\tilde{\beta}^{\text{sub}}_s, \beta^{\text{sub}}_s$ | Subcluster effect pooled toward its parent lineage, before and after within-lineage re-centring to sum to zero | 11, 12 |
| $\boldsymbol{\omega}, \eta^{\text{sub}}_n$ | Vector of re-centred subcluster coefficients (proportional across cutpoints), and the lineage contribution $\mathbf{s}_n\boldsymbol{\omega}$ | 22 |
| $\sigma_{\text{sub}}, z_s$ | Shared subcluster scale, $\mathcal{N}^+(0, 0.1^2)$, and its standard normal auxiliary | 11 |
| $\mathbf{u}, \sigma_g$ | Per-isolate GRM random effect replacing $\mathbf{s}_n\boldsymbol{\omega}$ in the comparison model, and its scale | 13 |
| **Heritability** | | |
| $V_A, V_{\text{pop}}, V_E$ | Additive genetic, population-structure and residual variance ($V_E = \sigma^2$ continuous, $\pi^2/3$ on the latent scale) | 26–28 |
| $h^2_{\text{n}}, h^2_{\text{b}}$ | Narrow- and broad-sense heritability; computed per cutpoint in the PPOM, and a lower bound under shrinkage | 28, 29 |

| | Logistic | Proportional odds (POM) | Partial proportional odds (PPOM) | Continuous |
|---|---|---|---|---|
| Phenotype $y_n$ | Binary, $y_n \in \{0, 1\}$, thresholded at a clinical breakpoint | Ordered category, $y_n \in \{1, \ldots, K\}$, from binned MICs | Ordered category, $y_n \in \{1, \ldots, K\}$, from binned MICs | $\log_2$-transformed MIC, $y_n \in \mathbb{R}$ |
| Likelihood and link | Bernoulli, logit link (Equation 25) | Ordered logistic, cumulative logit (Equation 22) | $K - 1$ Bernoulli terms per sample, cumulative logit (Equation 23) | Gaussian, identity link (Equation 24) |
| Location parameters | Intercept $\alpha$, anchored on the susceptible fraction of the reference subcluster (Equation 17) | $K - 1$ ordered cutpoints $c_k$, anchored on the $\log_2$-MIC breakpoint grid (Equation 18) | $K - 1$ ordered cutpoints $c_k$, anchored on the $\log_2$-MIC breakpoint grid (Equation 18) | Intercept $\alpha$, anchored on the mean $\log_2$-MIC of the reference subcluster (Equation 15) |
| Variant coefficients | One per variant, $\beta_v$ | One per variant, $\beta_v$, shared across all $K - 1$ cutpoints | One per variant *per cutpoint*, $\beta_{v,k}$, unconstrained across cutpoints | One per variant, $\beta_v$ |
| Shrinkage prior | Regularized horseshoe on $\beta_v^{\text{std}}$ (Equation 2) | Regularized horseshoe on $\beta_v^{\text{std}}$ (Equation 2) | Independent horseshoe per (variant, cutpoint), with $\tau$ and $c^2_{\text{slab}}$ shared across all $V(K - 1)$ coefficients (Equation 7) | Regularized horseshoe on $\beta_v^{\text{std}}$ (Equation 2) |
| Interpretation of $\exp(\beta)$ | Odds ratio for resistance versus susceptibility at the breakpoint | Cumulative odds ratio, constrained to be constant across MIC thresholds | Cumulative odds ratio *at each* MIC threshold, permitting threshold-dependent effects | Not applicable; $\beta$ is a fold change in MIC in doubling dilutions |
| Population structure | Lineage and subcluster effects $\boldsymbol{\omega}$ | $\boldsymbol{\omega}$, proportional across cutpoints | $\boldsymbol{\omega}$, proportional across cutpoints (only variant effects are freed) | Lineage and subcluster effects $\boldsymbol{\omega}$ |
| Heritability | Single $h^2$, $V_E = \pi^2/3$ | Single $h^2$, $V_E = \pi^2/3$ | One $h^2$ per cutpoint, computed from $\boldsymbol{\beta}_k$ | Single $h^2$, $V_E = \sigma^2$ |
| RATE values | One per variant, summing to 1 across all variants | One per variant, summing to 1 across all variants | One per variant per cutpoint, summing to 1 at each cutpoint | One per variant, summing to 1 across all variants |
| Prediction | `bernoulli_logit_rng` | `ordered_logistic_rr` | Analytic $N_{\text{test}} \times K$ category probability matrix | `normal_rng` |
| Evaluation metrics | bACC, sensitivity, specificity, AUC, Brier, $F_1$, VME, ME | bACC, PPV, RPS, RPSS | bACC, PPV, RPS, RPSS | RMSE, MAE, $R^2$, CRPS, essential agreement |
| Intended use | A single clinically relevant breakpoint is of interest; discards dilution resolution and depends on the breakpoint applied | Ordering and censoring of the dilution series are retained; assumes each variant acts equally at every threshold | As POM, but recovers threshold-dependent, conditional and compensatory effects, at the cost of $V(K - 1)$ coefficients | Fine-grained dilution series; sensitive to left- and right-censoring of the tested range |

**Supplementary Table ST2.** Summary of the four Bayesian GWAS models, each of which has an inference and a prediction version. All four share the standardized genotype matrix, the regularized horseshoe prior on variant effects, and the hierarchical lineage cluster and subcluster correction for population structure; they differ in how the minimum inhibitory concentration phenotype is represented and in whether variant effects are permitted to vary across MIC thresholds.

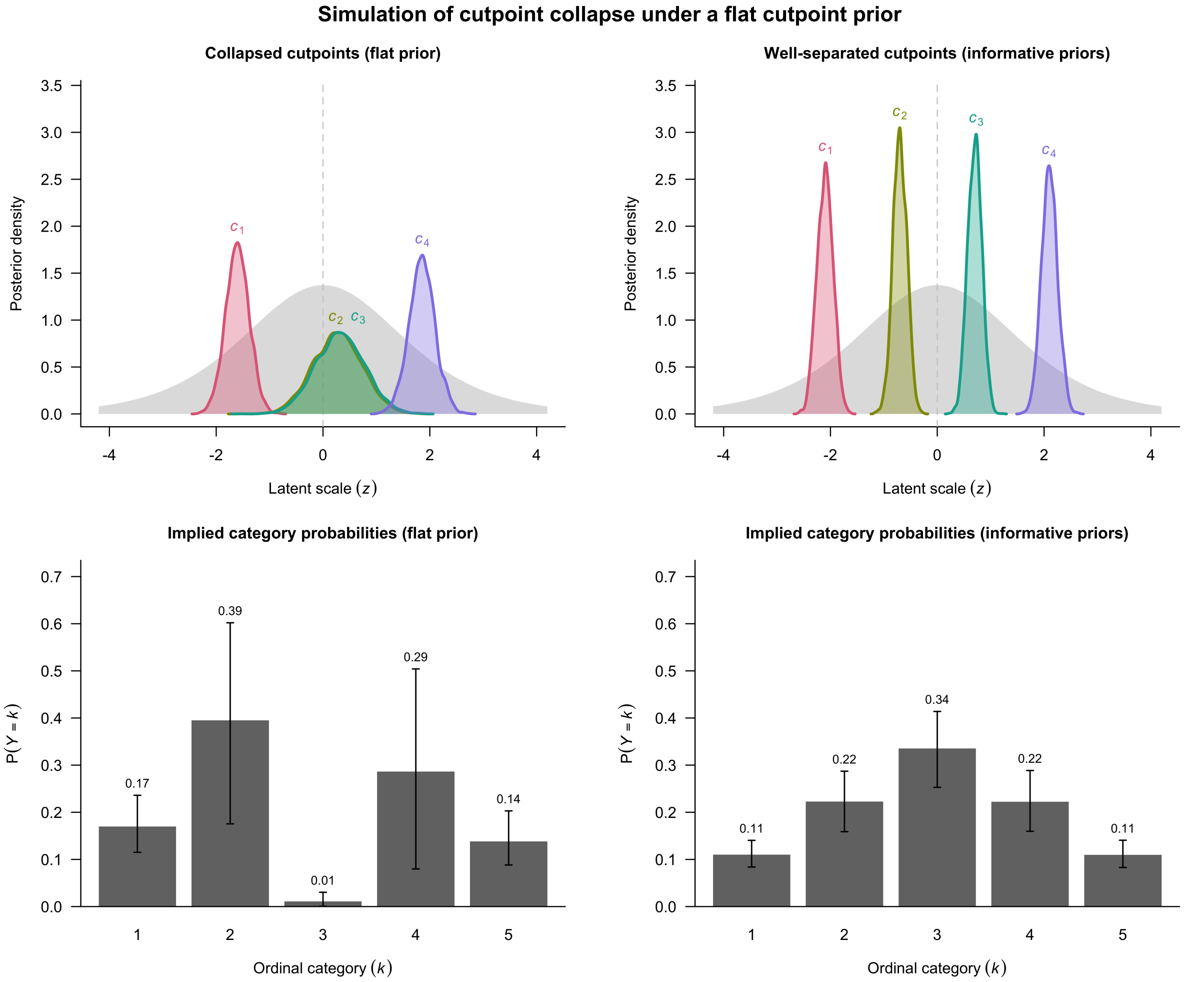


**Supplementary Figure S1.** Simulated cutpoint posterior distributions and their implied category probabilities showing how poor cutpoint separation renders some adjacent ordinal categories indistinguishable, a condition that may be ameliorated by informative priors. In this example, $4000$ draws were drawn from independent normal distributions for each of the four cutpoints. To simulate a collapse scenario, $c_2$ and $c_3$ were assigned nearly coincident locations and $c_2$ was given an inflated posterior standard deviation, whereas in the well-separated scenario all four cutpoints were placed at evenly spaced locations with small standard deviations. Implied category probabilities were then computed as $P(Y = k) = \text{logit}^{-1}(c_k) - \text{logit}^{-1}(c_{k-1})$, and summarized by their posterior means and $95\%$ credible intervals. The latent standard logistic (the assumed distribution of the continuous latent variable that underlies the ordinal response) is shown in gray.

| Variant effect | POM detection? | PPOM detection? |
|---|---|---|
| Variant causes the same change in log odds regardless of background resistance level (breakpoint chosen) | Yes | Yes |
| Variant causes different changes in log odds depending on the background resistance level (breakpoint chosen) | No (singular coefficient captures variable effects poorly) | Yes (multiple coefficients can describe variable effects well) |

**Supplementary Table ST3.** Comparison of variant effect detection in the proportional odds model (POM) and partial proportional odds model (PPOM).

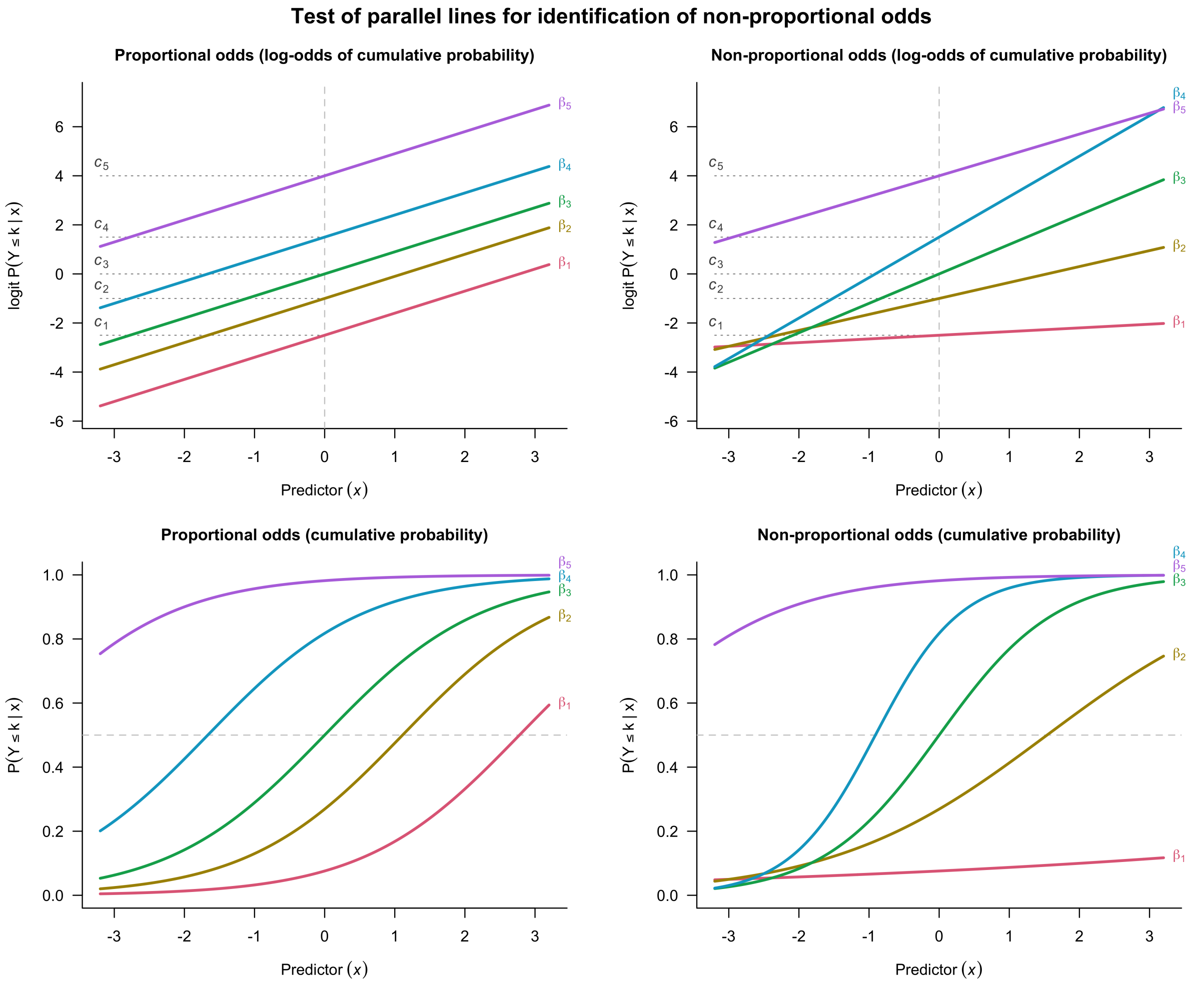


**Supplementary Figure S2.** Simulation of effect sizes $\beta_k$ for a genetic variant displaying proportional odds versus non-proportional odds, shown on both the log-odds and probability scale. Here, $\text{logit}[P(Y \leq k \mid x)] = c_k + \beta_k x$ for each category threshold $k$ in $1, 2, \ldots, K-1$, where $K$ is the number of categories (here, six). We see that a variant displaying proportional odds is defined by $\beta_1 = \cdots = \beta_{K-1} \equiv \beta$ for all $k$, resulting in parallel lines in the log-odds term $c_k + \beta_k x$, while intersections are observed for a variant displaying non-proportional odds due to differences in $\beta_k$.

| Pattern of coefficients across $K-1$ cutpoints | Interpretation | Example |
|---|---|---|
| All large, positive | At each cutpoint $c$, causes a large increase in the probability of a sample's phenotype being in any resistance category above $c$ compared to any resistance category below $c$ (proportional effect across all cutpoints) | ***Acquisition of resistance gene*** Acquisition of the *vanA* operon carried on the *Tn1546* transposon, which confers high-level vancomycin resistance in *Staphylococcus aureus* (Cetinkaya et al. 2000; Périchon and Courvalin 2009) |
| All small, positive | At each cutpoint $c$, causes a small increase in the probability of a sample's phenotype being in any resistance category above $c$ compared to any resistance category below $c$ (proportional effect across all cutpoints) | ***Variant causing change in efflux pump expression*** Single amino-acid substitutions in the *marR* repressor that reduce inhibition of the *marRAB* operon, causing upregulation of the *AcrAB-TolC* efflux pump and a subsequent small increase in ciprofloxacin resistance in *Escherichia coli* (Praski Alzrigat et al. 2017) |
| Only final coefficient ($\beta_{j,K-1}$) large, positive<br>Other coefficients near zero | Causes large increase in probability of a sample's phenotype being in the highest resistance category $K$, compared to any lower resistance category<br>Has no effect at other cutpoints | ***High-level resistance variant reliant on an epistatic interaction with a previously-acquired variant*** Acquisition of *parC* S87L or S87W, which do not confer increased ciprofloxacin resistance in isolation but result in 256-fold MIC increase relative to a reference strain when acquired alongside *gyrA* T83I in *E. coli* (Bruchmann et al. 2013) |
| Only first coefficient ($\beta_{j,1}$) large, positive<br>Other coefficients near zero | Causes large increase in probability of a sample's phenotype being in any resistance category higher than the lowest resistance category<br>Has no effect at higher cutpoints | ***Low-level resistance variant*** Acquisition of a *qnrA*-bearing plasmid, which confers a small increase in quinolone MIC in *E. coli* (Robicsek et al. 2006) |
| Mixed pattern | Complex, variable effect | ***Regulatory region variant*** |
| All near zero | Arises from insufficient evidence to support an effect on the phenotype or from strong evidence of no effect on the phenotype | ***Intergenic or neutral variant*** |

**Supplementary Table ST4.** Interpretation of the set of $K-1$ coefficients, $\{\beta_{j,1},\ldots,\beta_{j,K-1}\}$, for each variant $j$ in a partial proportional odds GWAS model with $K$ ordered categories.

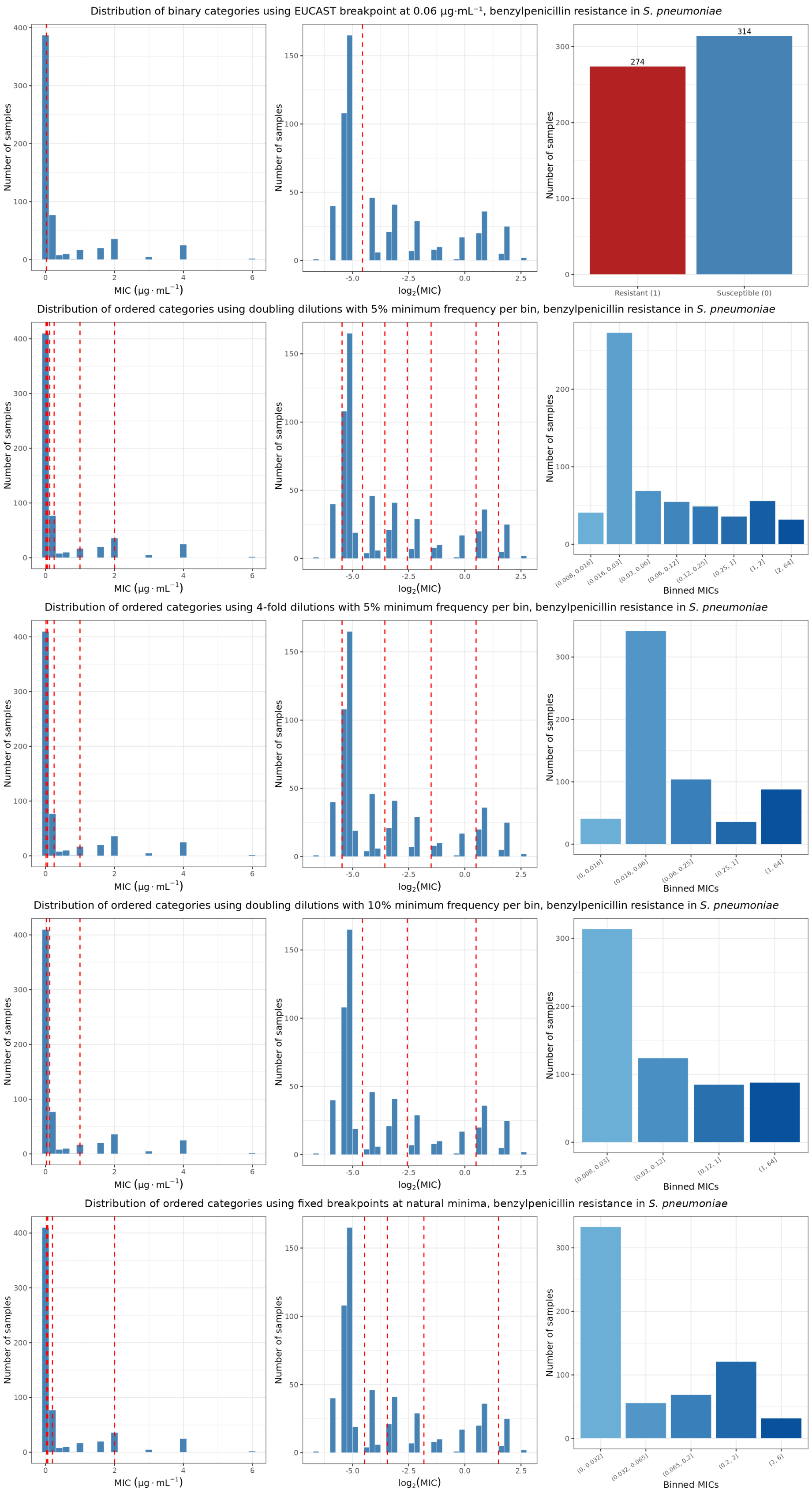


**Supplementary Figure S3.** Minimum inhibitory concentration (MIC) intervals or resistant-sensitive breakpoints chosen for *S. pneumoniae* with the benzylpenicillin resistance phenotype.

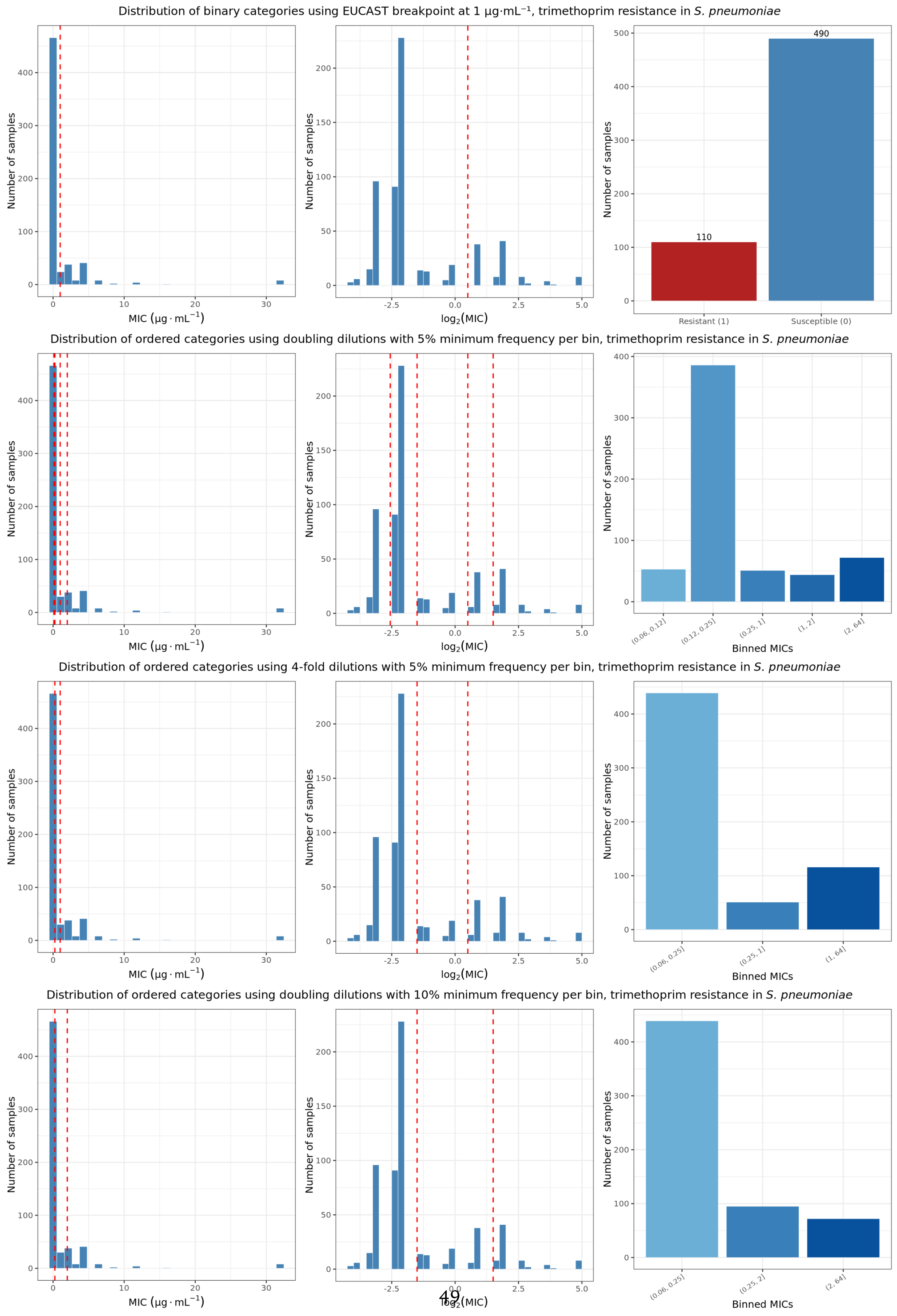


**Supplementary Figure S4.** Minimum inhibitory concentration (MIC) intervals or resistant-sensitive breakpoints chosen for *S. pneumoniae* with the trimethoprim resistance phenotype.

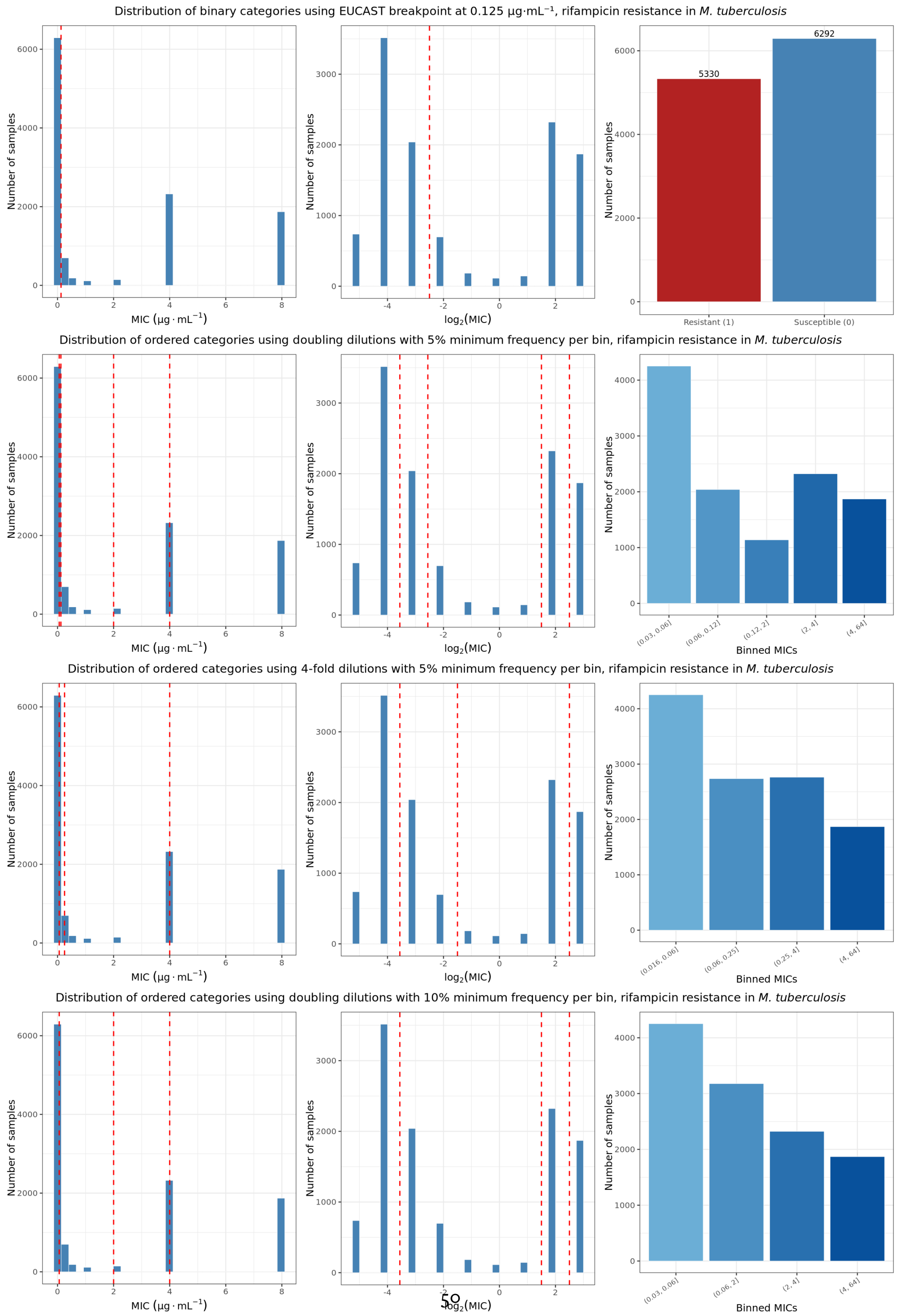




**Supplementary Figure S5.** Minimum inhibitory concentration (MIC) intervals or resistant-sensitive breakpoints chosen for *M. tuberculosis* with the rifampicin resistance phenotype.

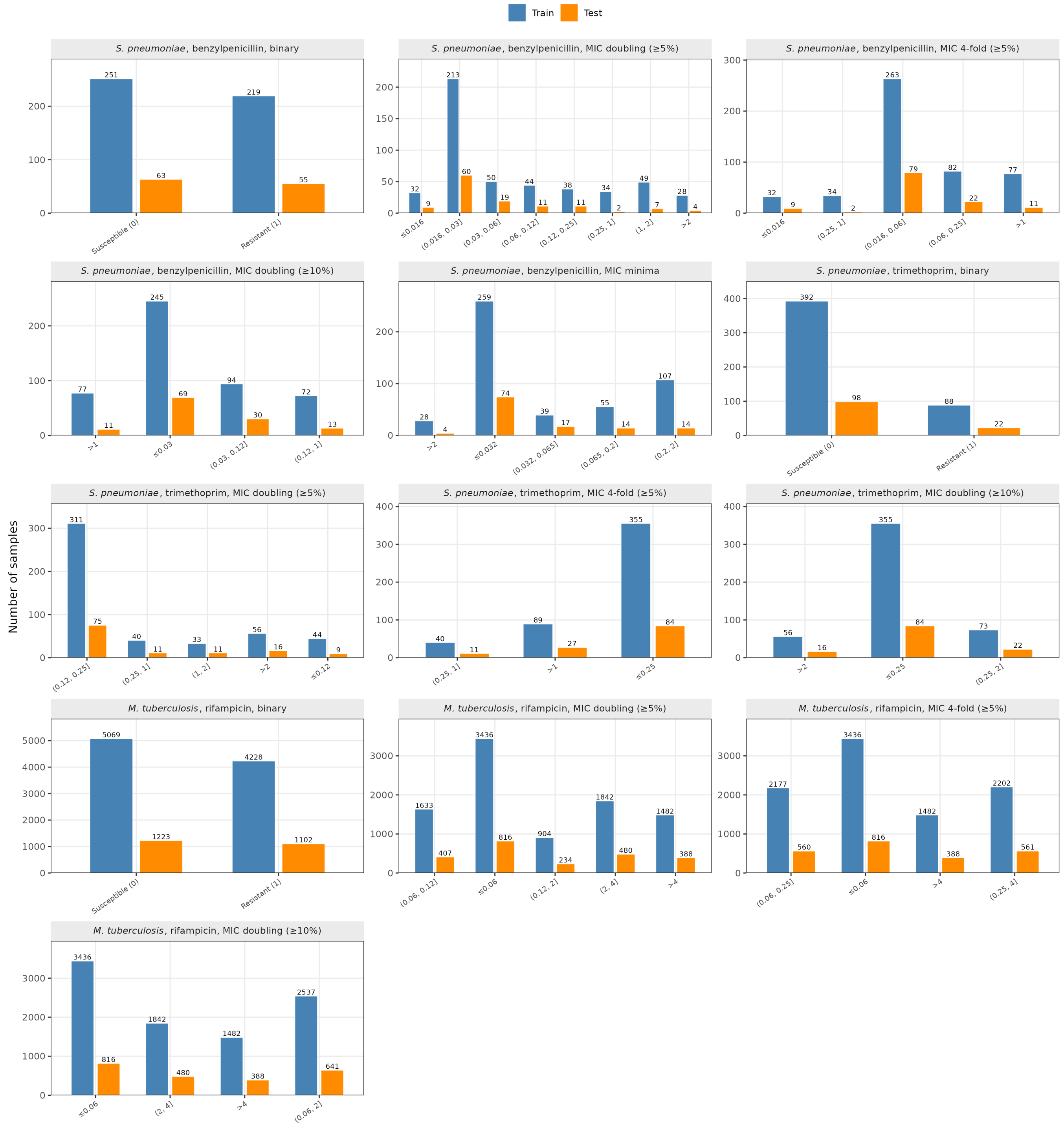


**Supplementary Figure S6.** Random 80:20 train:test splits used for each combination of species, AMR phenotype, and MIC binning strategy.

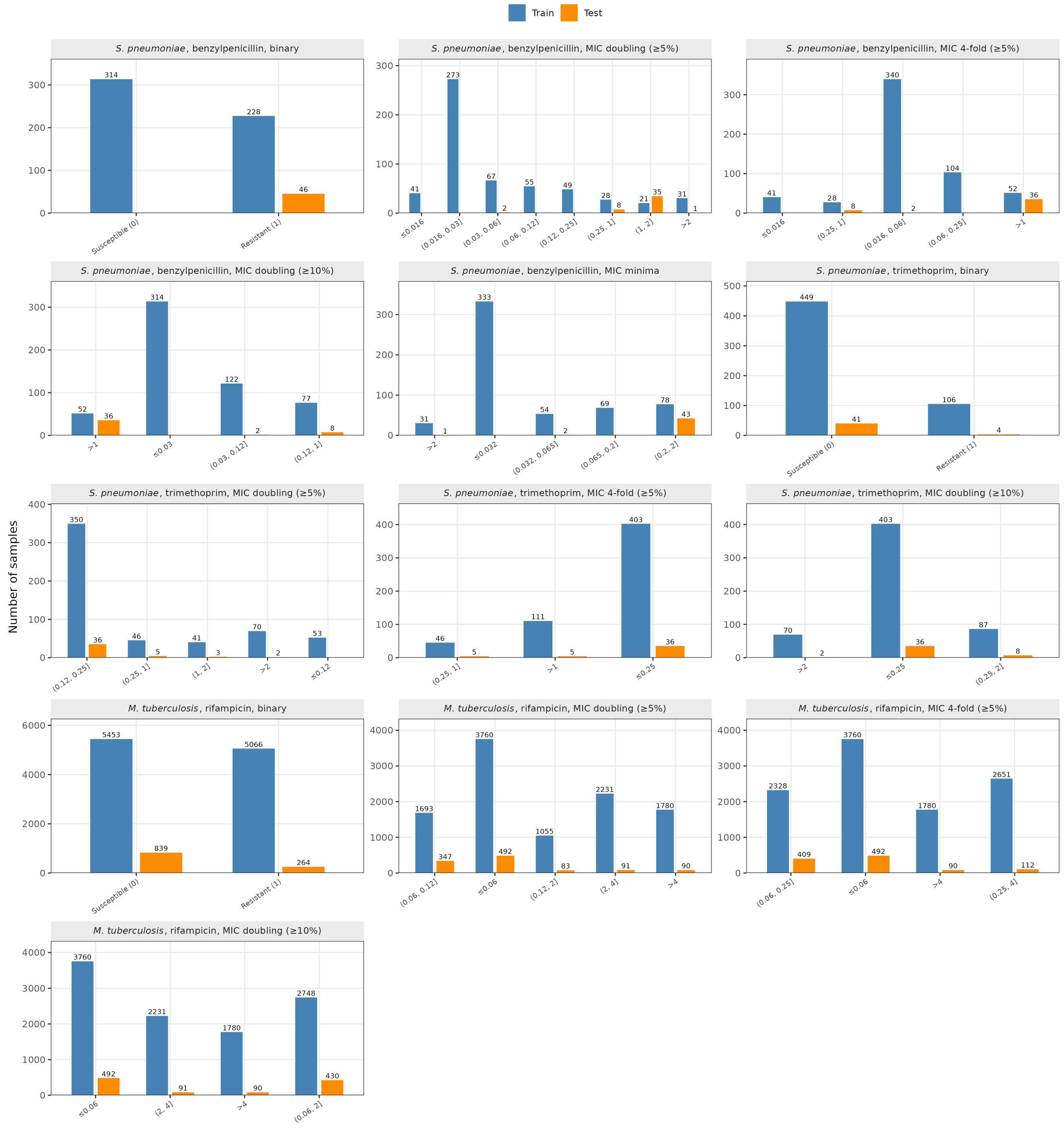


**Supplementary Figure S7.** Train:test splits in which all samples belonging to one lineage subcluster were withheld from the training dataset, shown for each combination of species, AMR phenotype, and MIC binning strategy.

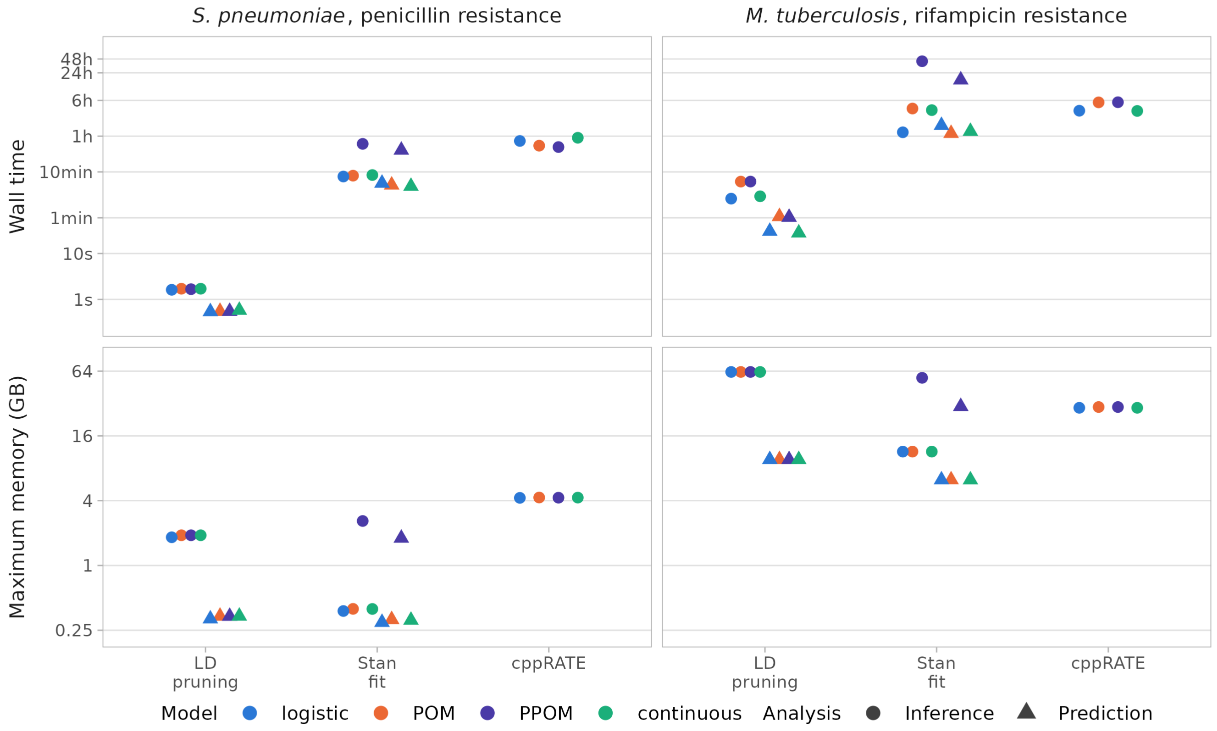


**Supplementary Figure S8.** Time and maximum memory consumption for LD pruning with BacPrune-Rust, Stan model fitting, and cppRATE using the GHOST workflow (Figure 2). The "minima" (*S. pneumoniae*, $K = 5$) and doubling dilution $\geq 10\%$ (*M. tuberculosis*, $K = 4$) discretizations were used for the ordered models. 32 or 48 CPUs were used, respectively (note that BacPrune-Rust is single-threaded). For PPOMs, the cppRATE value shown is the median across its $K - 1$ per-cutpoint runs. cppRATE was not run for prediction analyses. For inference, the *S. pneumoniae* runs contained 588 isolates (binary) or 611 isolates (continuous and ordered models) and 32,406 variants, and the *M. tuberculosis* runs contained 11,622 isolates and 75,272 variants; pruning retained 26,381, 26,512 and 40,941 variants respectively. For prediction, variants in perfect LD were pruned using the held-out test split rather than the training split, meaning the pruning scores were calculated on a fifth of the isolates (118/588, 123/611, and 2325/11,622), retaining 21,190, 21,746 and 26,551 variants.

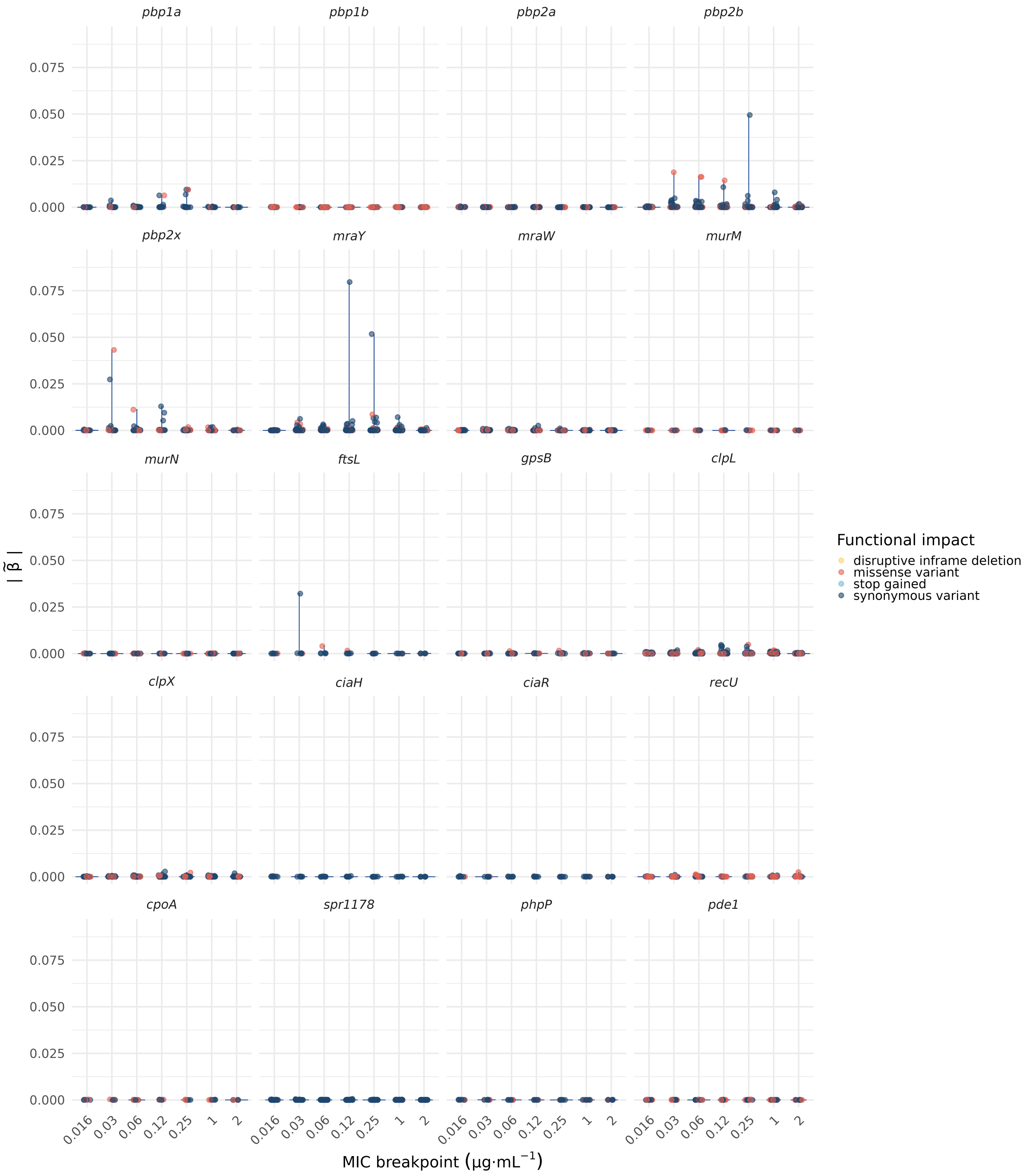


**Supplementary Figure S9.** Median coefficient magnitude, $|\tilde{\beta}|$, of all variants located within several key genes known to be associated with benzylpenicillin resistance in *S. pneumoniae* (*pbp2x*, *pbp2b*, *pbp1a*, and *mraY*), as well as those believed not to be involved in resistance (*pbp1b*, *pbp2a*), and others that have been associated with resistance in previous studies. Effects were fitted in a partial proportional odds model (PPOM) using MIC breakpoints calculated using the "standard" binning strategy, which uses every doubling dilution containing a minimum of 5% of isolate phenotypes (Supplementary Figure S3).

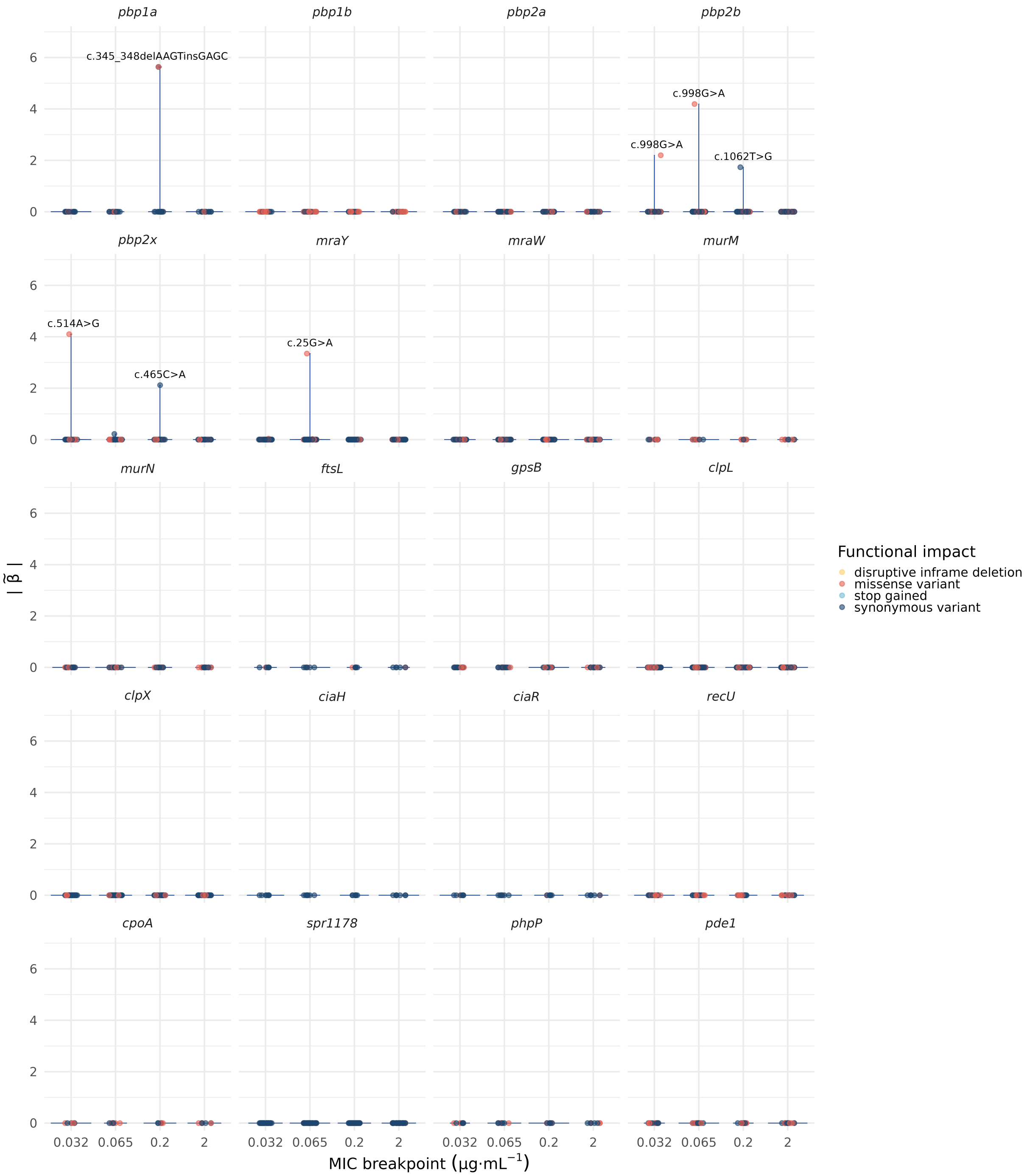


**Supplementary Figure S10.** Median coefficient magnitude, $|\tilde{\beta}|$, of all variants located within several key genes known to be associated with benzylpenicillin resistance in *S. pneumoniae* (*pbp2x*, *pbp2b*, *pbp1a*, and *mraY*), as well as those believed not to be involved in resistance (*pbp1b*, *pbp2a*), and others that have been associated with resistance in previous studies. Effects were fitted in a partial proportional odds model (PPOM) using MIC breakpoints calculated using the “minima” binning strategy (Supplementary Figure S3).

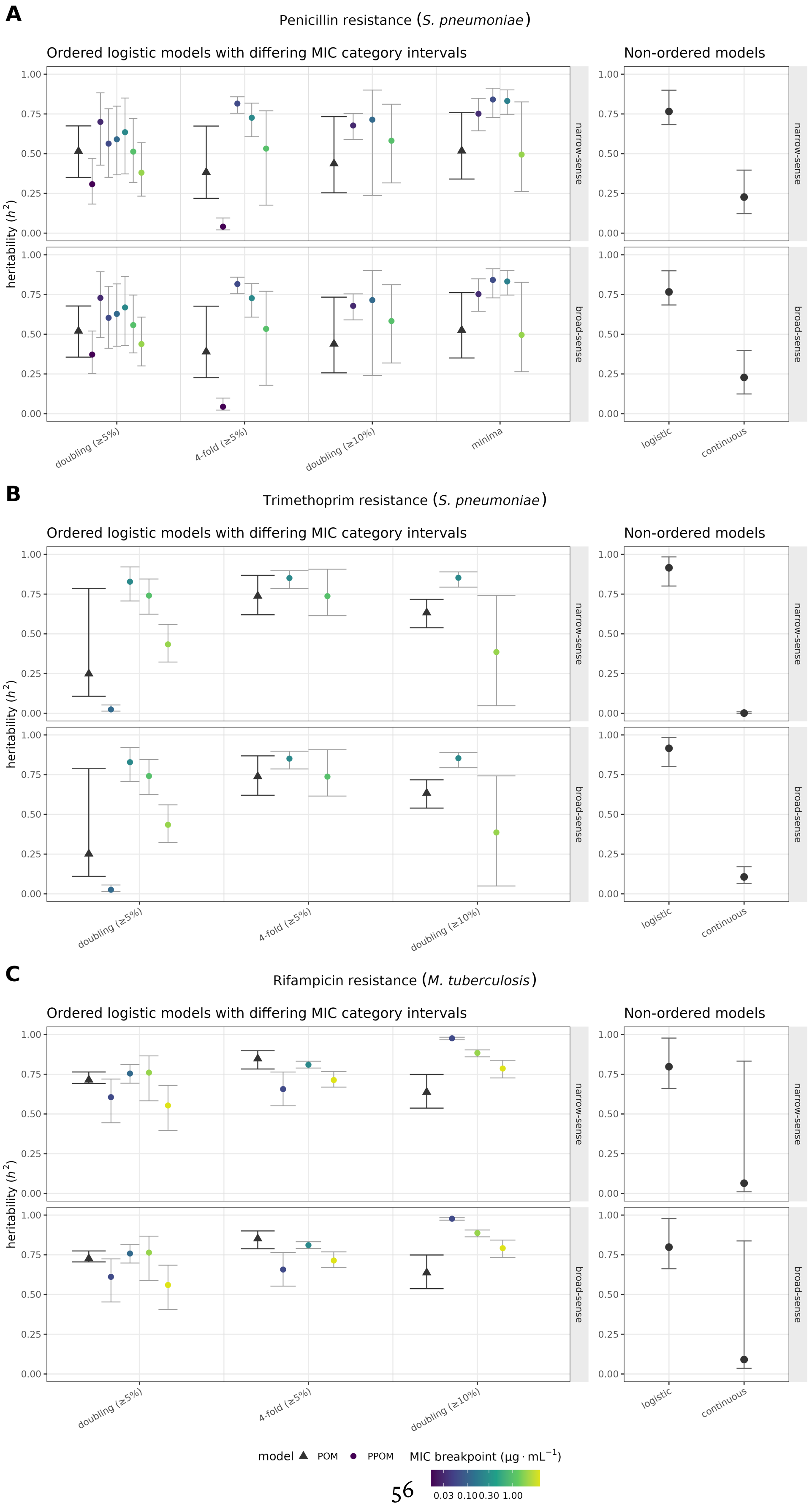


**Supplementary Figure S11.** 95% credible interval of broad- and narrow-sense heritability estimates across different phenotype discretization strategies. (A) *S. pneumoniae* with penicillin resistance phenotype. (B) *S. pneumoniae* with trimethoprim resistance phenotype. (C) *M. tuberculosis* with rifampicin resistance phenotype.

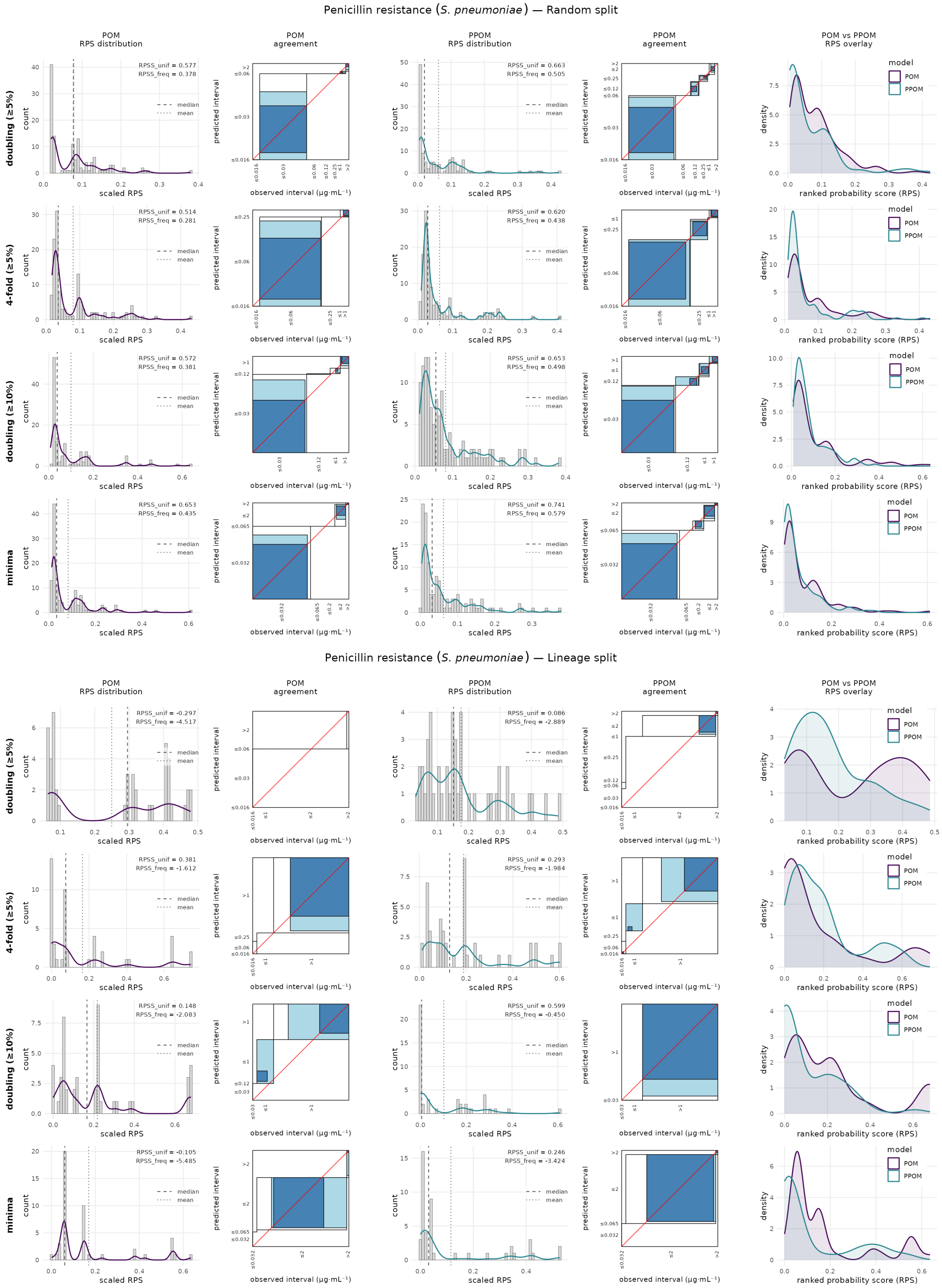


**Supplementary Figure S12.** Density plots of prediction errors as measured by RPS and RPSS, and agreement plots of predicted MIC versus observed phenotype, for POM and PPOM fit to *S. pneumoniae* with a penicillin resistance phenotype with different MIC discretization strategies. Top: Random train:test split. Bottom: one lineage subcluster withheld from training and used for prediction.

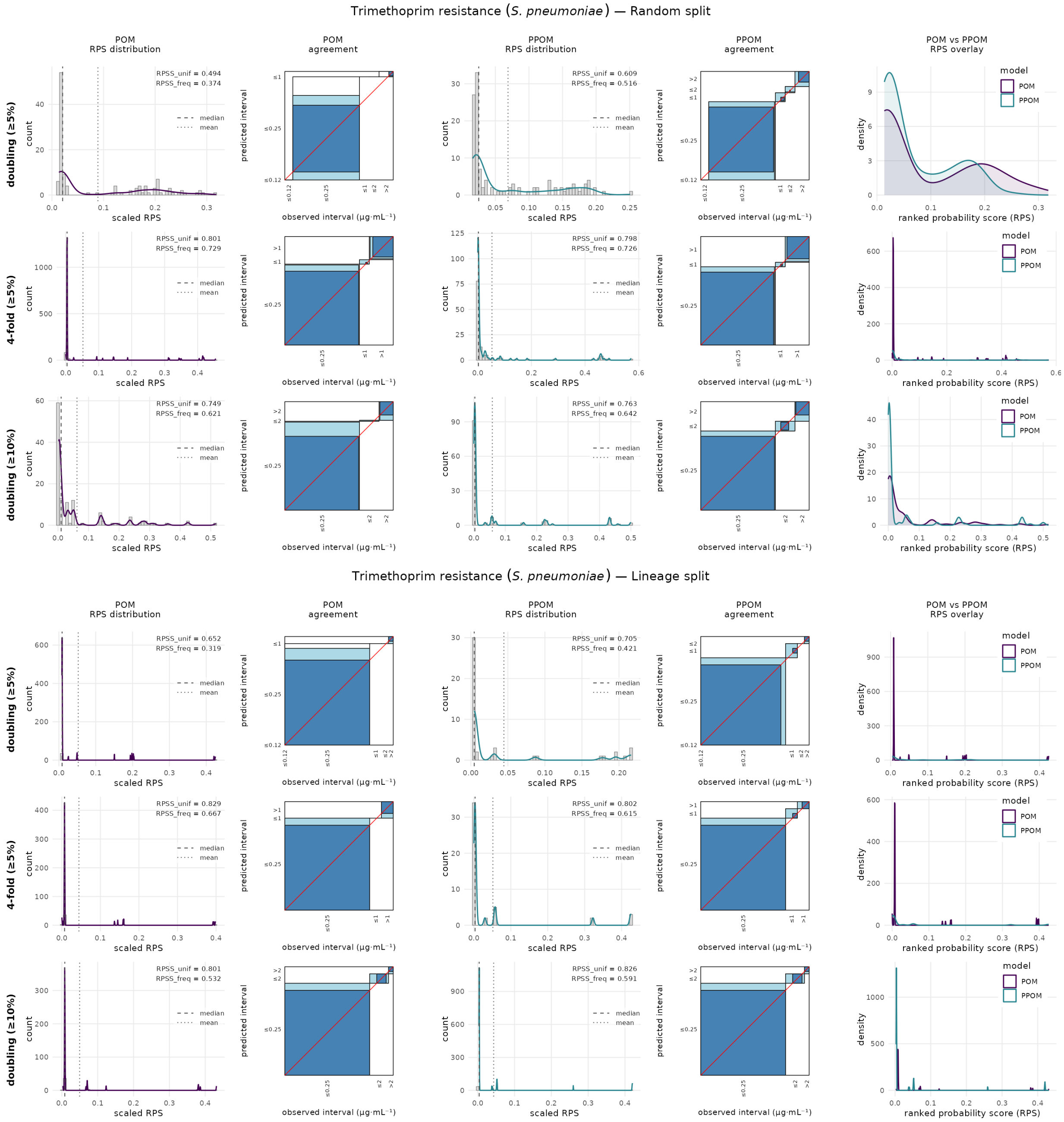


**Supplementary Figure S13.** Density plots of prediction errors as measured by RPS and RPSS, and agreement plots of predicted MIC versus observed phenotype, for POM and PPOM fit to *S. pneumoniae* with a trimethoprim resistance phenotype with different MIC discretization strategies. Top: Random train:test split. Bottom: one lineage subcluster withheld from training and used for prediction.

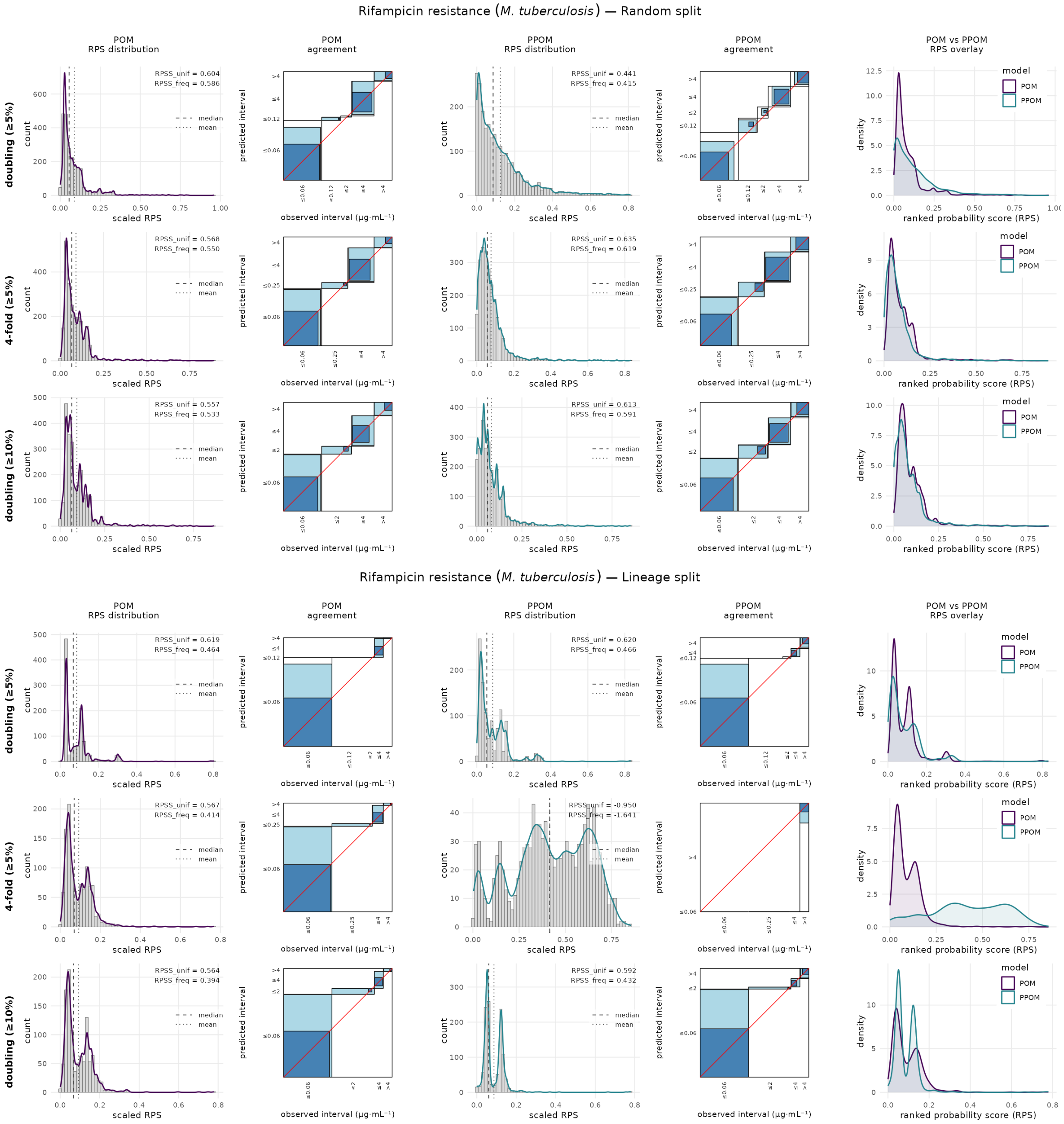


**Supplementary Figure S14.** Density plots of prediction errors as measured by RPS and RPSS, and agreement plots of predicted MIC versus observed phenotype, for POM and PPOM fit to *M. tuberculosis* with a rifampicin resistance phenotype with different MIC discretization strategies. Top: Random train:test split. Bottom: one lineage subcluster withheld from training and used for prediction.

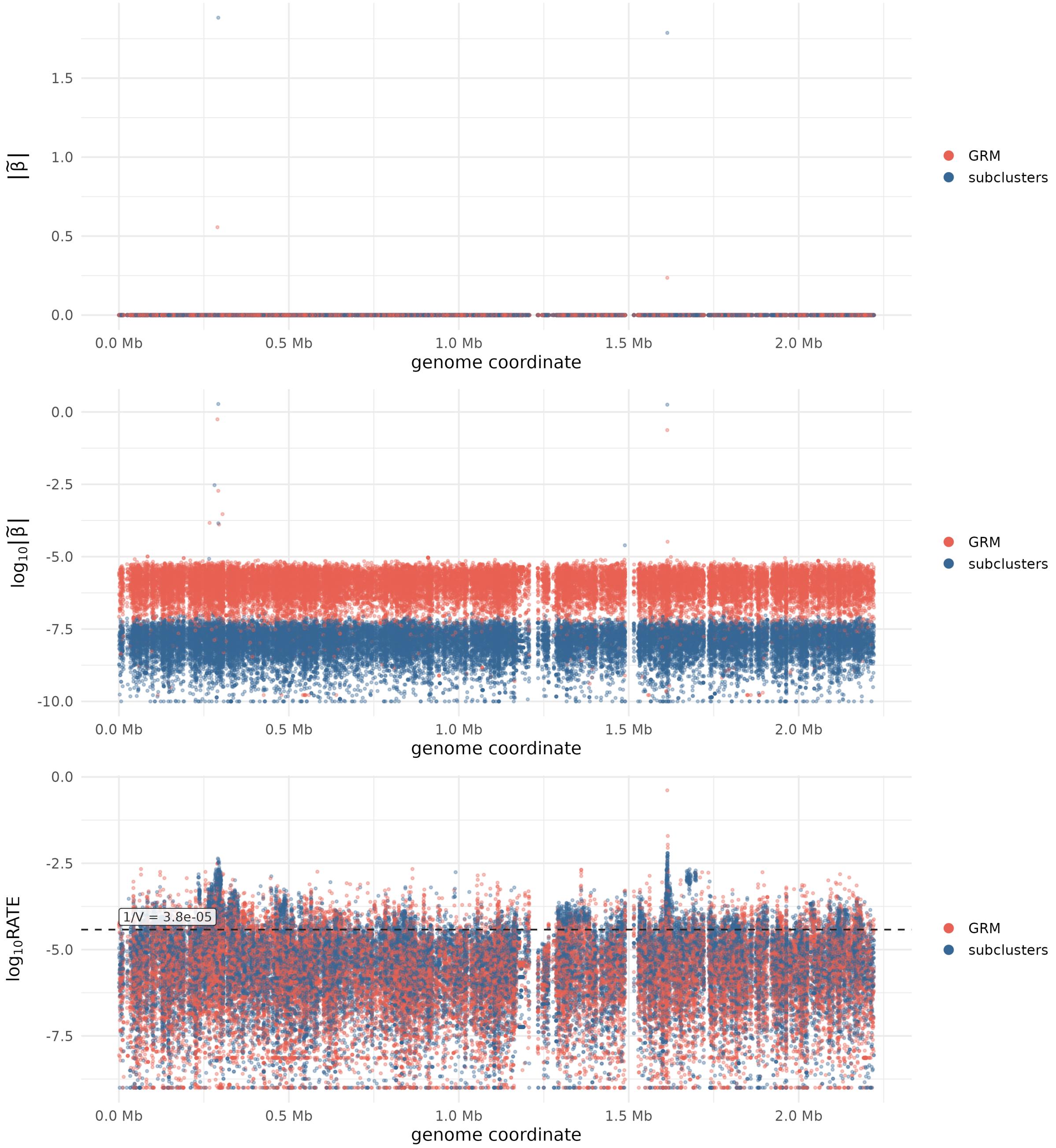


**Supplementary Figure S15.** Genome-wide variant effects and RATE values for 32,406 variants associated with penicillin resistance in $588$ *S. pneumoniae* isolates using the subclustering (blue) and genetic relatedness matrix (GRM, red) methods of population structure control. The top figure displays the posterior median absolute variant effect $|\tilde{\beta}|$, the middle their $\log_{10}$ scale, and the bottom their $\log_{10}$ RATE. The dashed line marks $1/V = 3.79 \times 10^{-5}$, the uniform share of the total RATE across the $V = 26{,}381$ LD-pruned variants. Both methods place their largest effects and highest RATE values at the same two regions, *pbp2x* ($\sim$0.29 Mb) and *pbp2b* ($\sim$1.61 Mb).

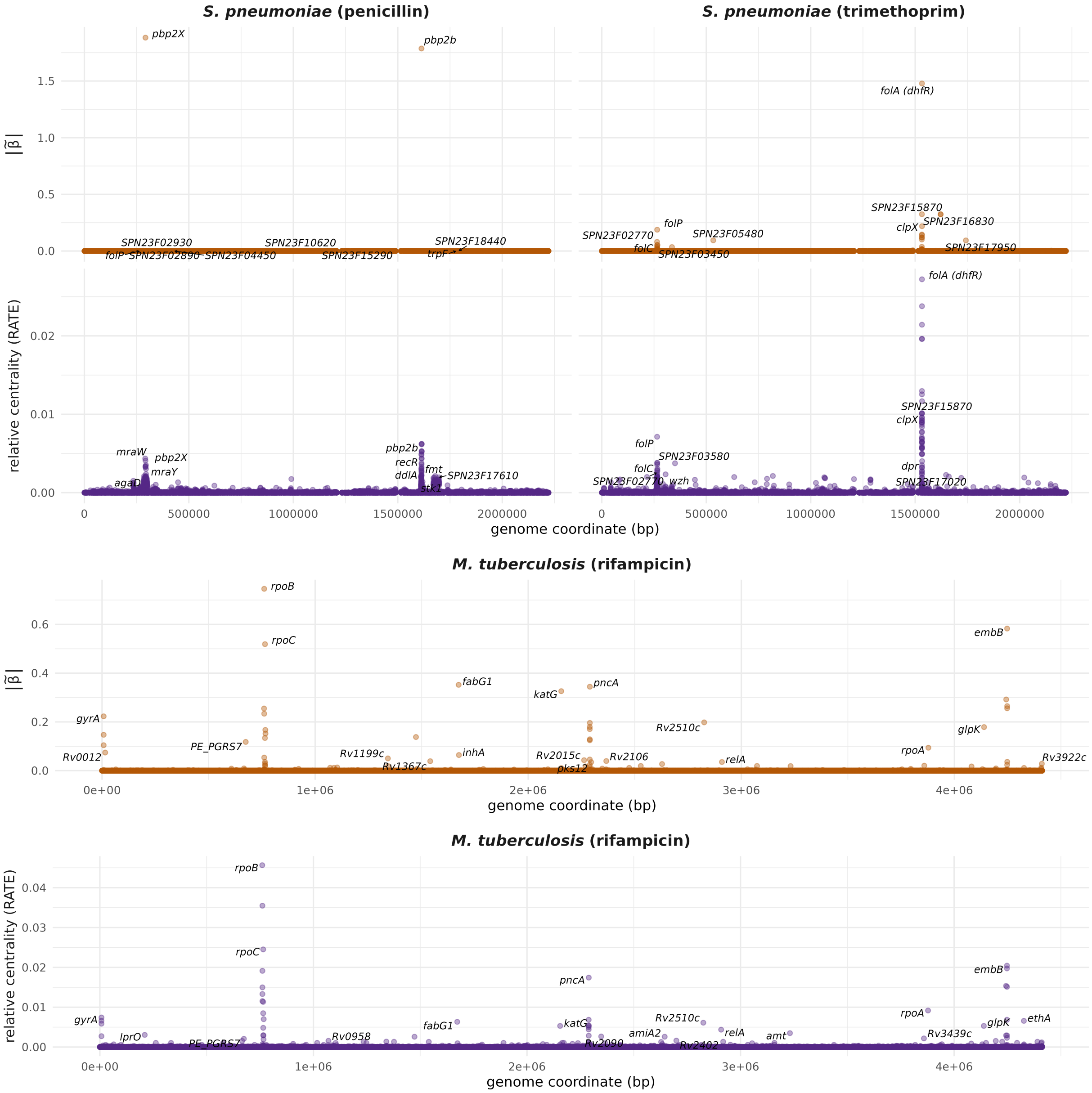


**Supplementary Figure S16.** Median coefficient magnitude, $|\tilde{\beta}|$, and the RATE values calculated from their values, using a logistic model with 32,406 variants associated with benzylpenicillin resistance in *S. pneumoniae*, 32,406 variants associated with trimethoprim resistance in *S. pneumoniae*, and 75,272 variants associated with rifampicin resistance in *M. tuberculosis*.

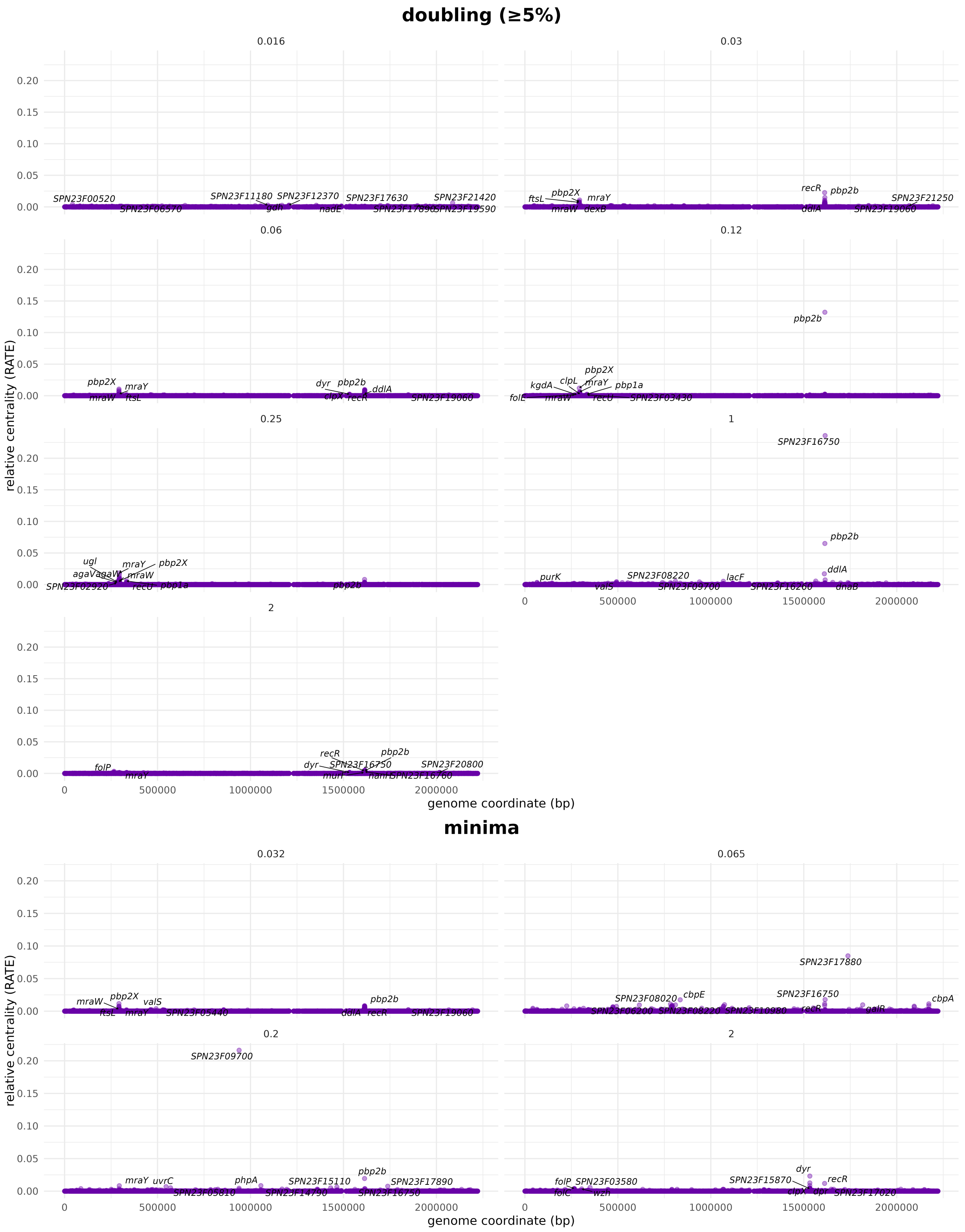


**Supplementary Figure S17.** RATE values calculated from variant coefficients from different breakpoints in a partial proportional odds model (PPOM), for 32,406 variants associated with benzylpenicillin resistance in *S. pneumoniae*. (A) Standard binning strategy (5% minimum frequency). (B) Minima binning strategy.

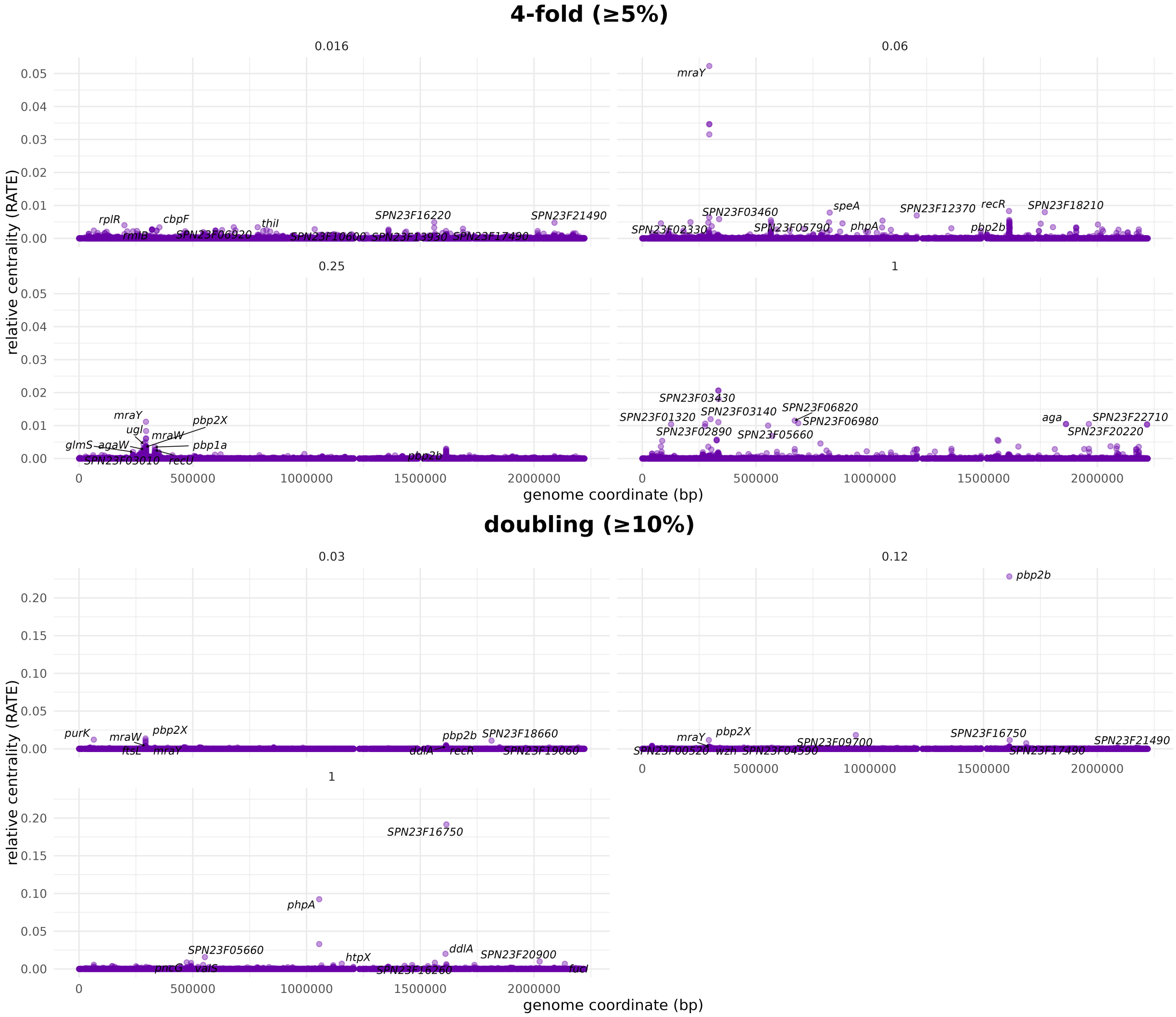


**Supplementary Figure S18.** RATE values calculated from variant coefficients from different breakpoints in a partial proportional odds model (PPOM), for 32,406 variants associated with benzylpenicillin resistance in *S. pneumoniae*. (C) Coarse binning strategy. (D) Standard binning strategy (10% minimum frequency).

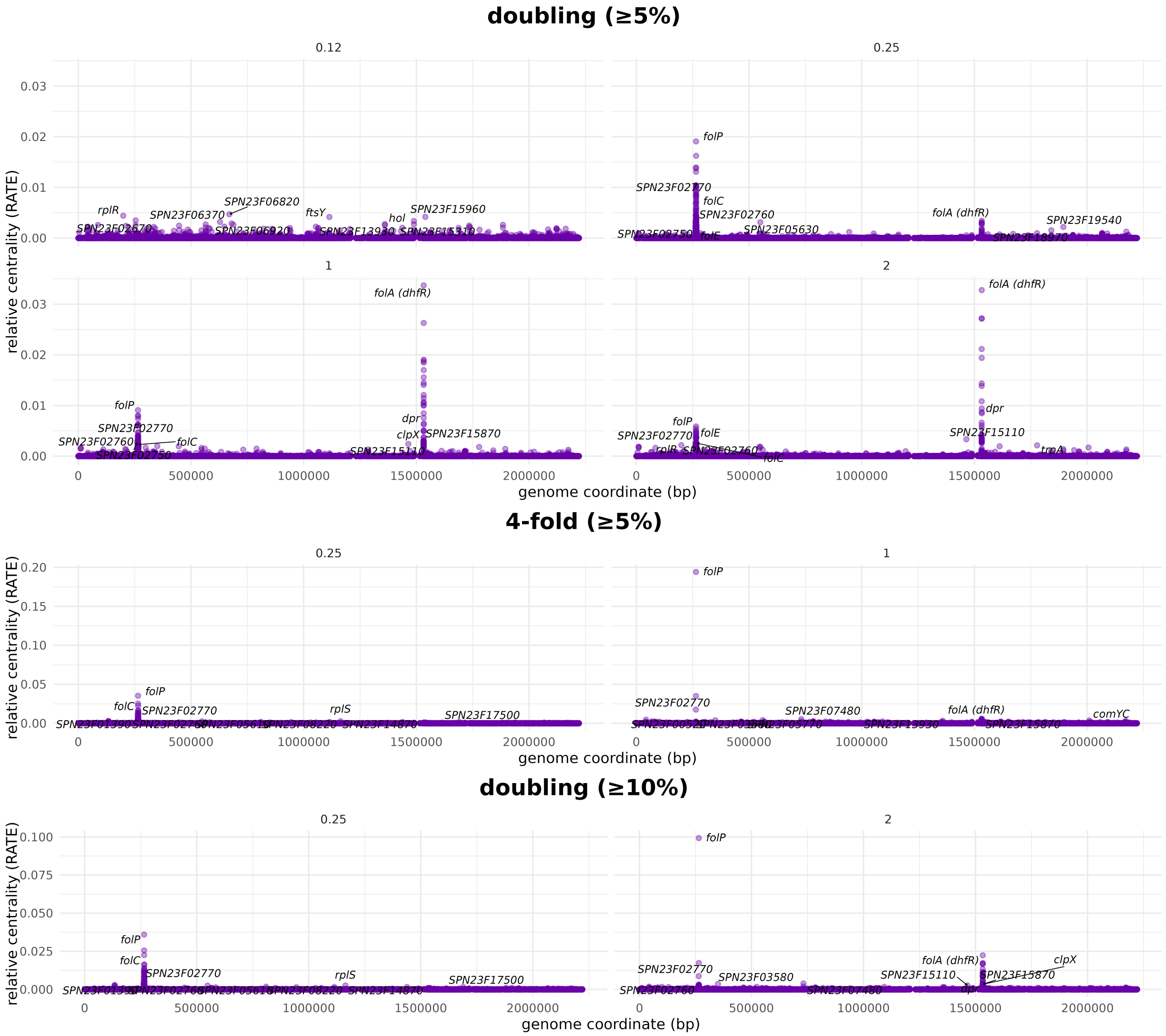


**Supplementary Figure S19.** RATE values calculated from variant coefficients from different breakpoints in a partial proportional odds model (PPOM), for 32,406 variants associated with trimethoprim resistance in *S. pneumoniae*. (A) Standard binning strategy (5% minimum frequency). (B) Coarse binning strategy. (C) Standard binning strategy (10% minimum frequency).

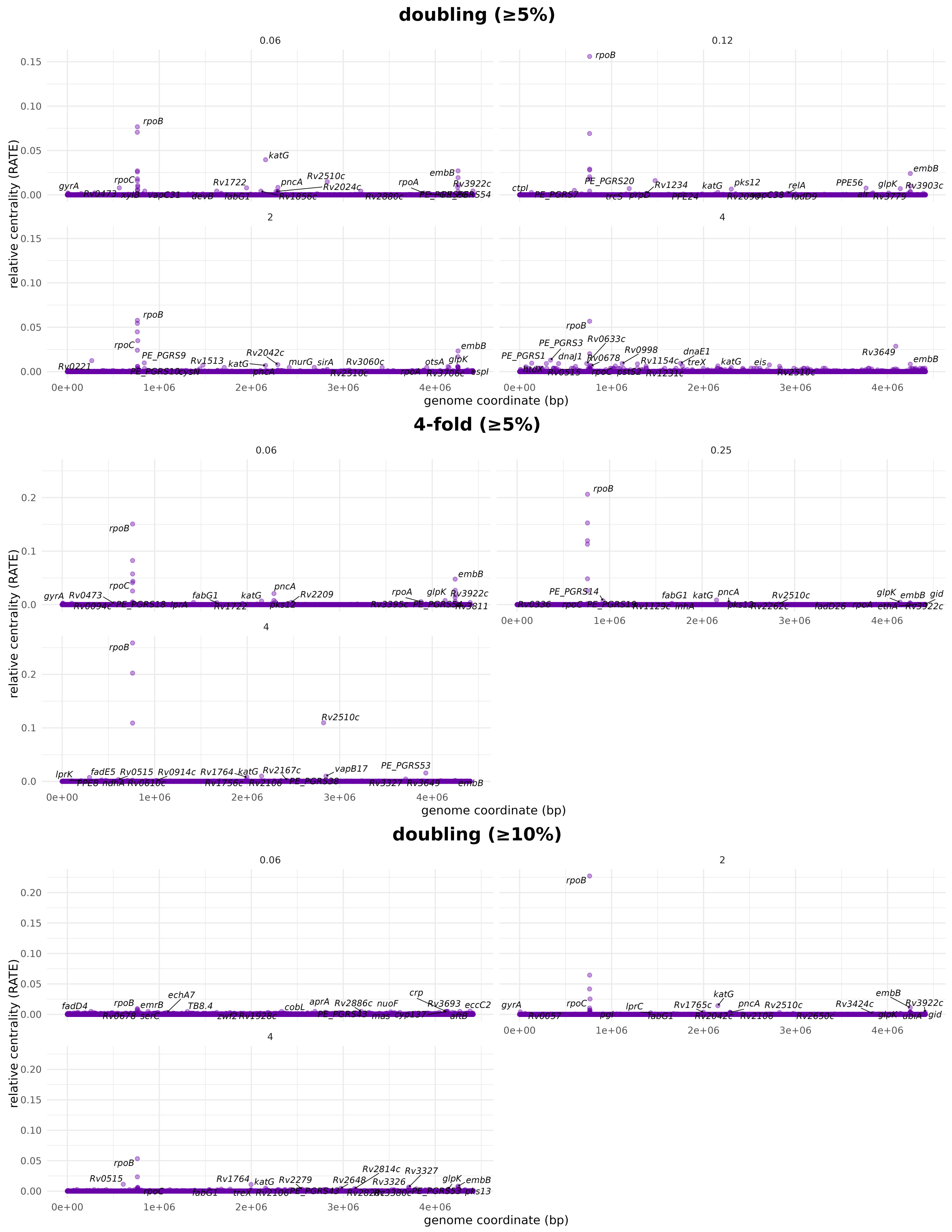


**Supplementary Figure S20.** RATE values calculated from variant coefficients from different breakpoints in a partial proportional odds model (PPOM), for 75,272 variants associated with rifampicin resistance in *M. tuberculosis*. (A) Standard binning strategy (5% minimum frequency). (B) Coarse binning strategy. (C) Standard binning strategy (10% minimum frequency).

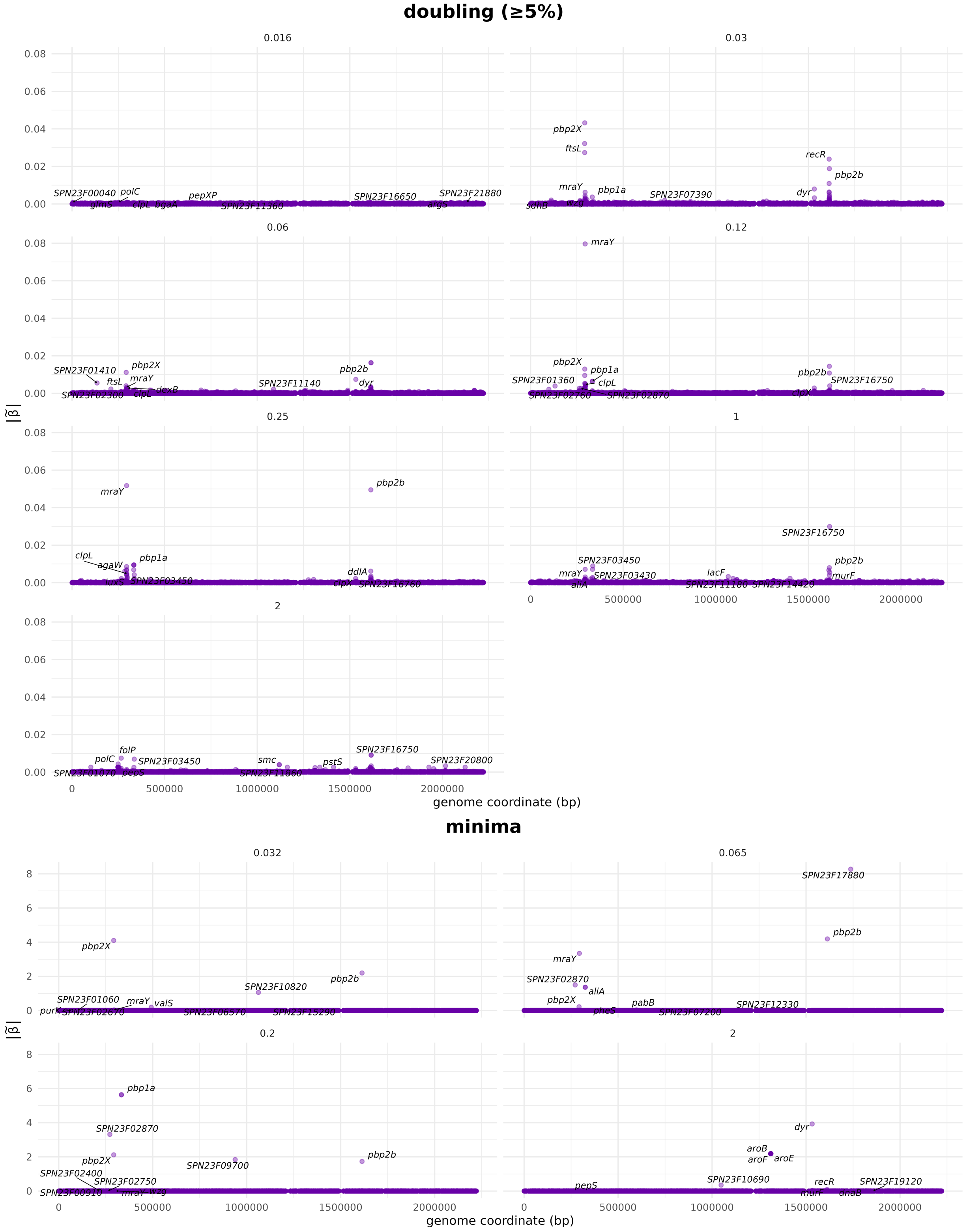


**Supplementary Figure S21.** Median coefficient magnitude, $|\tilde{\beta}|$, of variants at different breakpoints in a partial proportional odds model (PPOM), for 32,406 variants associated with benzylpenicillin resistance in *S. pneumoniae*. (A) Standard binning strategy (5% minimum frequency). (B) Minima binning strategy.

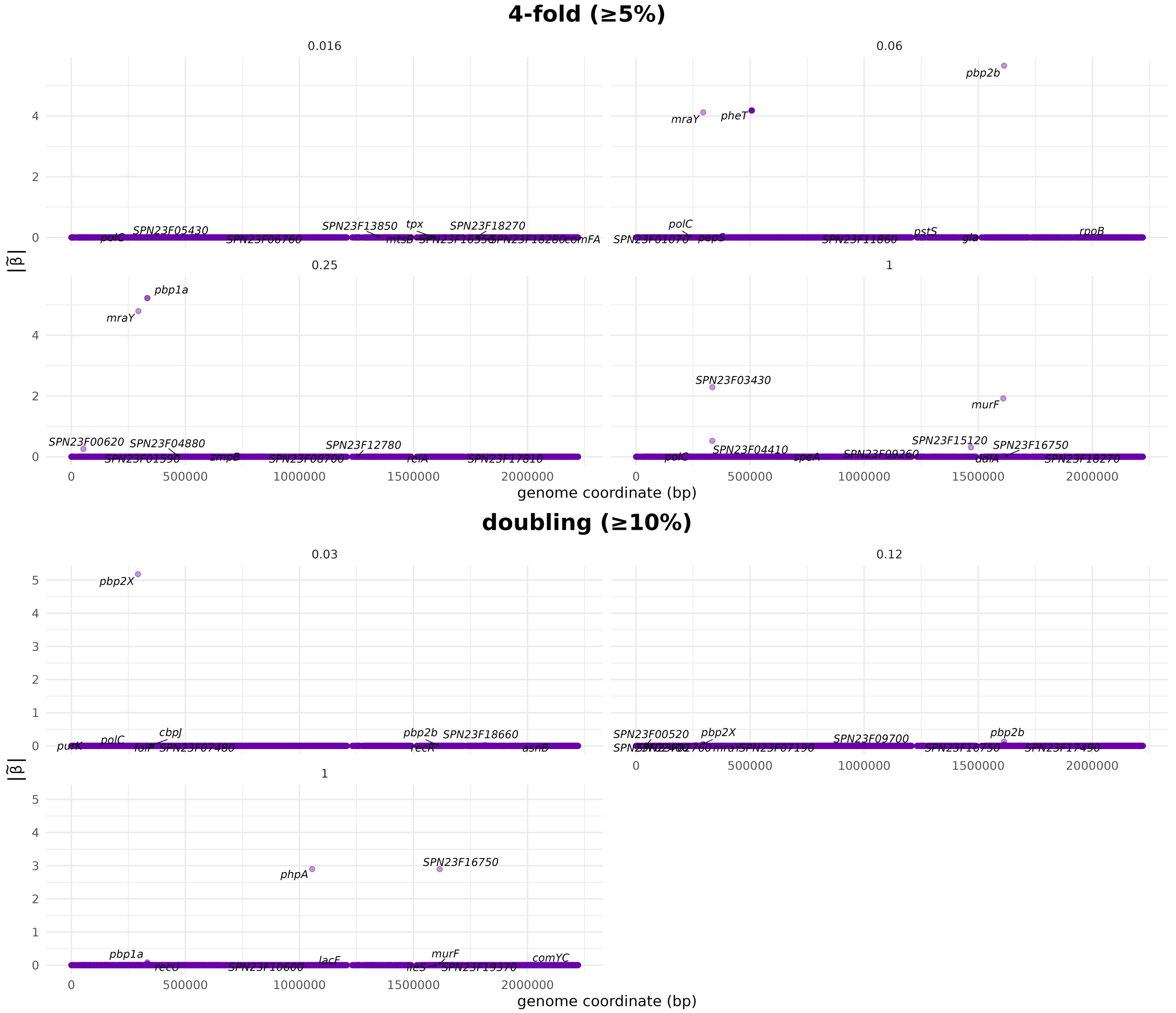


**Supplementary Figure S22.** Median coefficient magnitude, $|\tilde{\beta}|$, of variants at different breakpoints in a partial proportional odds model (PPOM), for 32,406 variants associated with benzylpenicillin resistance in *S. pneumoniae*. (C) Coarse binning strategy. (D) Standard binning strategy (10% minimum frequency).

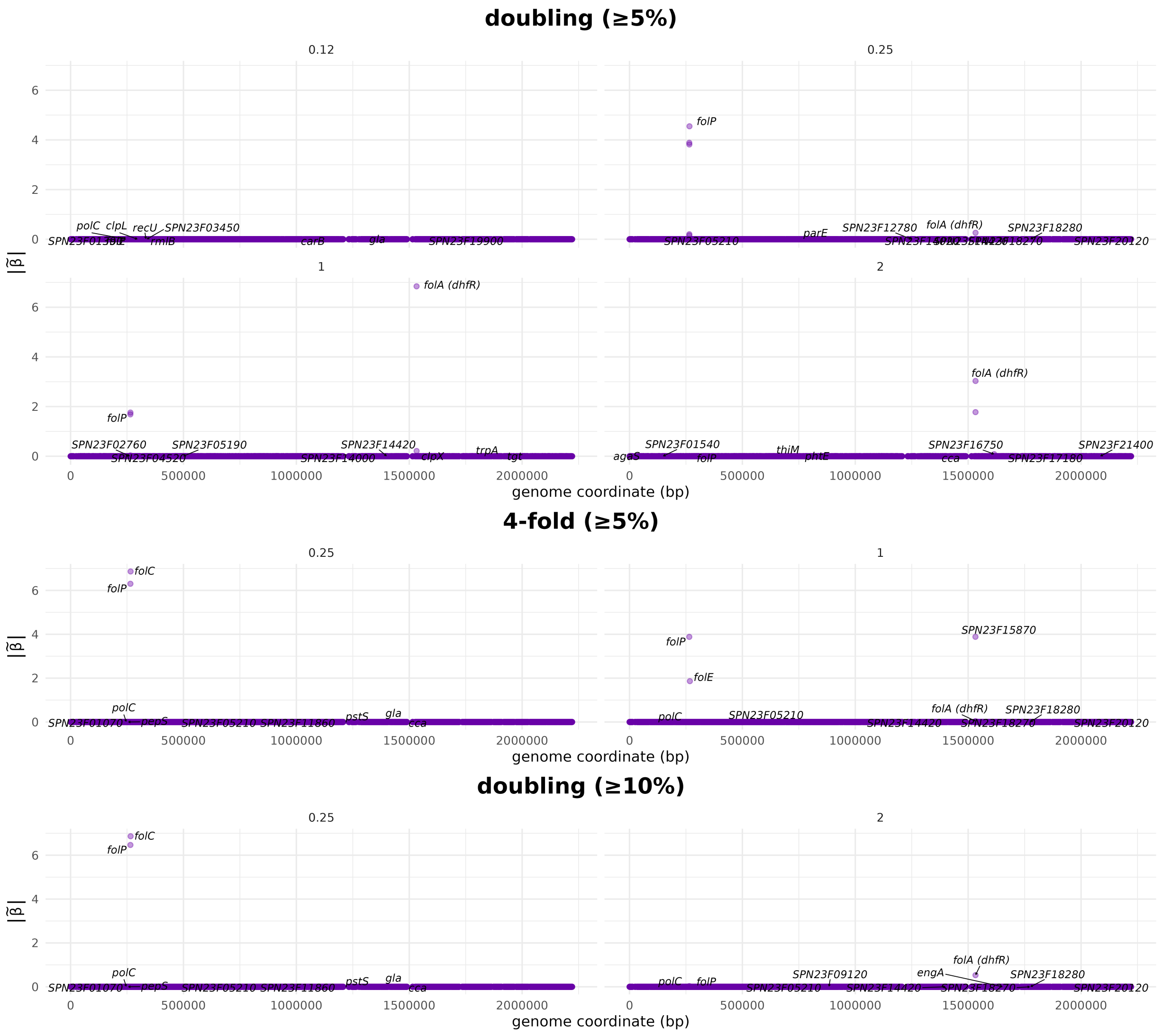

**Supplementary Figure S23.** Median coefficient magnitude, $|\tilde{\beta}|$, of variants at different breakpoints in a partial proportional odds model (PPOM), for 32,406 variants associated with trimethoprim resistance in *S. pneumoniae*. (A) Standard binning strategy (5% minimum frequency). (B) Coarse binning strategy. (C) Standard binning strategy (10% minimum frequency).

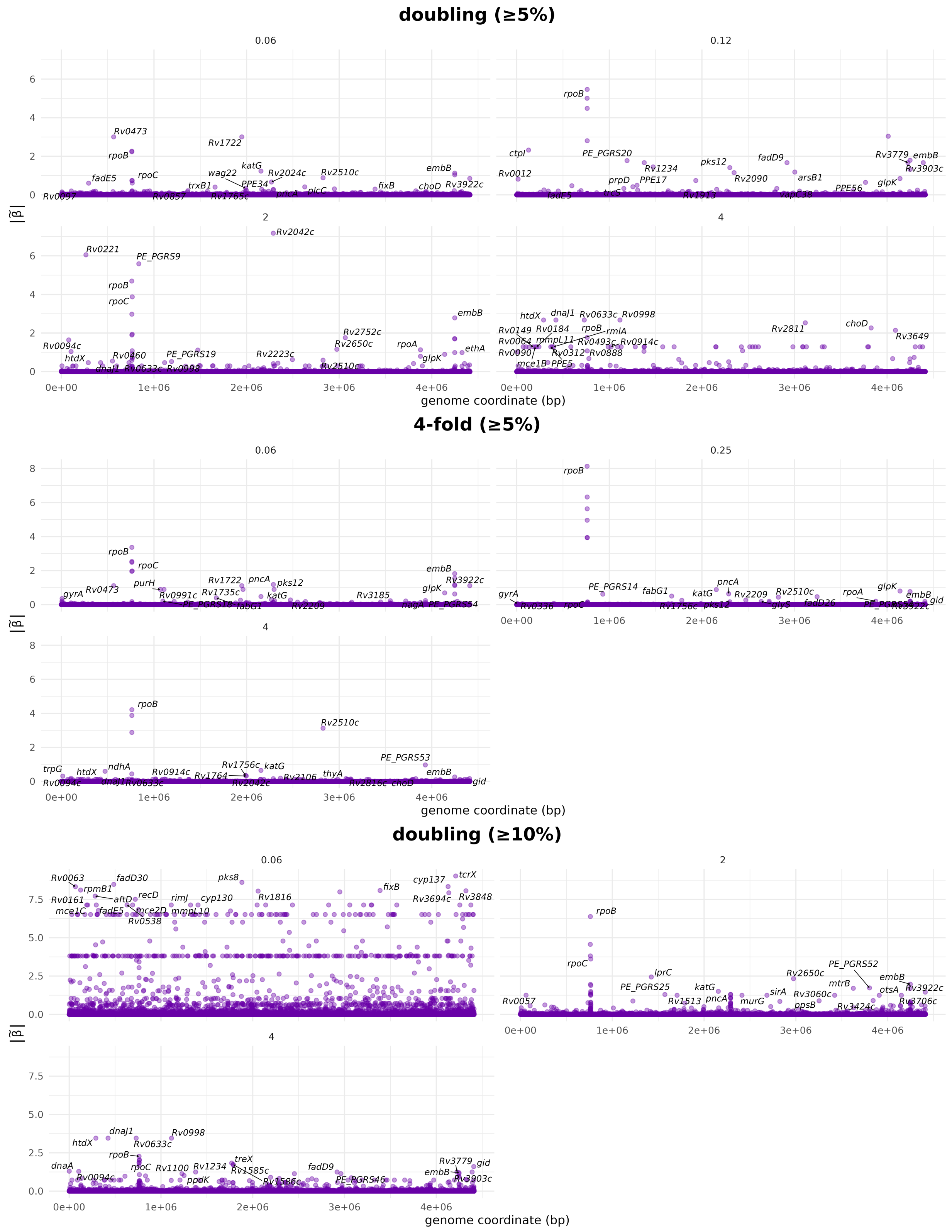


**Supplementary Figure S24.** Median coefficient magnitude, $|\tilde{\beta}|$, of variants at different breakpoints in a partial proportional odds model (PPOM), for 75,272 variants associated with rifampicin resistance in *M. tuberculosis*. (A) Standard binning strategy (5% minimum frequency). (B) Coarse binning strategy. (C) Standard binning strategy (10% minimum frequency).

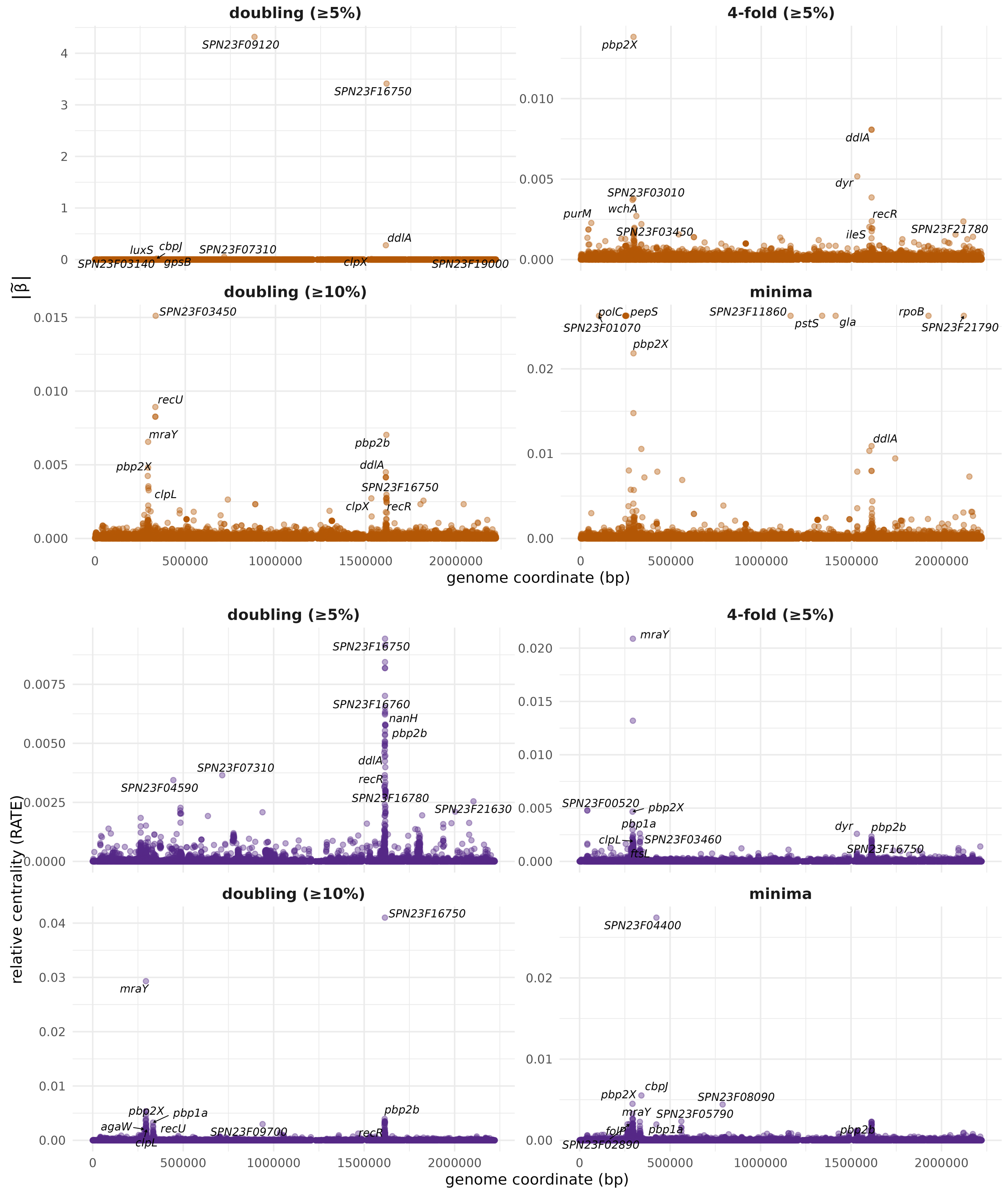


**Supplementary Figure S25.** Median coefficient magnitude, $|\tilde{\beta}|$, and the RATE values calculated from their values, using a proportional odds model (POM) with 32,406 variants associated with benzylpenicillin resistance in *S. pneumoniae*. (A) Standard binning strategy (5% minimum frequency). (B) Coarse binning strategy. (C) Standard binning strategy (10% minimum frequency). (D) Minima binning strategy.

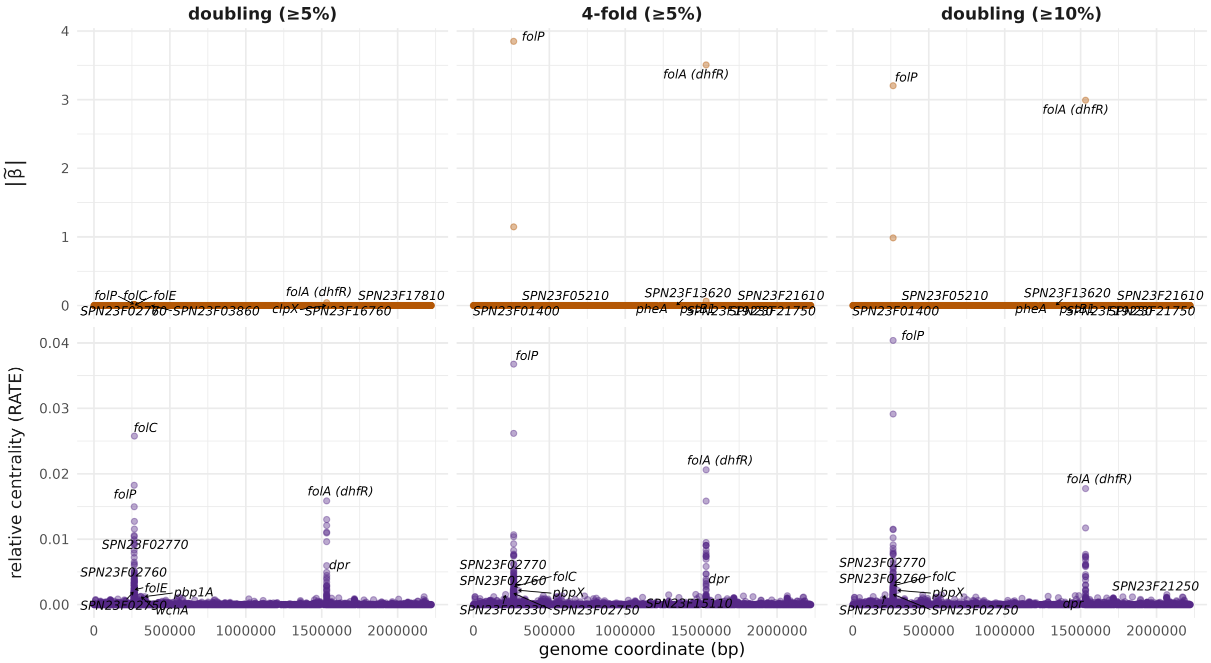


**Supplementary Figure S26.** Median coefficient magnitude, $|\tilde{\beta}|$, and the RATE values calculated from their values, using a proportional odds model (POM) with 32,406 variants associated with trimethoprim resistance in *S. pneumoniae*. (A) Standard binning strategy (5% minimum frequency). (B) Coarse binning strategy. (C) Standard binning strategy (10% minimum frequency).

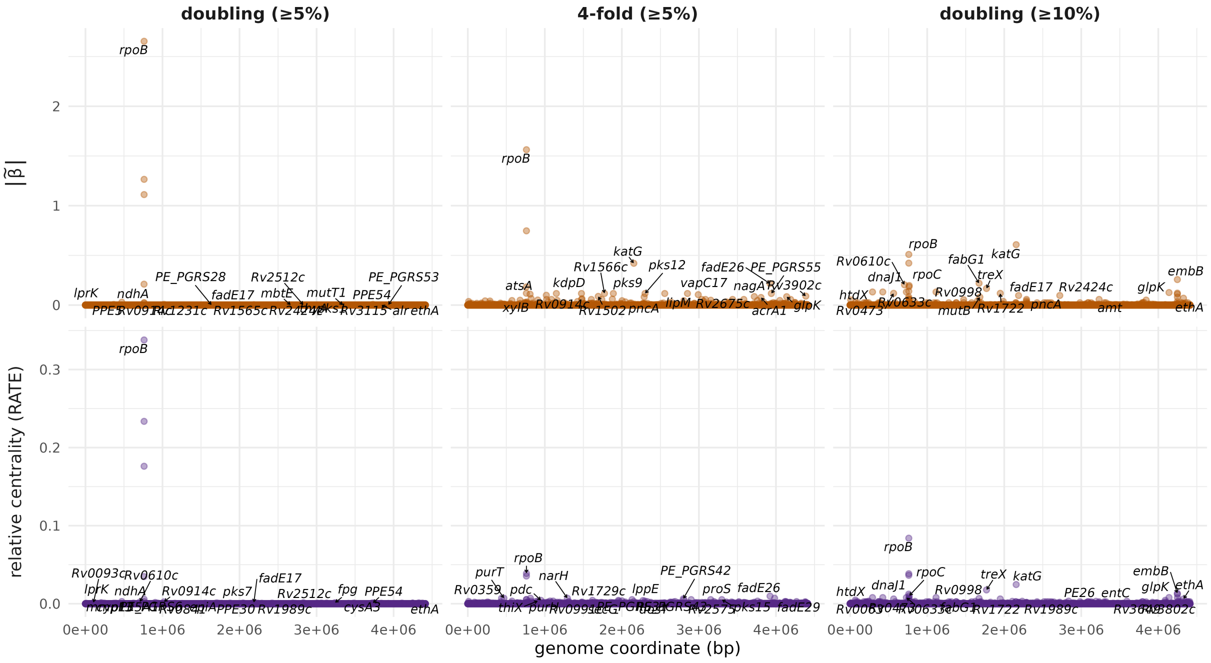


**Supplementary Figure S27.** Median coefficient magnitude, $|\tilde{\beta}|$, and the RATE values calculated from their values, using a proportional odds model (POM) with 75,272 variants associated with rifampicin resistance in *M. tuberculosis*. (A) Standard binning strategy (5% minimum frequency). (B) Coarse binning strategy.

**Supplementary Table ST8. Prediction evaluation metrics for ordered logistic models (POM and PPOM).** "Binning" describes the MIC discretization: *standard* uses the full doubling-dilution grid, *coarse* uses every-other doubling dilution, and *minima* binning was hand-binned at local minima in the phenotype; the minimum per-bin fraction enforced is shown in parentheses. Each MIC dataset was fit with both a proportional-odds model (POM) and a partial-proportional-odds model (PPOM). $K$ is the number of ordinal MIC categories after binning. $n_{train}$ is the number of isolates the model was fit on; $n_{test}$ is the number the metrics were computed on (for PPC, a random subset of the training dataset with a maximum of 500 isolates; for the random and single-lineage sets, a disjoint held-out subset). Balanced accuracy (bACC) is the mean over observed categories of the per-class balanced accuracy $(\text{sensitivity} + \text{specificity})/2$; PPV is the support-weighted mean of the per-class precision (positive predictive value). The mean and the median of the per-sample ranked probability score (RPS) across test isolates are reported, each scaled by $1/(K-1)$. Ranked probability skill score $RPSS_{unif}$ is calculated against predictions of uniform category frequencies, while $RPSS_{freq}$ is calculated against true category frequencies. Metrics with $^{\dagger}$ were calculated using only the categories observed at least once in the test dataset.

| Organism | AMR | Binning | K | Model | Evaluation | $n_{train}$ | $n_{test}$ | bACC | PPV | $RPS_{mean}$ | $RPS_{med}$ | $RPSS_{unif}$ | $RPSS_{freq}$ |
|---|---|---|---|---|---|---|---|---|---|---|---|---|---|
| *S. pneumoniae* | PEN | standard (≥5%) | 8 | PPOM | PPC | 611 | 123 | 0.689 | 0.657$^{\dagger}$ | 0.073 | 0.050 | 0.599 | 0.503 |
| | | | | | Random | 488 | 123 | 0.624 | 0.586$^{\dagger}$ | 0.062 | 0.020 | 0.663 | 0.505 |
| | | | | | Lineage | 565 | 46 | 0.620$^{\dagger}$ | 0.750$^{\dagger}$ | 0.176 | 0.152 | 0.086 | -2.889 |
| | | | | POM | PPC | 611 | 123 | 0.605 | 0.596$^{\dagger}$ | 0.079 | 0.044 | 0.570 | 0.467 |
| | | | | | Random | 488 | 123 | 0.590 | 0.549$^{\dagger}$ | 0.078 | 0.077 | 0.577 | 0.378 |
| | | | | | Lineage | 565 | 46 | 0.450$^{\dagger}$ | 0.000$^{\dagger}$ | 0.249 | 0.295 | -0.297 | -4.517 |
| | PEN | coarse (≥5%) | 5 | PPOM | PPC | 611 | 123 | 0.756 | 0.829$^{\dagger}$ | 0.046 | 0.004 | 0.725 | 0.662 |
| | | | | | Random | 488 | 123 | 0.708 | 0.806$^{\dagger}$ | 0.063 | 0.029 | 0.620 | 0.438 |
| | | | | | Lineage | 565 | 46 | 0.532$^{\dagger}$ | 0.603 | 0.189 | 0.129 | 0.293 | -1.984 |
| | | | | POM | PPC | 611 | 123 | 0.586 | 0.670$^{\dagger}$ | 0.079 | 0.029 | 0.528 | 0.420 |
| | | | | | Random | 488 | 123 | 0.579 | 0.696$^{\dagger}$ | 0.080 | 0.036 | 0.514 | 0.281 |
| | | | | | Lineage | 565 | 46 | 0.474$^{\dagger}$ | 0.737$^{\dagger}$ | 0.165 | 0.076 | 0.381 | -1.612 |
| | PEN | standard (≥10%) | 4 | PPOM | PPC | 611 | 123 | 0.801 | 0.771 | 0.066 | 0.025 | 0.718 | 0.649 |
| | | | | | Random | 488 | 123 | 0.767 | 0.762 | 0.081 | 0.055 | 0.653 | 0.498 |
| | | | | | Lineage | 565 | 46 | 0.500$^{\dagger}$ | 0.783$^{\dagger}$ | 0.102 | 0.006 | 0.599 | -0.450 |
| | | | | POM | PPC | 611 | 123 | 0.633 | 0.636$^{\dagger}$ | 0.102 | 0.079 | 0.565 | 0.458 |
| | | | | | Random | 488 | 123 | 0.664 | 0.475 | 0.100 | 0.038 | 0.572 | 0.381 |
| | | | | | Lineage | 565 | 46 | 0.549$^{\dagger}$ | 0.717$^{\dagger}$ | 0.218 | 0.167 | 0.148 | -2.083 |
| | PEN | minima (≥5%) | 5 | PPOM | PPC | 611 | 123 | 0.790 | 0.817 | 0.043 | 0.005 | 0.815 | 0.754 |
| | | | | | Random | 488 | 123 | 0.776 | 0.775$^{\dagger}$ | 0.062 | 0.032 | 0.741 | 0.579 |
| | | | | | Lineage | 565 | 46 | 0.731$^{\dagger}$ | 0.959$^{\dagger}$ | 0.116 | 0.031 | 0.246 | -3.424 |

*continued on next page*

*Table ST8 continued from previous page*

| Organism | AMR | Binning | K | Model | Evaluation | $n_{train}$ | $n_{test}$ | bACC | PPV | $RPS_{mean}$ | $RPS_{med}$ | $RPSS_{unif}$ | $RPSS_{freq}$ |
|---|---|---|---|---|---|---|---|---|---|---|---|---|---|
| | | | | POM | PPC | 611 | 123 | 0.577 | 0.682$^\dagger$ | 0.086 | 0.045 | 0.632 | 0.511 |
| | | | | | Random | 488 | 123 | 0.669 | 0.714$^\dagger$ | 0.083 | 0.034 | 0.653 | 0.435 |
| | | | | | Lineage | 565 | 46 | 0.660$^\dagger$ | 0.940$^\dagger$ | 0.169 | 0.060 | -0.105 | -5.485 |
| *S. pneumoniae* | TMP | standard (≥5%) | 5 | PPOM | PPC | 606 | 122 | 0.713 | 0.762$^\dagger$ | 0.060 | 0.005 | 0.660 | 0.547 |
| | | | | | Random | 484 | 122 | 0.670 | 0.703$^\dagger$ | 0.069 | 0.025 | 0.609 | 0.516 |
| | | | | | Lineage | 560 | 46 | 0.741$^\dagger$ | 0.839$^\dagger$ | 0.045 | 0.005 | 0.705 | 0.421 |
| | | | | POM | PPC | 606 | 122 | 0.552 | 0.669$^\dagger$ | 0.086 | 0.020 | 0.509 | 0.347 |
| | | | | | Random | 484 | 122 | 0.543 | 0.679$^\dagger$ | 0.089 | 0.021 | 0.494 | 0.374 |
| | | | | | Lineage | 560 | 46 | 0.660$^\dagger$ | 0.828$^\dagger$ | 0.053 | 0.009 | 0.652 | 0.319 |
| | TMP | coarse (≥5%) | 3 | PPOM | PPC | 606 | 122 | 0.831 | 0.894 | 0.055 | 0.002 | 0.788 | 0.679 |
| | | | | | Random | 484 | 122 | 0.809 | 0.868 | 0.053 | 0.003 | 0.798 | 0.726 |
| | | | | | Lineage | 560 | 46 | 0.775 | 0.885 | 0.051 | 0.003 | 0.802 | 0.615 |
| | | | | POM | PPC | 606 | 122 | 0.750 | 0.858$^\dagger$ | 0.074 | 0.003 | 0.715 | 0.568 |
| | | | | | Random | 484 | 122 | 0.792 | 0.860 | 0.052 | 0.004 | 0.801 | 0.729 |
| | | | | | Lineage | 560 | 46 | 0.775 | 0.898$^\dagger$ | 0.044 | 0.007 | 0.829 | 0.667 |
| | TMP | standard (≥10%) | 3 | PPOM | PPC | 606 | 122 | 0.827 | 0.878 | 0.057 | 0.004 | 0.771 | 0.615 |
| | | | | | Random | 484 | 122 | 0.848 | 0.879 | 0.059 | 0.003 | 0.763 | 0.642 |
| | | | | | Lineage | 560 | 46 | 0.863 | 0.925 | 0.043 | 0.004 | 0.826 | 0.591 |
| | | | | POM | PPC | 606 | 122 | 0.794 | 0.840 | 0.072 | 0.004 | 0.711 | 0.514 |
| | | | | | Random | 484 | 122 | 0.735 | 0.665 | 0.062 | 0.010 | 0.749 | 0.621 |
| | | | | | Lineage | 560 | 46 | 0.863 | 0.925 | 0.050 | 0.008 | 0.801 | 0.532 |
| *M. tuberculosis* | RIF | standard (≥5%) | 5 | PPOM | PPC | 11622 | 500 | 0.680 | 0.537 | 0.100 | 0.075 | 0.550 | 0.528 |
| | | | | | Random | 9297 | 2325 | 0.695 | 0.568 | 0.124 | 0.086 | 0.441 | 0.415 |
| | | | | | Lineage | 10519 | 1103 | 0.668 | 0.579$^\dagger$ | 0.086 | 0.054 | 0.620 | 0.466 |
| | | | | POM | PPC | 11622 | 500 | 0.663 | 0.572 | 0.095 | 0.057 | 0.572 | 0.551 |
| | | | | | Random | 9297 | 2325 | 0.678 | 0.586 | 0.088 | 0.055 | 0.604 | 0.586 |
| | | | | | Lineage | 10519 | 1103 | 0.627 | 0.564$^\dagger$ | 0.086 | 0.068 | 0.619 | 0.464 |
| | RIF | coarse (≥5%) | 4 | PPOM | PPC | 11622 | 500 | 0.819 | 0.736 | 0.078 | 0.063 | 0.631 | 0.614 |

*continued on next page*

*Table ST8 continued from previous page*

| Organism | AMR | Binning | K | Model | Evaluation | $n_{train}$ | $n_{test}$ | bACC | PPV | $RPS_{mean}$ | $RPS_{med}$ | $RPSS_{unif}$ | $RPSS_{freq}$ |
|---|---|---|---|---|---|---|---|---|---|---|---|---|---|
| | | | | | Random | 9297 | 2325 | 0.788 | 0.691 | 0.077 | 0.057 | 0.635 | 0.619 |
| | | | | | Lineage | 10519 | 1103 | 0.501 | 0.917† | 0.415 | 0.416 | -0.950 | -1.641 |
| | | | | POM | PPC | 11622 | 500 | 0.713 | 0.542 | 0.093 | 0.076 | 0.562 | 0.542 |
| | | | | | Random | 9297 | 2325 | 0.703 | 0.584 | 0.091 | 0.067 | 0.568 | 0.550 |
| | | | | | Lineage | 10519 | 1103 | 0.670 | 0.530 | 0.092 | 0.070 | 0.567 | 0.414 |
| | RIF | standard (≥10%) | 4 | PPOM | PPC | 11622 | 500 | 0.864 | 0.829 | 0.064 | 0.047 | 0.699 | 0.679 |
| | | | | | Random | 9297 | 2325 | 0.767 | 0.655 | 0.082 | 0.060 | 0.613 | 0.591 |
| | | | | | Lineage | 10519 | 1103 | 0.705 | 0.648 | 0.087 | 0.060 | 0.592 | 0.432 |
| | | | | POM | PPC | 11622 | 500 | 0.705 | 0.526 | 0.102 | 0.079 | 0.517 | 0.485 |
| | | | | | Random | 9297 | 2325 | 0.708 | 0.587 | 0.094 | 0.065 | 0.557 | 0.533 |
| | | | | | Lineage | 10519 | 1103 | 0.683 | 0.572 | 0.093 | 0.067 | 0.564 | 0.394 |

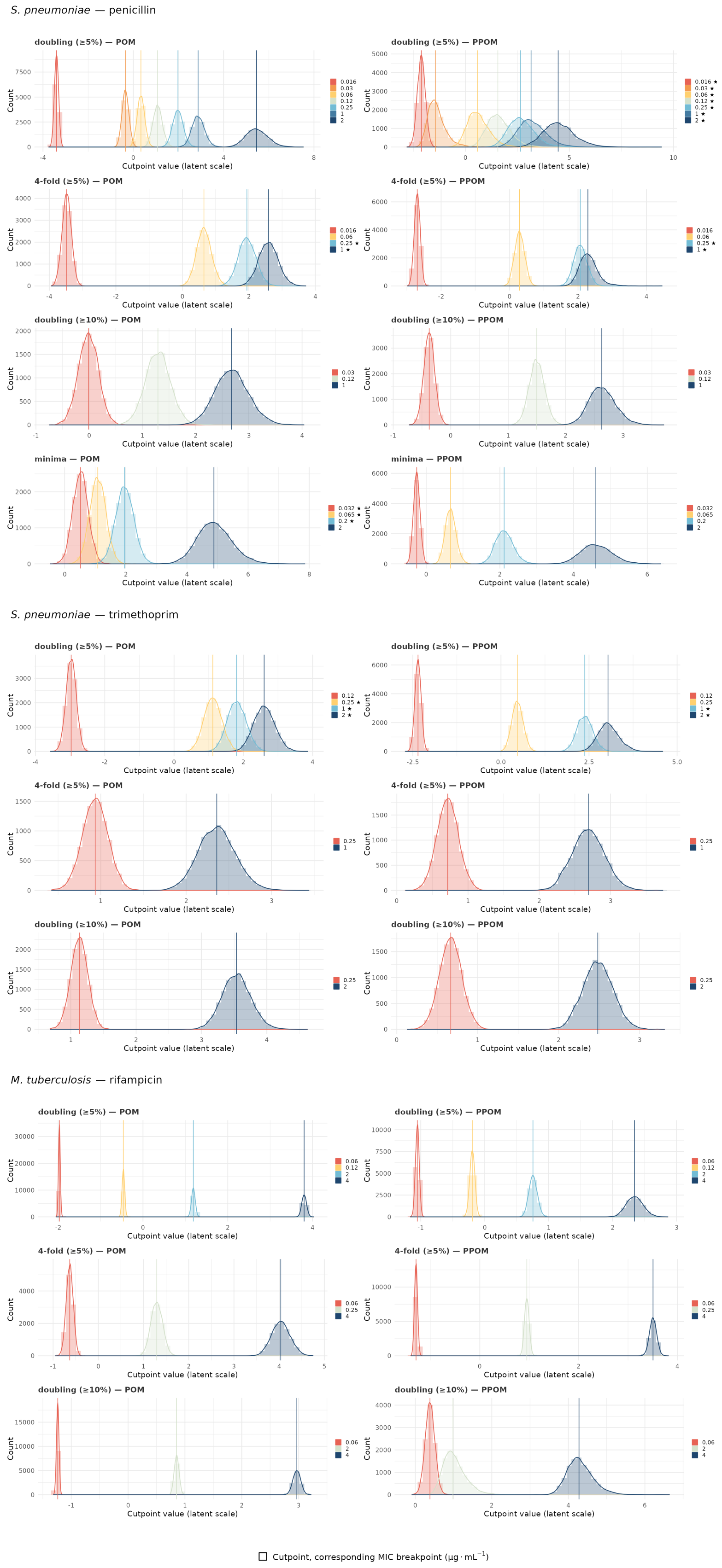


**Supplementary Figure S28.** Posterior cutpoint locations for various phenotype discretization strategies using a proportional odds model (POM) and a partial proportional odds model (PPOM). (A) Penicillin resistance in *S. pneumoniae*. (B) Trimethoprim resistance in *S. pneumoniae*. (C) Rifampicin resistance in *M. tuberculosis*.

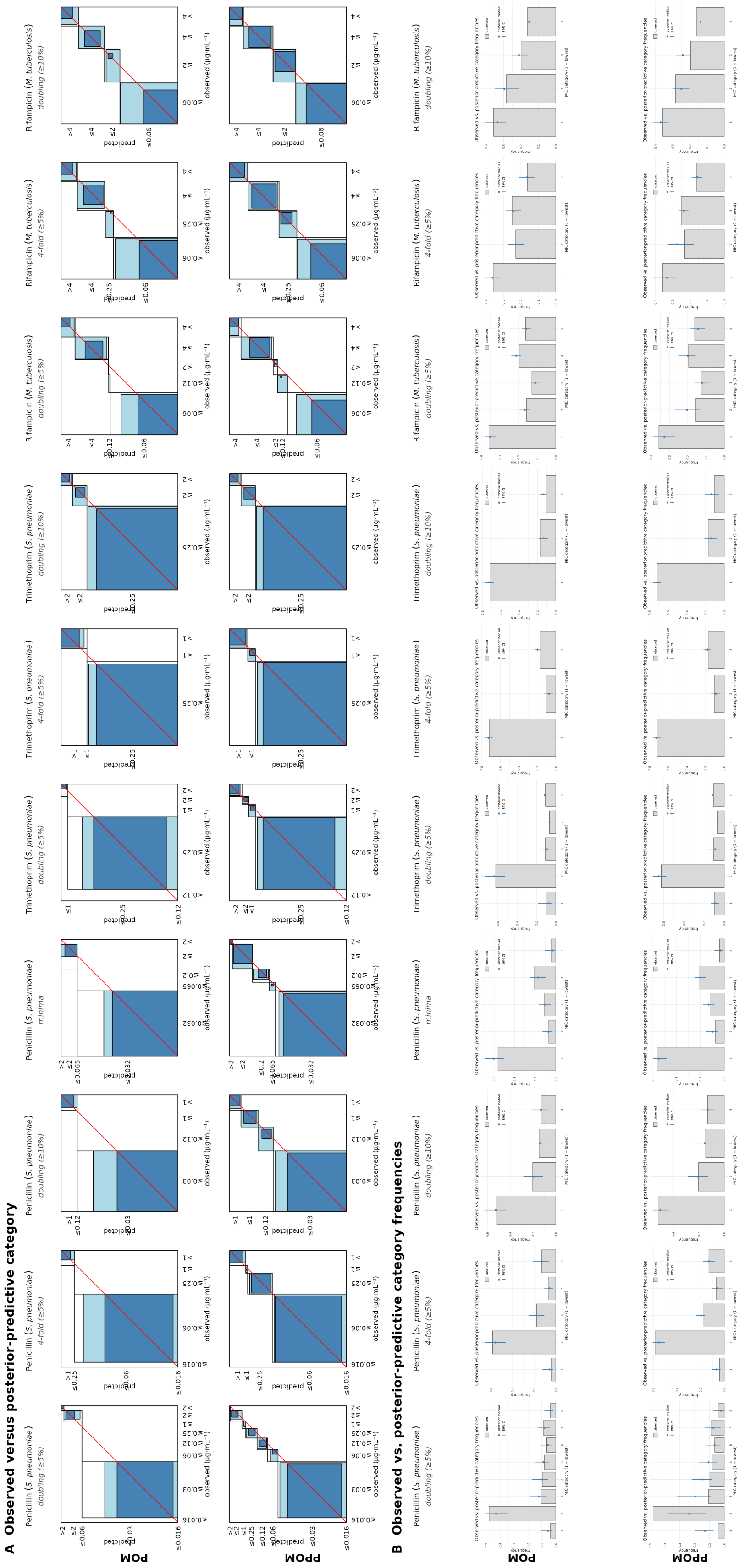


**Supplementary Figure S29.** (A) Observed versus posterior-predictive category for isolates used to fit the model. (B) Observed vs posterior-predictive category frequencies across all isolates, 89% credible intervals.

| Gene | Binning | Model | Position | RATE | $\lvert\beta\rvert$ | Residue | Variants | $n_{nt}$ | $n_{aa}$ | Effects |
|---|---|---|---|---|---|---|---|---|---|---|
| *rpoA* | – | logistic | 3877960 | 0.00917 | 0.094 | Val183 | Val183Gly, Val183Ala | 2 | 2 | missense |
| *rpoA* | doubling (≥5%) | POM | 3877949 | 4.67e−5 | 1.47e−7 | Thr187 | Thr187Ser, Thr187Ala, Thr187Pro | 3 | 3 | missense |
| | | PPOM | 3877960 | 0.00209 | 0.0624 | Val183 | Val183Gly, Val183Ala | 2 | 2 | missense |
| *rpoA* | doubling (≥10%) | POM | 3877960 | 2.00e−4 | 0.0122 | Val183 | Val183Gly, Val183Ala | 2 | 2 | missense |
| | | PPOM | 3877960 | 9.28e−5 | 0.304 | Val183 | Val183Gly, Val183Ala | 2 | 2 | missense |
| *rpoA* | 4-fold (≥5%) | POM | 3877939 | 4.13e−5 | 8.64e−6 | Asp190 | Asp190Gly | 1 | 1 | missense |
| | | PPOM | 3877960 | 0.00546 | 0.131 | Val183 | Val183Gly, Val183Ala | 2 | 2 | missense |
| *rpoB* | – | logistic | 760312 | 0.0456 | 0.255 | Val170 | high polymorphism | 10 | 5 | missense, synonymous |
| *rpoB* | doubling (≥5%) | POM | 761155 | 0.338 | 2.65 | Ser450 | Ser450*, Ser450Trp, Ser450Leu | 3 | 3 | missense, stop gained |
| | | PPOM | 761155 | 0.156 | 4.48 | Ser450 | Ser450*, Ser450Trp, Ser450Leu | 3 | 3 | missense, stop gained |
| *rpoB* | doubling (≥10%) | POM | 761139 | 0.084 | 0.509 | His445 | high polymorphism | 203 | 63 | frameshift, missense, stop gained |
| | | PPOM | 761155 | 0.227 | 6.38 | Ser450 | Ser450*, Ser450Trp, Ser450Leu | 3 | 3 | missense, stop gained |
| *rpoB* | 4-fold (≥5%) | POM | 761155 | 0.0388 | 1.56 | Ser450 | Ser450*, Ser450Trp, Ser450Leu | 3 | 3 | missense, stop gained |
| | | PPOM | 761155 | 0.259 | 4.21 | Ser450 | Ser450*, Ser450Trp, Ser450Leu | 3 | 3 | missense, stop gained |
| *rpoC* | – | logistic | 764817 | 0.0245 | 0.519 | Val483 | Val483Ala, Val483Gly | 2 | 2 | missense |
| *rpoC* | doubling (≥5%) | POM | 764817 | 1.67e−4 | 3.40e−7 | Val483 | Val483Ala, Val483Gly | 2 | 2 | missense |
| | | PPOM | 765462 | 0.0349 | 3.87 | Asn698 | Asn698Ser | 1 | 1 | missense |
| *rpoC* | doubling (≥10%) | POM | 766487 | 0.0101 | 0.197 | Pro1040 | Pro1040Thr, Pro1040Ala, Pro1040Ser | 3 | 3 | missense |
| | | PPOM | 765462 | 0.0253 | 3.61 | Asn698 | Asn698Ser | 1 | 1 | missense |
| *rpoC* | 4-fold (≥5%) | POM | 766772 | 4.51e−4 | 0.0227 | Val1135 | high polymorphism | 157 | 25 | conservative inframe insertion, disruptive inframe insertion, frameshift, missense, synonymous |
| | | PPOM | 764817 | 0.0437 | 1.98 | Val483 | Val483Ala, Val483Gly | 2 | 2 | missense |

**Supplementary Table ST5. Most significant variant in each *rpo* gene, per model and binning.** The variant with the largest RATE across cutpoints was chosen, and the $|\beta|$ at that cutpoint recorded. $n_{nt}$ and $n_{aa}$ are the numbers of unique nucleotide and amino acid alleles at the residue; positions with more than three protein alleles are marked *high polymorphism*.

| Organism | AMR | Evaluation | $n_{train}$ | $n_{test}$ | Sens | Spec | bACC | AUC | Brier | $F_1$ | VME | ME |
|---|---|---|---|---|---|---|---|---|---|---|---|---|
| *S. pneumoniae* | PEN | PPC | 588 | 118 | 0.796 | 0.957 | 0.876 | 0.921 | 0.098 | 0.857 | 0.204 | 0.043 |
| | | Random | 470 | 118 | 0.800 | 0.952 | 0.876 | 0.901 | 0.110 | 0.863 | 0.200 | 0.048 |
| | | Lineage | 542 | 46 | 0.761 | – | – | – | 0.104 | 0.864 | 0.239 | – |
| *S. pneumoniae* | TMP | PPC | 600 | 120 | 0.895 | 0.980 | 0.937 | 0.991 | 0.027 | 0.895 | 0.105 | 0.020 |
| | | Random | 480 | 120 | 0.818 | 0.959 | 0.889 | 0.991 | 0.037 | 0.818 | 0.182 | 0.041 |
| | | Lineage | 555 | 45 | 0.750 | 1.000 | 0.875 | 0.915 | 0.023 | 0.857 | 0.250 | 0.000 |
| *M. tuberculosis* | RIF | PPC | 11622 | 500 | 0.753 | 0.949 | 0.851 | 0.946 | 0.106 | 0.834 | 0.247 | 0.051 |
| | | Random | 9297 | 2325 | 0.666 | 0.947 | 0.806 | 0.884 | 0.135 | 0.772 | 0.334 | 0.053 |
| | | Lineage | 10519 | 1103 | 0.648 | 0.917 | 0.782 | 0.876 | 0.117 | 0.677 | 0.352 | 0.083 |

**Supplementary Table ST6. Prediction evaluation metrics for the logistic (resistant–sensitive) model.** $n_{train}$ is the number of isolates the model was fit on; $n_{test}$ is the number the metrics were computed on (for PPC, $n_{test}$ is a random subset of $n_{train}$ with a maximum of 500 isolates; for the random and single-lineage sets, a disjoint held-out subset). Resistance is treated as the positive class throughout. Sensitivity and specificity are the recall of resistant and of susceptible isolates, respectively, and bACC is their mean (balanced accuracy). AUC is the area under the ROC curve, Brier the mean squared error of the predicted resistance probability, and $F_1$ the harmonic mean of the precision and recall of the resistant class. VME and ME are the very-major and major error rates: VME is the fraction of resistant isolates reported susceptible ($1 -$ sensitivity), and ME the fraction of susceptible isolates reported resistant ($1 -$ specificity). An "–" indicates a metric that could not be computed as it is undefined when only one class is present in the test set.

| Organism | AMR | Evaluation | $n_{train}$ | $n_{test}$ | RMSE | CRPS | $R^2$ | MAE | EA |
|---|---|---|---|---|---|---|---|---|---|
| *S. pneumoniae* | PEN | PPC | 611 | 123 | 1.798 | 1.024 | 0.406 | 1.447 | 0.333 |
| | | Random | 488 | 123 | 2.029 | 1.154 | -0.037 | 1.692 | 0.236 |
| | | Lineage | 565 | 46 | 4.618 | 3.282 | -18.569 | 4.498 | 0.043 |
| *S. pneumoniae* | TMP | PPC | 606 | 122 | 1.567 | 0.860 | 0.239 | 1.219 | 0.525 |
| | | Random | 484 | 122 | 1.763 | 0.979 | 0.036 | 1.381 | 0.516 |
| | | Lineage | 560 | 46 | 1.422 | 0.785 | 0.095 | 1.026 | 0.717 |
| *M. tuberculosis* | RIF | PPC | 11622 | 500 | 2.148 | 1.479 | 0.478 | 1.803 | 0.286 |
| | | Random | 9297 | 2325 | 2.805 | 1.658 | 0.134 | 2.515 | 0.143 |
| | | Lineage | 10519 | 1103 | 2.554 | 1.541 | -0.139 | 2.358 | 0.064 |

**Supplementary Table ST7. Prediction evaluation metrics for the continuous ($\log_2$ MIC) model.** $n_{train}$ is the number of isolates the model was fit on; $n_{test}$ is the number the metrics were computed on (for PPC, a random subset of the training dataset with a maximum of 500 isolates; for the random and single-lineage sets, a disjoint held-out subset). All error metrics are on the $\log_2$ MIC doubling-dilution scale, comparing the predicted $\hat{y}$ to the observed $y$. RMSE is the root-mean-squared error and MAE the mean absolute error. CRPS is the continuous ranked probability score, which scores the full posterior predictive distribution. $R^2$ is the coefficient of determination. EA is essential agreement, the fraction of samples with $|\hat{y} - y| \leq 1$ (i.e. within one doubling dilution).
